# Transcriptional regulation of the response to water availability in the resurrection plant *Xerophyta elegans*

**DOI:** 10.64898/2026.04.01.716012

**Authors:** Eugene N. K. Kabwe, Michael P. Edwards, Rafe Lyall, Mamosa Ngcala, Stephen A. Schlebusch, Oliver Marketos, Zoran Nikoloski, Robert A. Ingle, Nicola Illing

## Abstract

Vegetative desiccation tolerance (VDT) has evolved independently across vascular plants, but its genetic basis remains poorly understood. Although VDT is associated with expansion of the *ELIP* gene family, the contribution of other lineage-specific expansions is unclear. We assembled genomes for *Xerophyta elegans* and *Xerophyta humilis*, identifying expanded gene families largely involved in chlorophyll metabolism and abscisic acid–mediated stress responses. Using a dense dehydration–rehydration transcriptome series in *X. elegans* seedlings, we reconstructed the regulatory network underlying VDT. Transcription factors from the *ABF*, *ZAT* and *HSFC* families were associated with early responses to desiccation. Key regulators of the seed maturation programme, including NAC transcription factors (*ATAF1* and *ANAC032*), *DOG* genes and the trihelix factor *ASIL1*, were also implicated. These findings indicate that VDT arises through integration of abiotic stress signalling with rewiring of the seed maturation network, enabling desiccation tolerance in vegetative tissues.

## Introduction

Vegetative desiccation tolerance (VDT) - the ability to recover from the loss of >95% of cellular water - was instrumental in allowing plants to colonise land. Although VDT was lost from most lineages following the evolution of vascular systems, it has re-evolved independently in 16 angiosperm genera, collectively known as resurrection plants^1^. Due to the commonalities between VDT and the production of orthodox seeds in angiosperms, including the accumulation of antioxidants, heat shock proteins (HSPs) and late embryogenesis abundant (LEA) proteins, it is thought that VDT re-evolved via co-option of the seed DT network^2^. However, whilst seed DT is regulated by the *LEC1-ABI3-FUS3-LEC2* (LAFL) master transcription factors (TFs)^3^, there is no evidence for involvement of these TFs in VDT^4,5^. The transcriptional regulatory network underpinning VDT thus remains unknown, and its identification is critical for understanding the evolution of this trait.

The Velloziaceae (Pandanales) comprise approximately 270 species across five genera. A basal genus, *Acanthochlamys*, consists of a single VDT species, *Acanthochlamys bracteata*, native to China^6,7^. The genus *Xerophyta*, found in the subtropical regions of sub-Saharan Africa, Madagascar, and the Arabian Peninsula, contains 45 species, all of which exhibit VDT^8^. Besides *Xerophyta elegans*, which is homoiochlorophyllous (Extended Fig. 1), all *Xerophyta* species are poikilochlorophyllous, degrading chlorophyll and dismantling plastids during desiccation^9^.

Gene regulatory network (GRN) reconstruction is a powerful method for identifying regulatory relationships that drive biological processes. To date, only one study has applied this approach to VDT, comparing the lycophyte *Selaginella sellowii* with its desiccation-sensitive sister species *S. silvestris*^4^. Although several angiosperm resurrection plant transcriptomic studies are available^10^, they lack sufficient temporal resolution for GRN inference which requires densely sampled time courses. Moreover, many studies have focused on adult plants, in which asynchronous leaf drying generates spatial heterogeneity in relative water content (RWC). In contrast, *Xerophyta* seedlings, which exhibit VDT immediately post-germination^11^, can be synchronised developmentally, produced in large numbers to enable dense temporal sampling, and - owing to their small size and rapid drying - show reduced water-content heterogeneity compared with adult tissues.

Here we present high-quality genome assemblies for *X. elegans* and *X. humilis*, enabling comparative genomic analyses for these two *Xerophyta* species, with *X. schlechteri*^12^ and the basal *A. bracteata*, in order to identify gene family expansions associated with the evolution of VDT in the Velloziaceae. Using densely sampled transcriptomic data from a dehydration–rehydration cycle in *X. elegans* seedlings, we reconstructed and analysed a GRN, with a focus on the regulation of expanded gene families and a TFs known to be important for seed maturation in Arabidopsis.

## Results and Discussion

### Genome assemblies of *X. elegans* and *X. humilis*

Using PacBio HiFi sequences with hifiasm^13^, we assembled the genomes of *X. elegans* and *X. humilis* (Table 1). K-mer frequency analysis indicated that *X. elegans* and *X. schlechteri* are diploid, while *X. humilis* is tetraploid (Supplementary Fig. 1). The resulting assemblies consisted of 142 contigs with a total length of 414 Mbp for *X. elegans*, and 6,892 contigs with a total length of 1.8 Gbp for *X. humilis* (Table 1), which agreed with their 1C values obtained by FACS analysis (Supplementary Fig.2). BUSCO analysis^14^ indicated that 97.2% and 95.9% of the BUSCO genes were complete in the assemblies of *X. elegans* and *X. humilis*, respectively. Repetitive elements constitute 59% and 65% of the *X. elegans* and *X. humilis* genomes, respectively (Table 1). These proportions are higher than the 18% observed in *X. schlechteri* and *A. bracteata*, partially explaining the larger monoploid genome sizes of *X. elegans* and *X. humilis*. An analysis of syntenic depth in the three assembled *Xerophyta* genomes supports a whole genome triplication event at the base of the *Xerophyta* lineage (Extended Figure 2).

**Table 1.**
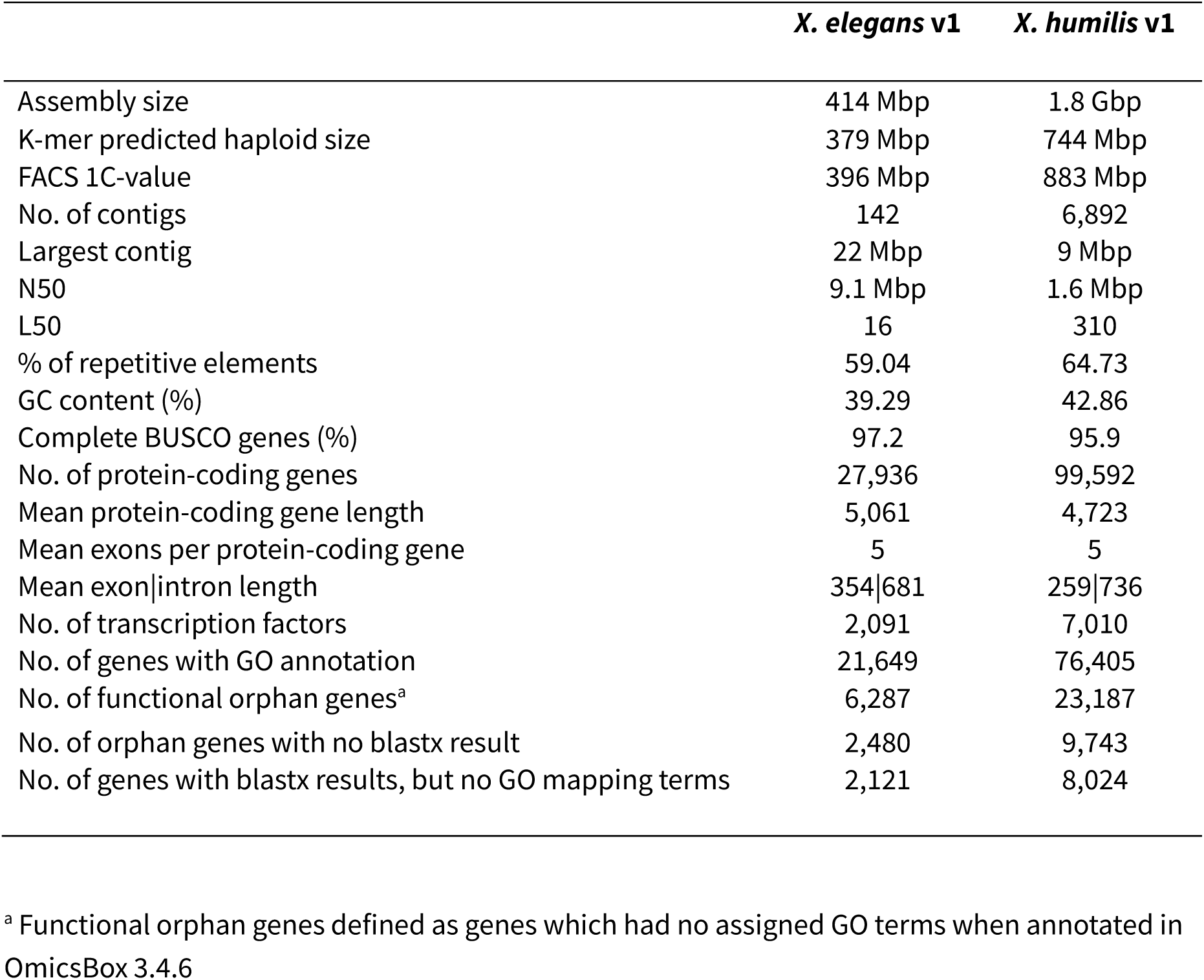
Statistics of the *Xerophyta elegans* and *Xerophyta humilis* genome assemblies and their genome size predictions.

### Gene family expansion in the Velloziaceae

Several comparative genomics studies have attempted to identify convergent features of VDT across all resurrection plants, but only expansion of the *EARLY LIGHT-INDUCED PROTEIN* (*ELIP*) gene family is consistently associated with VDT^15,16^. However, such cross-taxa approaches could hide the role of lineage-specific gene family expansions.

We used OrthoFinder^17^ to identify orthologous gene groups (orthogroups, OGs or gene families) shared between the Velloziaceae and 26 other angiosperm species and found 8,811 containing genes from ≥ 90% of the 30 species. CAFE^18^ analysis identified eleven OGs that were significantly expanded in the Velloziaceae (Fig. 1a). These families include *ELIPs*, *STAY-GREEN PROTEIN* (*SGR*), *C2H2 ZINC FINGER PROTEIN* (*ZAT*), *PYRABACTIN RESISTANCE1/PYR1-LIKE* (*PYL*), *DELAY OF GERMINATION 1 FAMILY* genes (*DOG*), *HEAT STRESS TRANSCRIPTION FACTOR C-type* (*HSFC*), *FCS-LIKE ZINC FINGER PROTEIN* (*FLZ*), and a family of proteins of unknown function that have a PTHR33782 domain (***Ex****panded in **Xe**rophyta* (*EXXE*)). Whilst the expansion of *ELIPs* has previously been linked to VDT in multiple resurrection plant species^16^, the other expanded gene families have not been previously reported as expanded.

**Figure 1.**
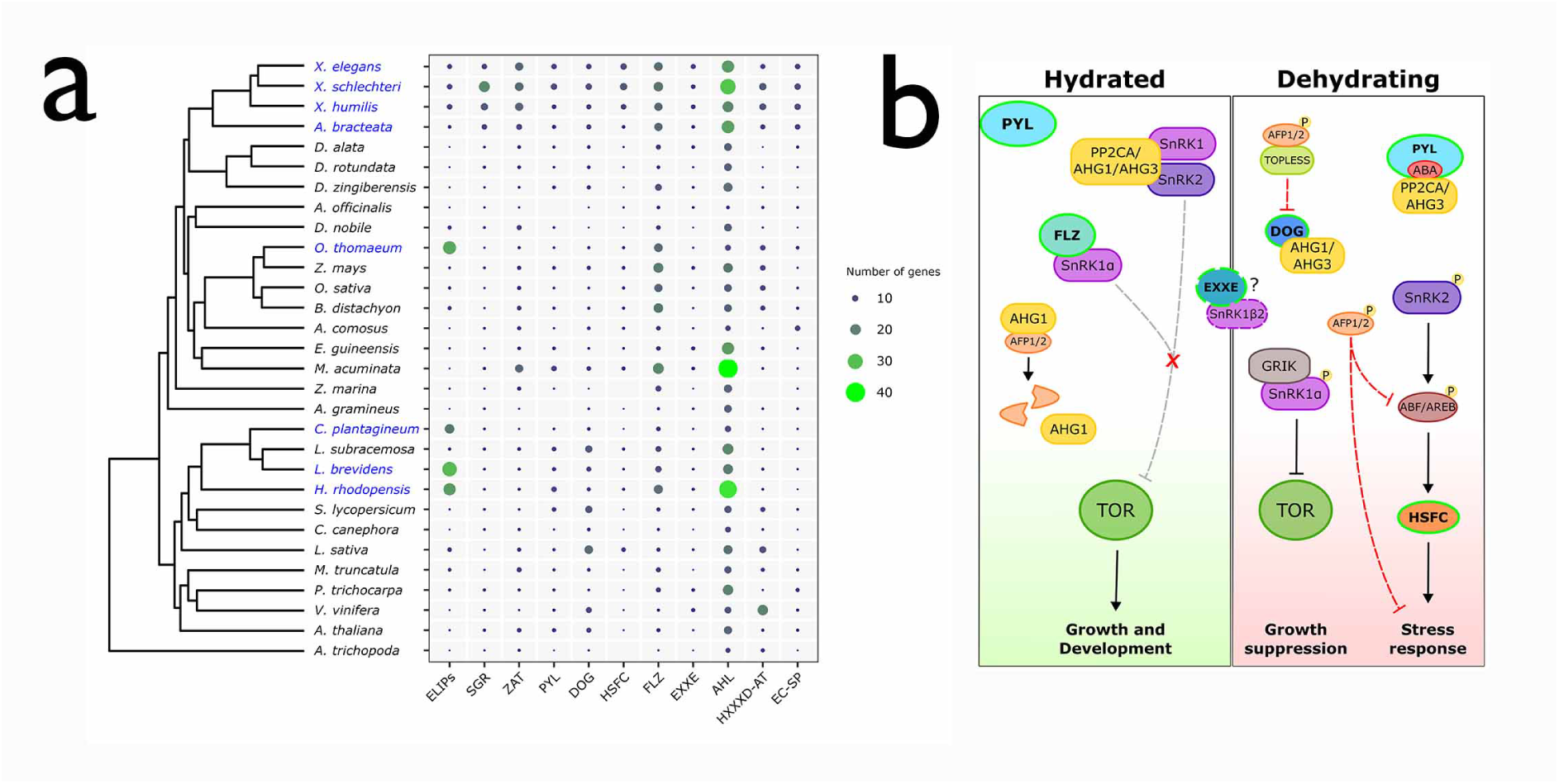
Expanded orthogroups in four Velloziaceae species. **a.** Shown are the number of genes in eleven orthogroups (OGs) that are significantly expanded in four Velloziaceae species compared to 26 other angiosperms, with species names at the base of the phylogenetic tree. Gene counts are normalised for ploidy in polyploid species. The phylogenetic tree was inferred using the STAG method. Species names in blue indicate desiccation tolerant plants and those in black indicate desiccation sensitive plants. The columns are ordered from left-to-right based on the lowest-to-highest number of genes in these orthogroups in *X. elegans*. The eleven expanded orthogroups comprise *EARLY LIGHT-INDUCED PROTEINS* (ELIPs), *STAY-GREEN PROTEIN* (SGR), *C2H2 ZINC FINGER PROTEIN* (ZAT), *ABSCISIC ACID RECEPTOR PYL PROTEIN* (PYL), *DELAY OF GERMINATION 1 PROTEIN* (DOG), *HEAT STRESS TRANSCRIPTION FACTOR C* (HSFC), *FCS-LIKE ZINC FINGER PROTEIN* (FLZ), *EXPANDED IN XEROPHYTA* (EXXE) orthogroup of proteins of unknown function that have a PTHR33782 family domain, *AT-HOOK MOTIF NUCLEAR LOCALIZED PROTEIN* (AHL), *HXXXD-TYPE ACYL-TRANSFERASE* (HXXXD-AT), and *EGG-CELL SECRETED PROTEIN* (EC-SP). **b.** Five of these expanded OGs (highlighted with green borders) are part of the ABA-signaling and growth arrest pathway. Under favourable conditions, Target of Rifampicin (TOR) is responsible for co-ordinating growth, but can be suppressed by Sucrose nonfermenting-related protein kinase 1 (SnRK1) in response to stress. The Group A PP2C phosphatases, including ABA hypersensitive germination (AHG1) and AHG3, keep SnRK1 and SnRK1 related protein kinase 2 (SnRK2) inactive. AHG1 also dephosphorylates the ABI5-binding protein (AFP), rendering it a target for degradation in the proteosome. ABA levels rise rapidly during dehydration stress and seed maturation, and the PYL ABA receptor sequesters the PP2Cs/AHG3 phosphatases. In the absence of FLZ, and the inhibitory PP2Cs, SnRK1 phosphorylates TOR terminating its activity. SnRK2 phosphorylates the ABRE binding transcription factors (AREB/ABFs) which in turn activate expression of Heat Shock Factor HSF-C. This signalling cascade and the stress response is terminated by rising levels of AFP which directly suppress expression of ABF/AREB TFs, as well as DOG, in collaboration with TOPLESS. Dashed lines indicate that the function of the interaction is unknown.

ELIPs are members of the light-harvesting chlorophyll a/b binding (LHC) protein superfamily and serve a photoprotective function^19^. It has been argued that ELIPs have specifically expanded in homoiochlorophyllous DT plants to protect and stabilize photosystem (PS) complexes^16,20^. However, we did not find an association between *ELIP* copy number and chlorophylly strategy; homoiochlorophyllous *X. elegans* contains eight *ELIP*s in its genome, while poikolochlorophyllous *X. schlechteri* and *X. humilis* contain nine (Extended Figure 3, Supplementary Note). The subfamily of *ZAT* TF genes (*C1-2i*) has also been linked to photoprotection in Arabidopsis^21^, and shows an ancient expansion in the Velloziaceae (Supplementary Note). Again, no association between gene copy number of chlorophylly strategy was observed in *Xerophyta*.

In contrast, we did find an association between *SGR* copy number and chlorophylly strategy, in line with the function of SGR family proteins in catalysing the first step of chlorophyll catabolism, triggering PS complex degradation^22^. The expanded *SGR* gene family consists of two subfamilies, the *Type I SGR* family and a *Type II SGR-like* (*SGRL*) family (Extended Fig. 4). We found an initial expansion of the *SGR* genes in *Acanthochlamys* and *Xerophyta*, and a subsequent species-specific expansion of *SGR1* by tandem duplication in *X. humilis* and *X. schlechteri.* Homoiochlorophyllous *X. elegans* has only seven *SGR* genes compared to 20 *SGR* genes in *X. schlechteri* (Extended Fig. 4, Supplementary Note).

Five of the expanded families, *PYL*, *DOG*, *HSFC*, *FLZ*, and *EXXE*, are linked to ABA signalling (Fig. 1b), which is central to the regulation of seed maturation and the vegetative stress response. PYL proteins sequester Group A PP2Cs in the presence of ABA, resulting in activation of sucrose non-fermenting 1-related protein kinase 2 (SnRK2), and subsequent phosphorylation of ABA-responsive TFs, including ABI5 and ABFs/AREBs^23,24^. The seed-specific group A PP2Cs (AGH1 and AHG3), which inactivate SnRK2 by dephosphorylation, are inhibited by the master regulator of seed dormancy, DOG1. SnRK2 would thus be active in the presence of high levels of DOG1, independent of ABA signalling^25^. The expansion of *DOG1* and *DOGL4* to three copies each occurred prior to the diversification of *Xerophyta* species (Extended Fig. 5, Supplementary Note). During drought stress, ABFs activate the heat shock transcription factors (*HSFA6A*, *HSFA6B* and *HSFC1*) in response to ABA signalling ^26^. While Arabidopsis has one *HSFC* orthologue, this family has expanded to eleven copies in *X. elegans* and twelve in *X. schlectheri* (Extended Fig. 6, Supplementary Note).

Increasing stress resilience in plants occurs at the expense of growth^27^. In the absence of ABA signalling, the PPC2:SnRK2 complex binds to SnRK1α, preventing it from suppressing growth^28^. Two of the expanded families (FLZ and EXXE) are linked to the SnRK1 signalling pathway. In Arabidopsis, SnRK1 activity is also suppressed by the binding of FLZ3 to SnRK1α^29^. Members of the *FLZ* family are conserved across all three *Xerophyta* species, and this expansion is likely to have occurred prior to their speciation (Supplementary Note). Members of the *EXXE* family have also been shown to interact with SnRK1, via the SnRK1β2 subunit^30^, but their function remains unclear.

### Identification of desiccation and rehydration responsive genes in *X. elegans* seedlings

To enable GRN reconstruction, we developed a two-leaf-stage *X. elegans* seedling model to profile gene expression across a dehydration–rehydration cycle (Supplementary Fig. 3). Mean seedling survival rate was 97±6.1% (across 64 experimental plates), and rates of water loss and gain were highly consistent (Fig. 2a). Rapid water loss was observed during the first 3 h of drying (100% to 76% RWC) and again after 9 h (>60% to <40% RWC by 12 h; Fig. 2a). Once re-watered, RWC content recovered to 90% within 8 h. The maintenance of chlorophyll content and thylakoid integrity, with the reversible dismantling of grana, are characteristic traits of adult *X. elegans* plants^31,32^ and also occurred in *X. elegans* seedlings (Fig. 2c, d).

**Figure 2.**
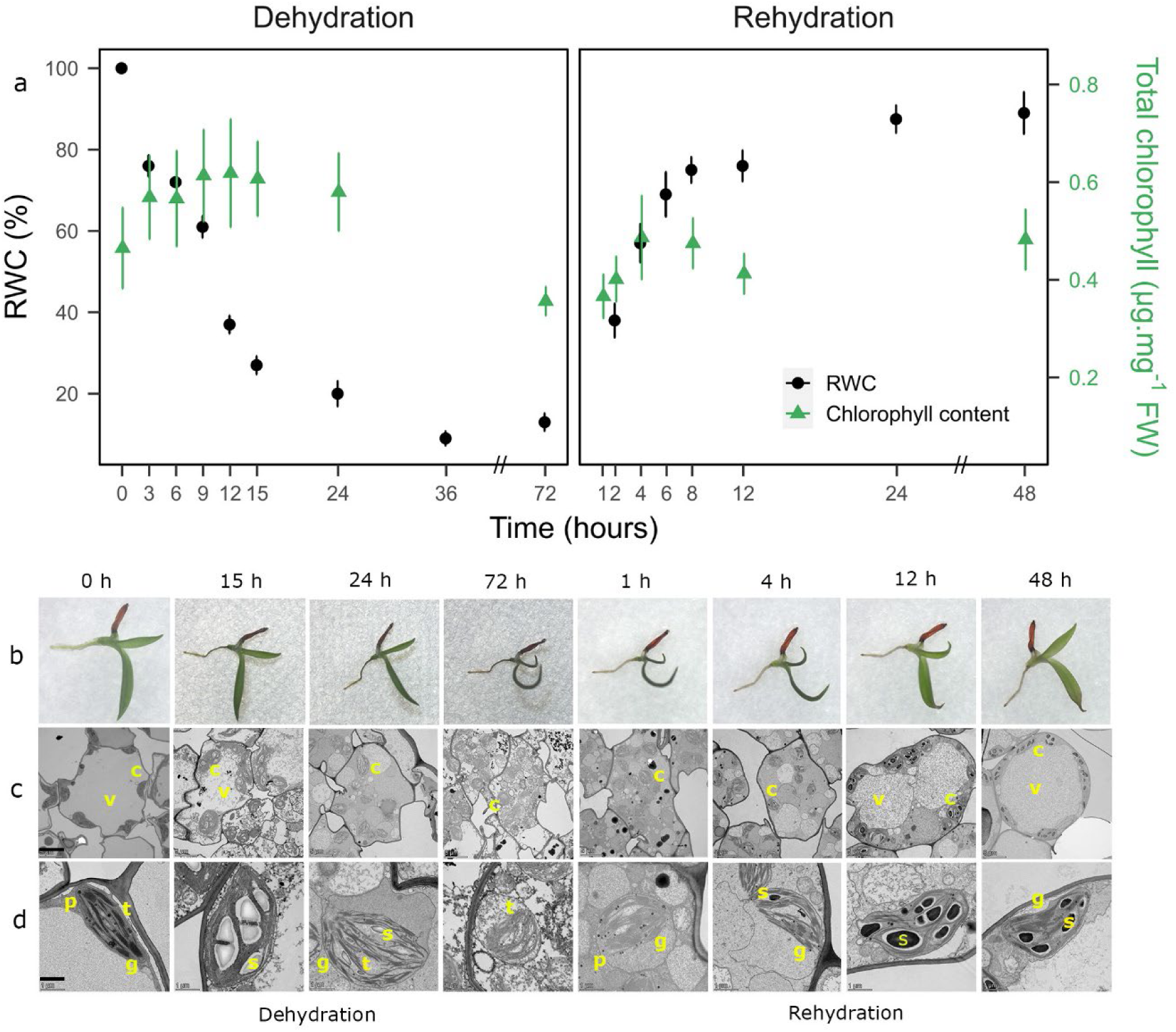
Physiological and morphological changes over a dehydration and rehydration gradient. **a**. Changes in relative water content (RWC, %) (n= 5) and total chlorophyll content (μg/mg FW) (n=7) in two-leaf stage seedlings of *Xerophyta elegans* during a dehydration-rehydration treatment. Wilcoxon pairwise tests between starting (T0) and all other time points identified no significant changes in chlorophyll content at significance level of p<0.05 (Benjamini-Hochbergadjusted). **b**. Light microscope photographs of a dehydrating and rehydrating two leaf-stage seedling of *X. elegans*. Transmission electron micrographs show changes in **c.** chloroplast ultrastructure. Yellow letters indicate vacuole (v), chloroplast (c), starch granule (s), grana (g), thylakoid membrane (s), and plastoglobuli (p). Scale bars indicate 1000 μm (**b**) and 1 μm (**c**).

RNA-seq datasets were generated by sampling seedlings over nine time points during dehydration and six during rehydration (Supplementary Table 1). Principal component analysis separated the samples by RWC on PC1 and rehydrating seedlings from dehydrating seedlings on PC2 (Fig. 3a). The overlap between fully hydrated seedlings (0 h) and seedlings 48 h post-rehydration indicates that, at the transcriptomic level, *X. elegans* seedlings fully recover from dehydration within 48 h of re-watering.

**Figure 3.**
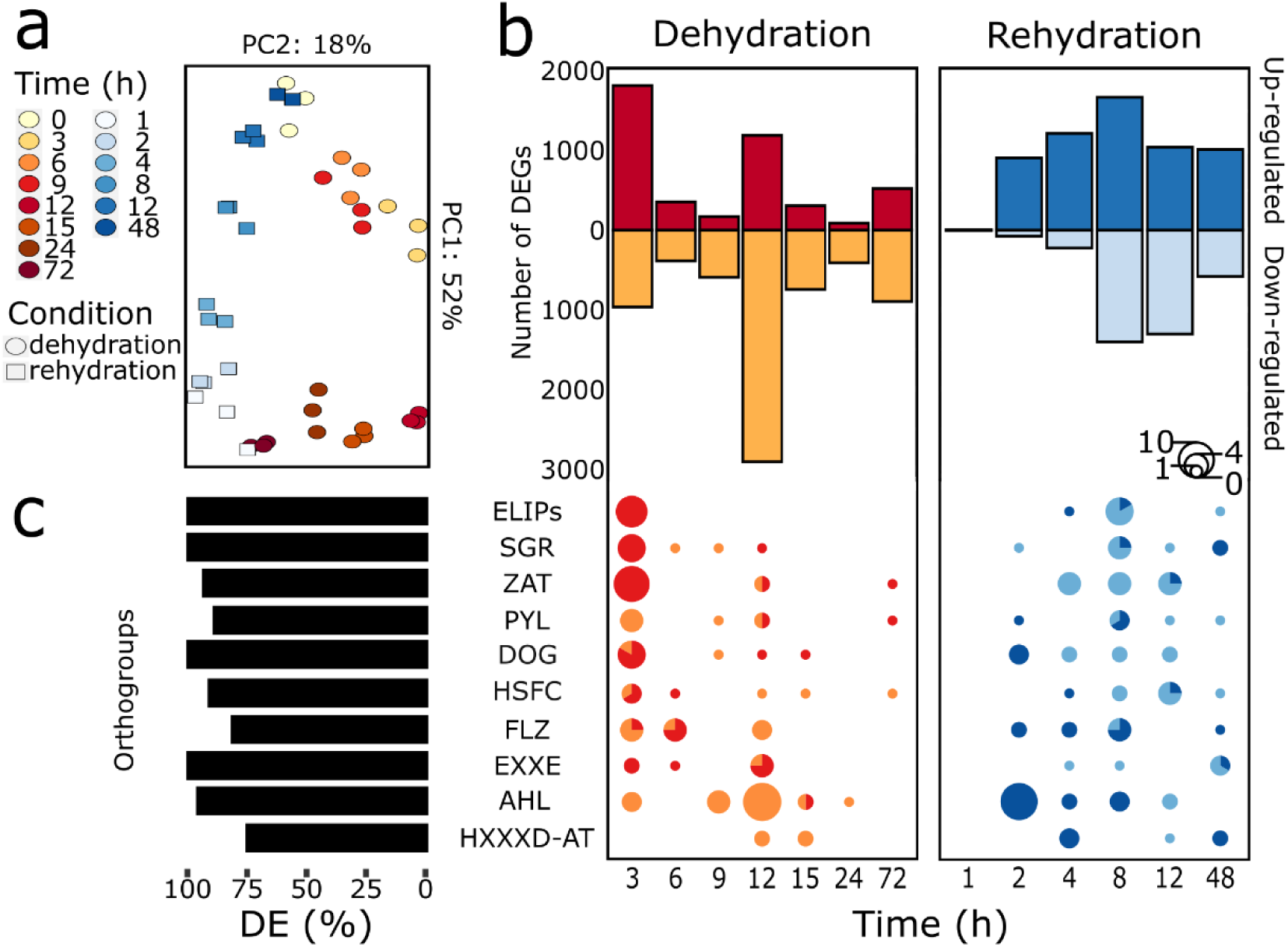
Temporal shifts in the transcriptome of *Xerophyta elegans* seedlings in a dehydration-rehydration cycle. **a**. Principal component analysis of the normalized read counts for the transcriptome (n=3) across the time series in dehydrating (circles) and rehydrating (squares) seedlings. **b.** Earliest time point of diffierential expression during dehydration and rehydration for all diffierentially expressed genes (DEGs) and expanded orthogroup members. Bar plot (upper segment) shows the number of DEGs (Wald test with BH multiple testing, cut-offi set at |log2FC| > 1.5 and FDR < 0.05) that are first diffierentially expressed at each time point (with respect to the reference time points 0 h or 72 h) for each series respectively, and pie-chart plots (lower segment) for DEGs in the expanded orthogroups (OGs) only. Size of the pie chart indicates the number of members which are first diffierentially expressed at a given time point and direction of regulation is indicated by colour. For the dehydration series dark red indicates upregulation while orange downregulation. For the rehydration dark blue indicates upregulation while light blue downregulation. The members of EC-SP OG were not diffierentially expressed and are not included. **c**. Bar plot showing the percentage of total genes in each expanded OG which are diffierentially expressed during the dehydration-rehydration time-series.

We identified diffierentially expressed genes (DEGs) in the dehydration/rehydration series using fully hydrated (0 h) and desiccated (72 h) seedlings as reference points, respectively. To characterize the temporal dynamics of expression, we determined the number of genes identified as DEGs for the first time at each time point during dehydration and rehydration respectively (Fig. 3b). For the dehydration series, the highest number of up-regulated DEGs (1,822) occurred at 3 h, capturing the early response to drying (Fig. 3b). Whilst 965 DEGs were down-regulated at this time point, it is not until 12 h (RWC <40%) that we observed the highest number of down-regulated DEGs (2,916) (Fig. 3b). Although RWC declined to 20% by 24 h of drying, we observed more up-regulated DEGs at 72 h of drying than at either 15 or 24 h, indicating that transcription can occur at low RWC. There was little change in transcript abundance by 1 h post-rehydration, despite the initiation of granal re-assembly in plastids (Fig. 2d). However, by 2 h of rehydration there were 910 up-regulated DEGs. The number of up-regulated DEGs increased as the seedlings rehydrated, peaking at 1,674 after 8 h, whilst that of down-regulated DEGs remained low until 8 h post-rehydration.

The expanded families involved in photoprotection and chlorophyll degradation - the *ELIPS* and most members of the *SGR* and *ZAT* families - were up-regulated within 3 h of dehydration. Families involved in ABA signalling, such *DOG*, *HSFC*, and *EXX*E were up-regulated within 3 hBy contrast, most members of the *PYL* family were down-regulated during early drying, with only two members up-regulated during the latter stages of dehydration. The observed down-regulation of members of the *AHL* family is consistent with their roles in promoting plant growth and developmental processes^33^ (Supplementary Note).

WGCNA^34^ co-expression network analysis of target genes (TGs) resulted in 11 clusters (Supplementary Fig. 4), with an average of ∼960 genes per cluster. Seven of the TG clusters contain members of the expanded families (Fig. 4a). TG Clusters 1, 3, 5 and 7 contain genes down-regulated early on during dehydration (Fig. 4a), distinguished by how rapidly down-regulation occurs and how quickly expression levels recover. TG clusters 2, 4 and 6 consist of genes up-regulated in response to drying (Fig. 4a). TG clusters 2 and 4 exhibit a biphasic expression pattern with an initial peak at 3 h of drying followed by a more prominent peak at 12 h of drying (RWC <40%). After this point expression levels for TG cluster 4 immediately begin to decline while those of TG cluster 2 only decrease at the onset of rehydration. In contrast, genes in TG cluster 6 only increase in expression at low RWC, peaking at the beginning of rehydration. TG clusters 2, 4 and 6 contain members of every expanded gene family except *HXXXD-AT* (Supplementary Table 2). Notably, a majority of the members of the light-response-related *ELIPs* and *SGR* as well as the ABA-signalling related, *DOG* and *EXXE*, gene families were present in these TG clusters. Orthologues of several Arabidopsis seed-specific maturation genes^2^ including *AtEM6*, *1-Cys peroxiredoxin (PER1)*, *DSI-1-VOC* and *oleosins* were grouped into TG2 and TG4 (Supplementary Table 3).

**Figure 4.**
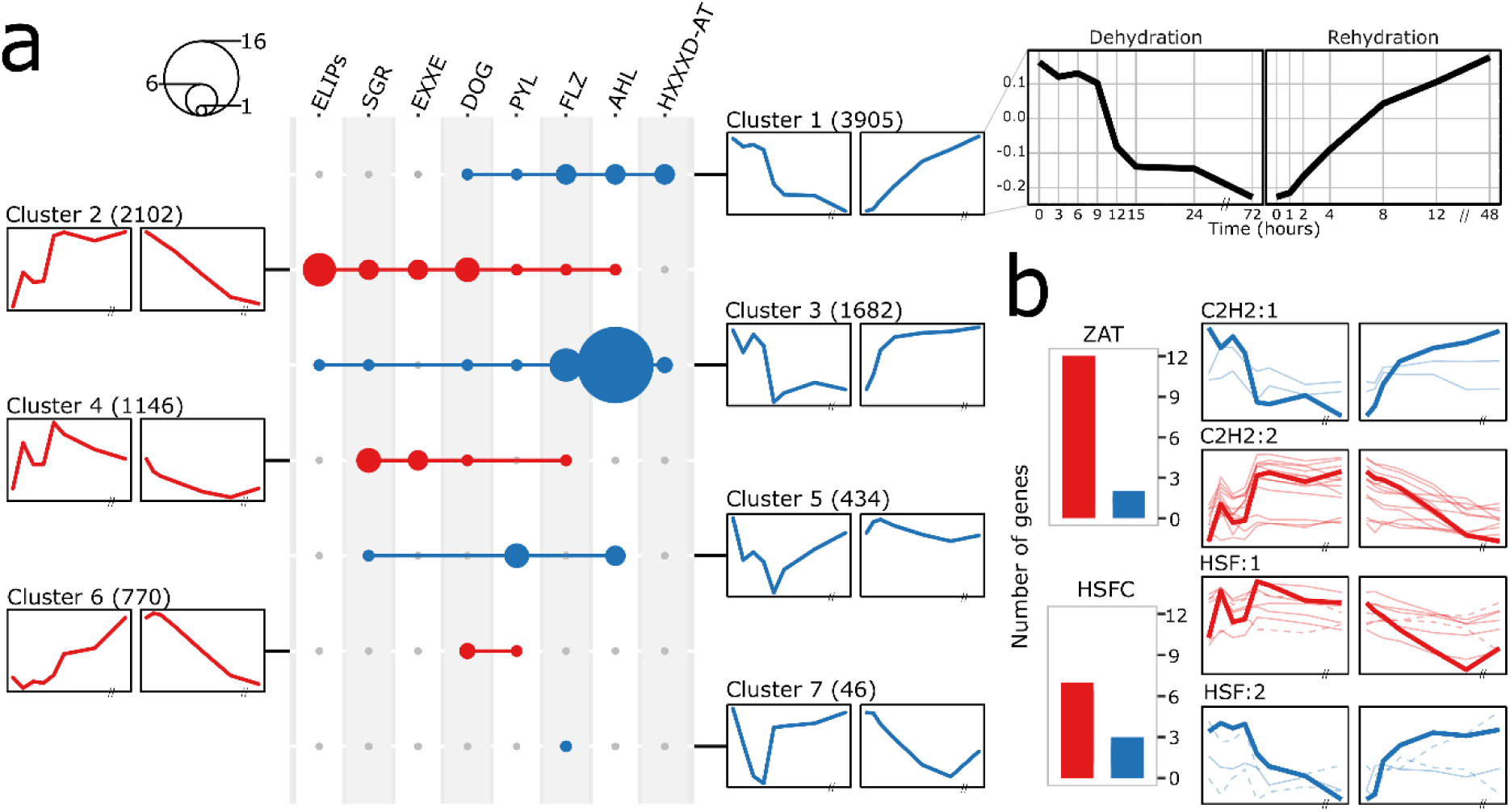
Temporal patterns in orthogroup member gene expression in a dehydration-rehydration series. **a**. DEG expression profiles across the dehydration and rehydration time course were clustered using WGCNA. Members of the expanded orthogroups (OGs) are present in seven of the 11 clusters. The number of expanded orthogroup genes present in these seven clusters is shown in an upset plot. Dot size on the plot represents the number of members of an OG present in a given cluster. Eigengene profiles for each cluster are plotted to either side of the plot and the size of the cluster given in parentheses. A detailed eigengene profile is shown at the top right to illustrate the full time course. **b**. Transcription factors were grouped into families and their expression profiles clustered hierarchically. Bar plots show the number of *ZAT* and *HSFC* expanded orthogroup member genes present in the respective TF family clusters (*C2H2* and *HSF*, respectively). The eigengene profiles for each cluster are shown to the right and overlay the individual member gene expression profiles for that cluster. The colour indicates whether a cluster profile shows increasing (red) or decreasing (blue) expression levels during early and/or late drying. For the HSF profile plots, *HSF-C1* clade members are indicated with a dashed line while *HSF-C2* members are indicated with a solid line.

Of these seven TG clusters (Fig. 4a), TG clusters 1, 2, 3 and 5 are enriched for biological process gene ontology (GO) terms (Supplementary Table 4). Most of the enriched terms for TG clusters 1 and 3 are related to growth and development as well as stimulus/plant hormone responses. This is in line with the presence of the majority of the growth promoting *AHL* family members in cluster 3. TG cluster 2 was primarily enriched for stress response-related terms, including: response to heat, water deprivation, and ABA, and for light response terms including photoprotection, in line with the presence of many *ELIPs* and seed maturation genes in this cluster. Finally, TG cluster 5 has only a few enriched terms, including skotomorphogenesis and chlorophyll biosynthetic process, both processes important for recovery of chloroplasts during rehydration. TG clusters 8, 9, 10 and 11 did not contain members of the expanded gene families but were significantly enriched for GO terms (Supplementary Table 4).

### Consensus-based GRN reconstruction identifies the regulation involving expanded gene families in *X. elegans*

Prior to GRN reconstruction, we clustered the expression profiles of the TF families (Supplementary Table 5). Two of the expanded families are classified as TFs by iTAK^35^: the *ZAT* (*C2H2* family) and *HSFC* (*HSF* family). Most of the diffierentially expressed members of both *HSFC* and *ZAT* belong to clusters up-regulated during drying (‘HSF:1’ and ‘C2H2:2’, Fig. 4b). The *HSF* family members in cluster ‘HSF:1’ are predominantly from the monocot specific HSFC*2* subfamily whereas cluster ‘HSF:2’ contains predominantly *HSFC1* subfamily members (Extended Figure 6, Supplementary Table 2). In wheat, two members of the *HSFC2* subfamily (*TaHSFC2a* and *TaHsfC2d*) are expressed at much higher levels than members of the *HSFA* subfamily during grain filling and maturation^36^.

To identify the TF regulators of and within the expanded gene families during VDT, we reconstructed a GRN using the TF profiles as the predictors of the TG profiles in four diffierent reconstruction approaches; Context Likelihood of Relatedness (CLR)^37^, Elastic Net regression^38^, Gene Network Inference with Ensemble of Trees (GENIE3)^39^ and Network Deconvolution^40^. We integrated the four networks, as a consensus approach has been shown to provide a more robust GRN^41^. The consensus network consisted of 574 TF-TG and 2,906 TF-TF regulatory interactions. The large number of predicted TF-TF interactions is not surprising given that the GRN was constructed using a consensus approach where the dimensions of TFs and TGs were reduced by considering the representative eigengene profiles for the 11 TG clusters (Supplementary Fig. 4) and 58 TF clusters (Supplementary Table 5).

Promoters of the TG clusters are expected to be enriched for the TFBDs of the predicted regulators. To test this hypothesis, we performed an *in silico* validation of the GRN by motif enrichment analysis (Methods). The highest true positive rates (TPR), over the majority of the network, were observed when motif enrichment was performed with proximal promoter regions (Fig. 5b). Enrichment relative to a non-DE set of background gene promoters resulted in the highest TPR values of ∼0.29 when considering the top 10% of the network. These values are in line with other GRNs reconstructed in eukaryotes^41^.

**Figure 5.**
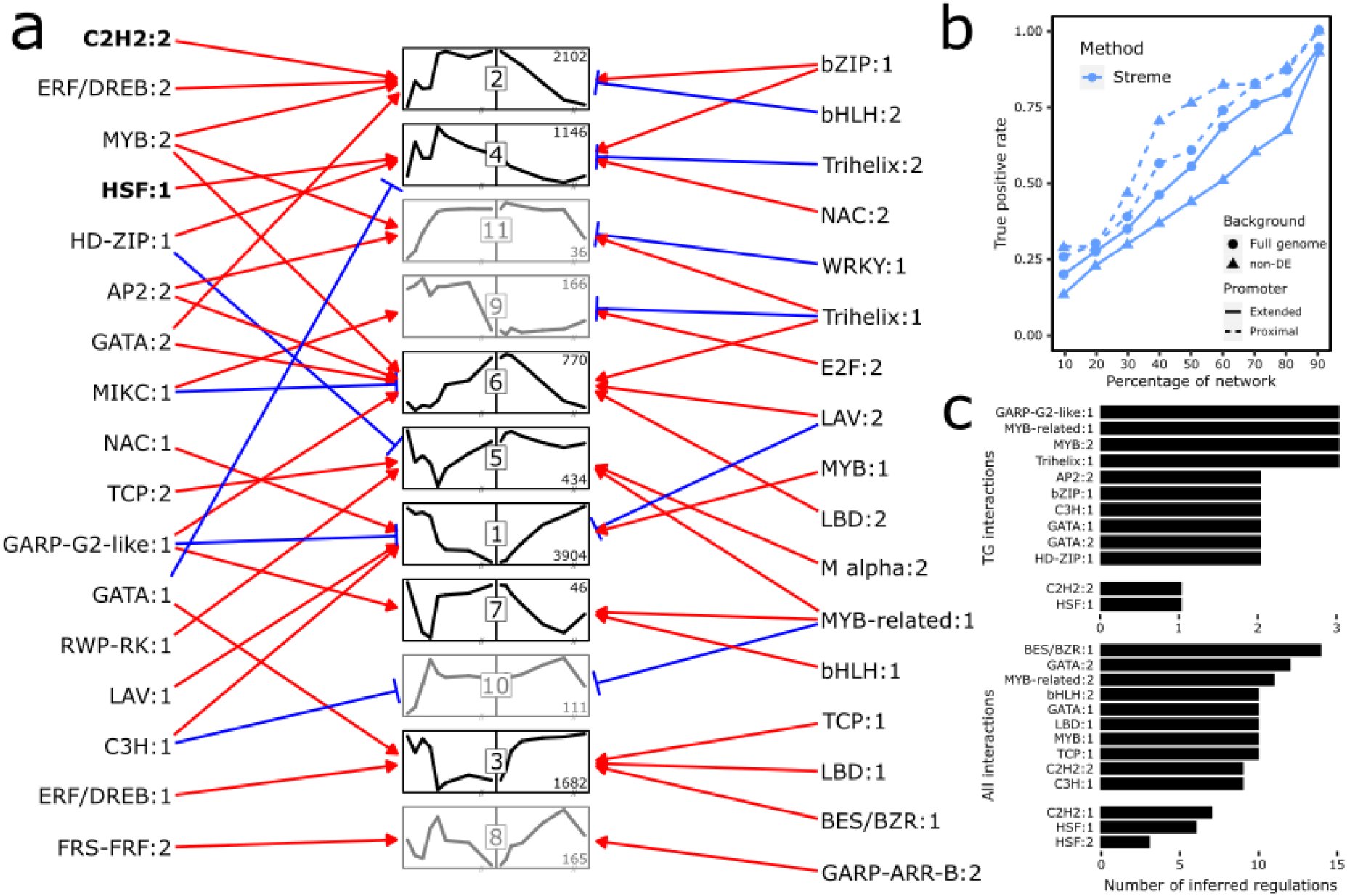
Reconstruction of gene regulatory network using a consensus approach. **a**. Bipartite plot of the transcription factor to target gene (TF-TG) interactions predicted in the top 10% of interaction in the consensus network. TG clusters are designated by their number and their eigengene profile is shown in the inlay panels. TG clusters which do not contain any members of the expanded OGs are indicated with a lighter shade relative to those that contain expanded OG members. TG Cluster size is indicated to the top/bottom right of each profile. TF clusters are designated by both TF family and cluster number and are present to either side of the TG cluster profiles with arrows indicating regulatory relationships. Red arrows denote activation and blue flat-head arrows denote repression. The TF clusters which contain the expanded ZAT and HSF-C OGs are indicated in bold. **b**. Plot of the true positive rate from motif enrichment when used to validate all TF-TG interactions. Dashed lines denote use of the proximal promoter region (500bps upstream of TSS) while the solid line denotes the use of an extended promoter region (2000bps upstream through to 500bps downstream of TSS). The background gene set used is denoted by the shape of the points. **c**. Bar plots showing the top ten TF clusters as ranked by number of TF-TG interactions and all interactions (both TF-TF and TF-TG) they regulate in the top ten percent of the consensus network. Clusters for the *C2H2* and *HSF* families have been included along with the top ten rank regulators.

We ranked TF clusters on the number of predicted TG regulations (Figure 5b). When considering TF-TG interactions, ‘Golden2, authentic response regulators type B (ARR-B), Psr1-Golden2-like:1’ (GARP-G2-like:1), ‘MYB-related:1’, ‘MYB:2’, and ‘Trihelix:1’ were jointly the top ranked TF clusters with each having three inferred regulations. When considering all interactions, ‘BES/BZR:1’ was the highest ranked TF cluster with 14 predicted regulations (Supplementary Fig. 5). Homologues of the Arabidopsis *BEH2* and *BEH4* genes are represented in ‘BES/BZR:1’ (Supplementary Table 5). The *BZR/BES1* family stimulate growth in transgenic Arabidopsis plants and act downstream of brassinosteroid signalling^42^. TG cluster 3 is associated with seedling growth and development which positions ‘BES/BZR:1’ as a potential regulator of growth and development.

We next evaluated the top 10% interactions in the consensus network, including 50 TF-TG, to check for evidence supporting the hypothesis that VDT re-evolved via co-option of transcriptional regulators of the seed maturation network^2^. In agreement with our previous study on *X. humilis*^5^, members of the seed LAFL network were not predicted to be regulators of the desiccation induced TGs in the *X. elegans* GRN. The B3 domain containing LAV family (Supplementary Table 5), includes the LAFL network genes, *ABI3*, *LEC2* and *FUS3*. Of these, *X. elegans ABI3* is included in cluster ‘LAV:2’, however it is expressed at very low levels throughout desiccation, with a minor peak in expression during early rehydration (Extended Figure 7). Similarly, *X. elegans* homologues of the seed maturation TF ABI5 map to ‘bZIP:2’ and are not upregulated during dehydration, though both show upregulation during rehydration (Extended Fig. 7).

Co-expression network analysis has identified two desiccation tolerance transcriptional subnetworks which act downstream of the LAFL master regulators in Arabidopsis^43^. Homologues of four of the identified seed master regulators, *DOGL4,* two NAC domain TFs, *ANAC032* and *ATAF1*, and the ethylene responsive factor, *RAP2-13/ERF58*, were expressed in the *X. elegans* seedlings. Two *X. elegans DOGL4* paralogues are expressed at very high levels in in the late stages of desiccation (Extended Fig. 8b). DOGL4 is a major inducer of reserve accumulation during seed maturation in Arabidopsis^44^. Unlike DOG1, DOGL4 is induced by ABA, and directly activates the expression of seed maturation-specific genes^44^. Both NAC TFs, *ANAC032* and *ATAF1*, showed the typical biphasic activation of expression as seedlings desiccate (Extended Fig. 8 e,f). In Arabidopsis, ANAC032 activates chlorophyll degradation, suppresses photosynthesis and triggers the carbon starvation response^45,46^. Arabidopsis TREHALASE I (TRE1) and NINE-CIS-EPOXYCAROTENOID DEHYDROGENASE (NCED3) enzymes are direct targets of ATAF1^47^. Homologues of these enzymes show a similar biphasic expression to *ATAF1* in *X. elegans* (Extended Fig. 8 g,h). In Arabidopsis, *ERF58* is specifically expressed in mature seeds and controls the completion of seed germination downstream of the phytochromes *phyA* and *phyB* ^48^. The *RAP2-13/ERF58* TF is included in the ‘ERF-DREB:2’ cluster, though it is expressed at relatively low levels throughout the dehydration-rehydration cycle (Extended Fig. 8a).

The most abundant transcript in ‘Trihelix:1’ is the *6b-INTERACTING PROTEIN-LIKE1* (*ASIL1*) TF, well known as a key integrator within the seed maturation network^49^, acting as a temporal regulator during seed filling and repressing the seed maturation programme during germination^50^. *ASIL1* expression increases during the late phases of desiccation and peaks within two hours of rehydration in *X. elegans* seedlings (Extended Fig. 8c), consistent with its proposed role of suppressing genes involved in the seed maturation programme^49^. *DE1 BINDING FACTOR* (*DF1*) is expressed at the highest level in ‘Trihelix:2’, and is up-regulated transiently during dehydration, and again during rehydration (Extended Fig. 8d*). DF1* regulates expression of pectin during seed maturation^51^.

The early drying-responsive TG clusters 2 and 4 are both predicted to be activated by ‘bZIP:1’ in the GRN (Fig. 5a). *XhABFA*, a member of the *ABA-RESPONSIVE ELEMENT BINDING FACTORS* (*ABF*) bZIP subfamily, and homologue of *Arabidopsis ABF2/AREB1,* has previously been shown to drive transcription of *ABI3* regulon homologues in *X. humilis*^5^. *X. elegans ABF* homologues are found within ‘bZIP:1’, with *ABF2*/*AREB1* and *AREB2* being expressed at the highest levels (Extended Fig. 9, supporting a role as candidate key regulators of desiccation-responsive target genes in *Xerophyta* species. Genes in the TG cluster 2 and 4 are also regulated by members of the expanded *ZAT* and *HSFC* families with ‘C2H2:2’ activating cluster 2 and ‘HSF:1’ activating cluster 4 (Fig. 5a).

Overall, the GRN points at the importance of homologues of the ABF2/AREB1 and AREB2 transcription factors (in the ‘bZIP:1’ TF cluster), and TFs in the expanded ‘C2H2:2’ ZAT and ‘HSF:1’ clusters, as activators of the target gene clusters, TG2 and TG4 which respond early to desiccation (Extended Fig. 10). Although the canonical seed maturation TFs, *ABI3* and *ABI5* are not expressed in *X. elegans* seedlings as they desiccate, other seed TFs are. These include *ATAF1* and *ANAC032* which are activated early during desiccation, *ERF58* and *DOGL4* which are activated in later stages, and the Trihelix repressor *ASIL1* which is active in the earliest stages of rehydration (Extended Fig. 10).

## Conclusion

Using a comparative genomics approach, we report the identification of expanded gene families in *Xerophyta* associated with VDT. Using a seedling model, we generated a fine scale transcriptomics time course to dissect TF-target gene interactions during VDT through GRN reconstruction. Expanded gene families related to light response (*ELIPs* and *SGRs*) and ABA-signalling (*EXXE* and *DOG*) were up-regulated as part of the response to drying. The regulation of the response to desiccation and rehydration in *X. elegans* has likely evolved from integration of TFs from both the abiotic stress response as well as TFs active during seed maturation and germination.

## Materials and Methods

### Plant and seed collection

*X. elegans* adult plants were collected from the banks of a stream near Monk’s Cowl in the Maloti-Drakensberg Park World Heritage Site, KwaZulu-Natal, South Africa (Extended Fig 1) (Lat/Lon: −29.05399, 29.40145), and grown under glass house conditions at the University of Cape Town. Mature seeds pods were harvested from the same population at the Monk’s Cowl site (Extended Fig 1). The seed pods were allowed to dry out at ambient temperature, and seeds were collected into paper bags by gently shaking the seed pods. The seeds were stored on desiccant at ambient temperature until needed. These collections were approved by Ezemvelo/KZN Wildlife permit number OP 3925/2019.

### Seedling sterilisation and germination

Sterilisation of the seedlings was performed prior to germination. Seeds were visually inspected and sorted into batches of 100. Sterilisation began with heat treatment for an hour at 65 °C. The seeds were then treated sequentially with the following solutions with vortexing: 100% ethanol for 10 seconds, 70% ethanol solution with 1µl/ml Triton-X100 for 20 seconds and then a 15% H2O2 solution with 1µl/ml Triton-X100 for a minute. After these treatments, the seeds were washed three times by inversion with dH2O. The sterilised seed batches were transferred onto sterile filter paper to dry. Each batch was plated onto a half strength Murashige and Skoog (MS) medium agar plate, parafilmed and transferred to a plant growth chamber, maintained at 22 °C and with a 16/8h day/night cycle, to germinate. The seedlings reached the two-leaf stage of development within ∼2 weeks.

### Seedling Drying and rehydration protocol

To perform reproducible drying and rehydration the seedlings were kept within the plant growth chamber. The seedlings dried/rehydrated on filter paper within an open face Petri dish placed within a transparent box to slow the rate of drying (Supplementary Fig 6). Each sample consisted of 20 seedlings. Prior to the experiment, the seedlings were plated onto filter paper with 2ml of dH2O and transferred back to the plant growth chamber overnight. This helped the seedlings reach turgor and improved seedling survival rate. The morning of an experiment, the pre-selected seedlings were transferred to 2ml Eppendorf tubes and weighed to determine fresh weight. The seedlings were transferred to fresh filter paper with 1ml dH2O on a plate. The boxes were placed within the growth chamber at the start of the day cycle and the plates uncovered to allow drying to commence. For rehydration experiments the seedlings were allowed to dry for 3 days until air dry. A desiccated sample was then taken (DeT72). To begin rehydration, 2ml dH2O was added to each plate, which were sealed with parafilm to prevent water loss. The boxes were then placed back in the plant growth chamber at the beginning of the day cycle.

To demonstrate the reproducibility of this drying/rehydration protocol, the relative water content (RWC) of the seedlings was determined over a series of five experiments. The sets of 20 seedlings were weighed at their respective time point. Dry weight was determined following oven drying of the sample at 60 °C for 48 hours. The following formula was used to determine the percentage RWC:

RWC (%) = [Fresh weight (FW) - Dry weight (DW)] / [Turgid weight (TW) - Dry weight (DW)] x 100

Along with RWC, the survival of seedlings following complete desiccation was assessed. All experiments included a recovery plate where after five days of rehydrating the seedlings were examined for damage to one or both leaves and the presence of new growth.

A plate with 20 seedlings for the measurement of chlorophyll content and ultrastructural changes were also included in the experimental design.

### Measurement of total chlorophyll content

A sample size of five seedlings was used for each chlorophyll measurement. Seedlings were placed into 200µl acetone (100%) and homogenised. The solution was incubated at 4°C in the dark for 48 hours. The absorbance at 662nm and 645nm were measured with a spectrophotometer and total chlorophyll content determined with the following formula: (7.05 x A662) + (18.09 x A645)^52^.

### Ultrastructure section

*X. elegans* two-leaf seedlings were processed for transmission electron microscopy (TEM) ultrastructure using a protocol described by Farrant et al. (1999), with minor modifications^53^. Briefly, dehydrating and rehydrating seedlings were fixed in 2.5% glutaraldehyde in 0.1 M phosphate buffier (pH 7.4) containing 0.5% caffieine, followed by washing and performing the second fixation in 2% osmium tetraoxide. Samples were dehydrated in a graded ethanol series at room temperature prior to infiltrating with epoxy resin^54^. The final step involved embedding samples in pure Spurr’s resin and hardening in a 60 °C oven for at least 16 h. Subsequently, 90 nm thick, ultrathin sections were produced from the sample resin blocks before staining them with 2% uranyl acetate and 1% lead citrate. Stained sections were examined using TEM at both low (2000x) and high (11 000x) magnifications. The examination covered the general cell structure at low magnification and individual chloroplasts in various leaf sections at high magnification. A total of 25 images were taken in each biological repeat at every time point, comprising 5 images depicting the overall cell structure and 20 images displaying individual chloroplasts from diffierent leaf regions.

### RNA extraction and sequencing

For RNA-seq at each time point the 20-seedling sample was flash frozen in liquid nitrogen and transferred directly to a −80 °C freezer for storage until extraction.

Total RNA was extracted with the Direct-Zol RNA miniprep kit (Zymo research) but with an initial extraction protocol employing a LiCL extraction solution and PCI treatment. Frozen seedlings samples were homogenised using a Retsch mill. The pulverised samples were resuspended in 400µl of cooled extraction buffier (150mM LiCl, 50mM Tris-HCL, 5mM EDTA and 1% SDS in DEPC-treated water) with vortexing for 2 minutes. 400µl of PCI (pH 4) was added and the sample vortexed again for 30 seconds. Two rounds of centrifugation each for 12 minutes at 12000g and 4 °C were carried out with the further addition of 400µl of chloroform prior to the second round. The aqueous layer was transferred to a fresh 2ml tube, vortexed briefly and incubated for 5 minutes at room temperature. A further 200µl of chloroform was added at which point the prior vortexing and incubation steps were repeated. Trizol reagent was then added to the sample and the Direct-Zol kit protocol performed with the following notes: a 15-minute DNAse I treatment was carried out prior to elution and an additional wash was added at each wash step in the protocol. RNA concentration and integrity were assessed using a Nanodrop1000 (Thermo) and by gel electrophoresis employing a denaturing agarose gel. For the latter, well defined and bright 18S and 28S bands were used as markers for intact RNA. Samples were stored at −80 °C. For shipment to Beijing Genomic Institute (BGI) at ambient temperature, RNA samples were added to GenTegraRNA matrix and dried for 4 hours with a Savant Speedivac plus system

High-quality RNA was also isolated from *X. elegans* adult leaves, roots, seeds and seedlings using the Zymo Direct-zol RNA Miniprep kit and directly sent to Inqaba biotec for Iso-seq sequencing on the PacBio Sequel II system. The PacBio “extracthifi” script (https://github.com/PacificBiosciences/extracthifi) was used to extract HiFi reads. LIMA was used to remove incomplete reads and primer sequences. The IsoSeq3-refine tool was then used to remove redundant reads, non-full-length reads and trim poly-A tails. Lastly, the IsoSeq3-cluster tool was used to cluster full-length non-redundant reads into a final set of polished, full-length isoforms.

### Nuclear DNA content in Xerophyta plants

Flow cytometry was employed to estimate the genome size of *Xerophyta* species, using a method adapted from Dolezel et al. (2007)^55^. *Raphanus sativus* L. ‘Saxa’ (1.11 pg) and *Solanum lycopersicum* (1.96 pg) were used as reference standards^56^. Fresh leaves (400-600 mg) were harvested on the experiment day from the plant growth room and finely chopped with a sterile razor blade in 1 mL of ice-cold Otto I solution [0.1 M citric acid, 0.5% (v/v) cell culture Tween 20] in a plastic Petri dish. The homogenate was filtered through a Falcon 40 μm nylon cell strainer, and isolated nuclei were pelleted by centrifugation at 100*x g* for 6 minutes. The resulting supernatant was removed, leaving approximately 100 μL liquid above the pellet. An equal volume of fresh ice-cold Otto I solution was added, and the nuclear suspension was stored at 4 °C. A four times volume of Otto II [0.2 M Na_2_HPO_4_.12H_2_O] solution, along with 50 μg ml^-1^ RNase A and 50 μg mL^-1^ of propidium iodide was added to the nuclear solution, immediately prior to the FACS analysis. The fluorescent signal of the stained nuclear suspension was measured on a flow cytometer (BD LRFortessa^TM^, Becton, Dickinson and Company, USA). The nuclear DNA content of *Xerophyta* plants was calculated using the following formula:

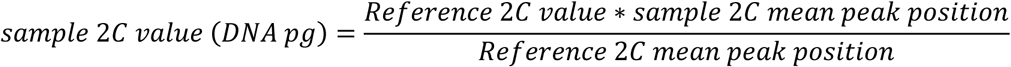

The DNA mass was converted from picograms to the number of base pairs as follows:

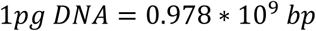

### DNA extraction and sequencing

High molecular weight genomic DNA was extracted from 1.5 g of fresh adult *X. elegans* or *X. humilis* leaf tissue. Briefly, fresh leaf tissue was ground in liquid nitrogen followed by lysis using a buffier consisting of 100 mM Tris, 50 mM EDTA, 500 mM NaCl, 1.25% SDS, 2% PVP-40 and sodium bisulphite. The DNA was then precipitated in isopropanol and resuspended in TE buffier before it was eventually purified by centrifugation with Phenol-Chloroform-Isoamyalchohol (25:24:1) followed by chroloform:isoamyl alcohol (24:1). The purified genomic DNA was sequenced using PacBio circular consensus sequencing (CCS) on the PacBio Sequel II system. The sequencing of *X. elegans* was done at the Georgia Genomics and Bioinformatics Core (GGBC) at the University of Georgia, USA, and the sequencing of *X. humilis* was done at Inqaba Biotec in Pretoria, South Africa. SMRT Link was used to generate long and high-fidelity (HiFi) reads from the sequencing. The quality of the reads was assessed using the LongQC tool^57^. The sequencing of both genomes using PacBio HiFi technology yielded 29x coverage for *X. elegans*, and 56x coverage for *X. humilis*.

### Genome assembly and completeness assessment

To assemble the *X. elegans* genome, we used the Hifiasm assembler^13^ using default parameters for a HiFi-only assembly. One round of duplicate trimming of the assembly was done using purge_dups^58^. To assemble the *X. humilis* genome, we used similar parameters as for *X. elegans*, but the homozygous coverage was set to 16, the expected haplotype number was set to 8 and all types of haplotigs were purged with a similarity threshold of 0.95. No duplicate trimming was performed for the *X. humilis* assembly. Lastly, the primary and alternate assembly were merged to create a complete assembly consisting of all 4 haplotypes. Organelle genomes were aligned to the assemblies using Minimap2^59^ to identify potential organelle genome sequences, which were subsequently removed. Completeness of the assemblies based on the embryophyte lineage was assessed using BUSCO v5.2.2^14^.

### Genome annotation

Repeat sequences were identified using the EDTA pipeline^60^. The *de novo* transposable element library identified by RepeatModeler was combined with the non-redundant TE library (version 19) of the TRansposable Elements Platform (TREP)^61^ and was used to softmask the genome using RepeatMasker. Protein-coding gene prediction for *X. elegans* was performed by first using the PASA pipeline^62^ to generate full-length transcripts from the *X. elegans* PacBio Iso-seq data. The short-read RNA-seq reads were then mapped to the genome using STAR v2.7.10a^63^, to use with BRAKER2^64^ as RNA evidence. The exonerate v2.2.0 tool^65^ was used for gene prediction based on protein homology from the protein sequences of *Arabidopsis thaliana*, *Triticum aestivum*, *Oryza sativa* and *Zea mays*. Lastly, EVidenceModeler v1.1.1^66^ was used to combine the diffierent prediction methods and create a non-redundant high-confidence gene list. The protein-coding gene prediction for *X. humilis*, *X. schlechteri* and *A. bracteata* were performed using the same methodology with the exception of the use of PacBio Iso-Seq data. Functional annotation of the genes was performed using Blast2GO with the target database restricted to *Arabidopsis thaliana*. Transcription factor prediction was done using iTak^35^.

### Comparative genomics

We used OrthoFinder v2.5.5^17^ with the parameters -I 1.25, -S diamond, -M msa and -A mafft to generate orthogroups for comparative genomics. The comparison included 30 angiosperm species namely *Xerophyta elegans*, *Xerophyta schlechteri*, *Xerophyta humilis*, *Acanthochlamys bracteata*, *Dioscorea alata*, *Dioscorea rotundata*, *Dioscorea zingiberensis*, *Asparagus officinalis*, *Dendrobium nobile*, *Oropetium thomaeum*, *Zea mays*, *Oryza sativa*, *Brachypodium distachyon*, *Ananas comosus*, *Elaeis guineensis*, *Musa acuminata*, *Zostera marina*, *Acorus gramineus*, *Craterostigma plantagineum*, *Lindernia subracemosa*, *Lindernia brevidens*, *Haberlea rhodopensis*, *Solanum lycopersicum*, *Coffiea canephora*, *Lactuca sativa*, *Medicago truncatula*, *Populus trichocarpa*, *Vitis vinifera*, *Arabidopsis thaliana* and *Amborella trichopoda*. After the orthogroups were generated, the polyploid genomes’ numbers were normalised by dividing the number of identified genes by 4 for the tetraploid *X. humilis* and by 8 for the octoploid *C. plantagineum.* Integers were then rounded up to the nearest whole number. Orthogroups with at least 90% of the species present were used with the rooted phylogenetic tree, generated using the STRIDE method, as input for CAFE v5.0.0^18^ to identify the significantly expanded orthogroups in the Velloziaceae with a p-value cutoffi of 0.05.

### RNA-seq library construction, quality control and alignment

RNA quality was assessed upon arrival at BGI using bioanalyser (Agilent technologies 2100 bioanalyser system). The DNBSeq platform was employed for strand-specific mRNA library preparation employing poly-A tail selection. Initial data filtering of raw reads was performed by BGI using the SOAPnuke package^67^. Quality of the clean reads was assessed using fastqc^68^. The 42 RNA-seq libraries produced a total of 860 million paired reads which were aligned to the *X. elegans* genome assembly to determine transcript level abundance Alignment of the reads to the *X. elegans* genome assembly was accomplished with STAR^63^. In accordance with the size of the genome assembly the option genomeSAindexNbases was set to 13. To obtain gene level abundances, featureCounts was used with the *X. elegans* genome annotation while excluding multi-mapping reads^69^.

### Diffierential gene expression analysis

The R package DESeq2 was employed for diffierential gene expression analysis^70^. DEGs were identified by pairwise analysis with the Wald test for each series (dehydrating and rehydrating) separately. For the dehydrating series diffierential expression (DE) was determined relative to fresh seedlings (DeT00); for the rehydrating series, DE was determined relative to desiccated seedlings (DeT72). A gene was considered DE if the absolute value of its log-fold-change was larger than 1.5 and the Benjamin-Hochberg (BH) adjusted p-value was smaller than 0.05. To improve the estimated log-fold-change (LFC) for genes with low or variable levels of expression, LFC shrinkage was carried out with the ashr package^71^. Temporal patterns of DE were determined to identify the earliest time point at which a gene was found to be diffierentially expressed. GO enrichment was determined using the Fisher exact test with the weight01 algorithm of topGO^72^. These results were summarised with the viSEAGO (Wang’s method derived semantic similarity distance and ward.D2 aggregation criterion) packages^73^.

### Gene regulatory network reconstruction

All DEGs were classified as either transcription factors (TFs) or target genes (TGs) by using the iTAK identified TF list. Clustering of the TGs based on their expression profiles was performed with the WGCNA package^34^. The EdgeR package was used to obtain normalised counts per million (cpm)^74^. Hierarchical clustering using Pearson correlation was performed with the ‘signed hybrid’ network type option. Dynamic tree cutting was carried out with a mergeCutHeight of 0.2 followed by partitioning around the medoids for unassigned genes. The biological homogeneity index (BHI) was determined for the set of clusters using the clValid R package^75^. A soft thresholding power of 13 was chosen to maximize the BHI value, of 0.355, at a reasonable scale free topology model fit and mean network connectivity. The expression profiles of each TF family were then hierarchically clustered. The optimal number of clusters for each family was determined by the silhouette index.

GRN reconstruction was performed with the TG and TF cluster eigengenes, the TF cluster eigengenes serving as predictors of the TG cluster eigengenes. The GRN reconstruction was carried out as described in Arend et al. (2022)^76^. Opting for a wisdom of the crowd strategy, a consensus network was obtained by applying the following five approaches: Context Likelihood of Relatedness (CLR)^37^, Elastic Net regression^38^, Gene Network Inference with Ensemble of Trees (GENIE3)^39^ and Network Deconvolution^40^. Inferring the consensus network was accomplished using the Borda count election method^76^. The number of edges from each network to consider for the consensus network was determined by the smallest number of edges from each of the networks used to generate the consensus network. The top 10% of all network edges were used for network visualisation and further analyses.

The interactions of the consensus network were evaluated through analysis of the promoter regions of the TG clusters based on enrichment of the cis-regulatory elements of the TF family predicted as a regulator. Promoter analysis was carried out with programmes of the MEME suite [Streme^77^, Tomtom^78^ and SEA^79^] and Homer package^80^. To this end, we used the JASPAR 2024 core plant collection of transcription factor binding motifs (TFBMs)^81^. Simple Enrichment Analysis (SEA) was employed for motif enrichment directly with the TFBMs. De novo identification of enriched motifs using the Streme and Homer was also carried out. These enriched motifs were then aligned to the JASPAR core plant TFBMs using the Tomtom motif alignment tool. For all approaches both a proximal (500bps immediately upstream of the TSS) and extended (2000bps upstream through to 500bps downstream of the TSS) promoter region was considered. Additionally, two separate backgrounds against which motif enrichment was determined were also considered. A background consisting of all gene promoter regions (excluding those of the TG cluster itself) and another using only promoter regions from non-diffierentially expressed genes. Confusion matrices were generated for each approach to retrieve the sensitivity and specificity of the predictions.

### Databases

The genomic DNA raw reads of *X. elegans* and *X. humilis*, as well as the *X. elegans* time course RNA-seq dataset has been uploaded to NCBI as part of BioProject PRJNA1242325. The genome assemblies and annotations generated in this study have been upload onto the CoGe database (https://genomevolution.org/coge/) under ID: 69056 for *X. humilis* and ID: 69057 for *X. elegans*.

To enable interactive exploration of the transcriptomic data in this study, we developed the *Xerophyta* Data Explorer (https://xerophyta-data-explorer.streamlit.app), a web application for querying and visualizing gene expression across the time series datasets. Users can retrieve genes by *X. elegans* gene ID, *Arabidopsis thaliana* orthologue locus ID, Gene Ontology term, enzyme code, or protein domain, and explore inferred gene regulatory networks (See Supplementary Tables S2, S3, S5). Expression profiles and underlying data are available for download. Source code is archived at Zenodo (DOI: 10.5281/zenodo.16678827).

## Acknowledgements

This work was funded by the European Union’s Horizon 2020 Research and Innovation programme (H2020-MSCA-RISE-2018) under the project acronym RESIST (grant no. 823746), as well as the European Regional Development Fund through Programme Research Innovation and Digitalisation for Smart Transformation, Grant agreement № BG16RFPR002-1.014-0003-C01. We thank Inqaba Biotec^TM^ for providing the resources to sequence the genome of *X. humilis* under the Africa Genome Challenge 2021 project, and Robert VanBuren (Michigan State University, USA) for the sequencing of the *X. elegan*s genome. Jonathan de Wet is thanked for giving us permission to collect *X. elegans* seeds from the stream on his property in the Drakensberg. We thank Kelly Shepherd (University of Cape Town) for assisting with the DNA extraction for *X. humilis*. We are grateful to Mohamed Jaffier and Keren Cooper (both University of Cape Town) as well as Otto Baumann (University of Potsdam) for their guidance with the TEM analyses. Computations in this work were performed using facilities and computing resources provided by the University of Cape Town’s ICTS High Performance Computing team: hpc.uct.ac.za, as well as facilities provided by ZIM, Universität Potsdam. E.N.K.K. and M.N. were funded by the National Research Foundation (NRF) of South Africa.

---

**Extended Figure 1:**
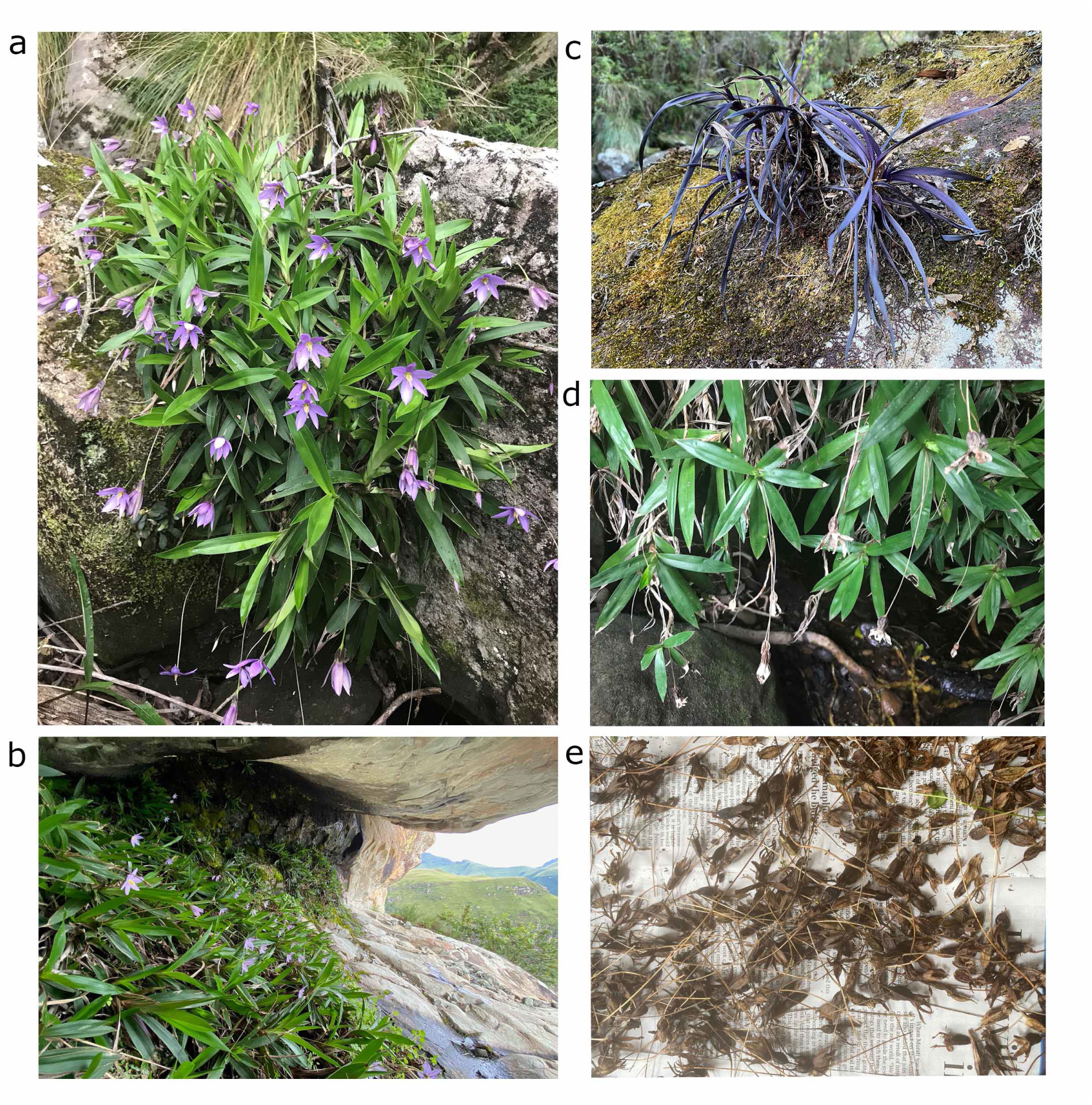
*Xerophyta elegans* in its natural habitat **a.** *X. elegans* grows abundantly in the Drakensberg, in deep shade along streams. **b**. *X. elegans* also grows in the shade of rocky overhangs. **c.** *X. elegans* dries down completely in the dry season, in winter. Purple anthocyanins mask the retained chlorophyll. **d.** *X. elegans* flowers between November and March when it sets seed. **e.** Seeds for these studies were collected from seed pods collected in the wild.

**Extended Figure 2.**
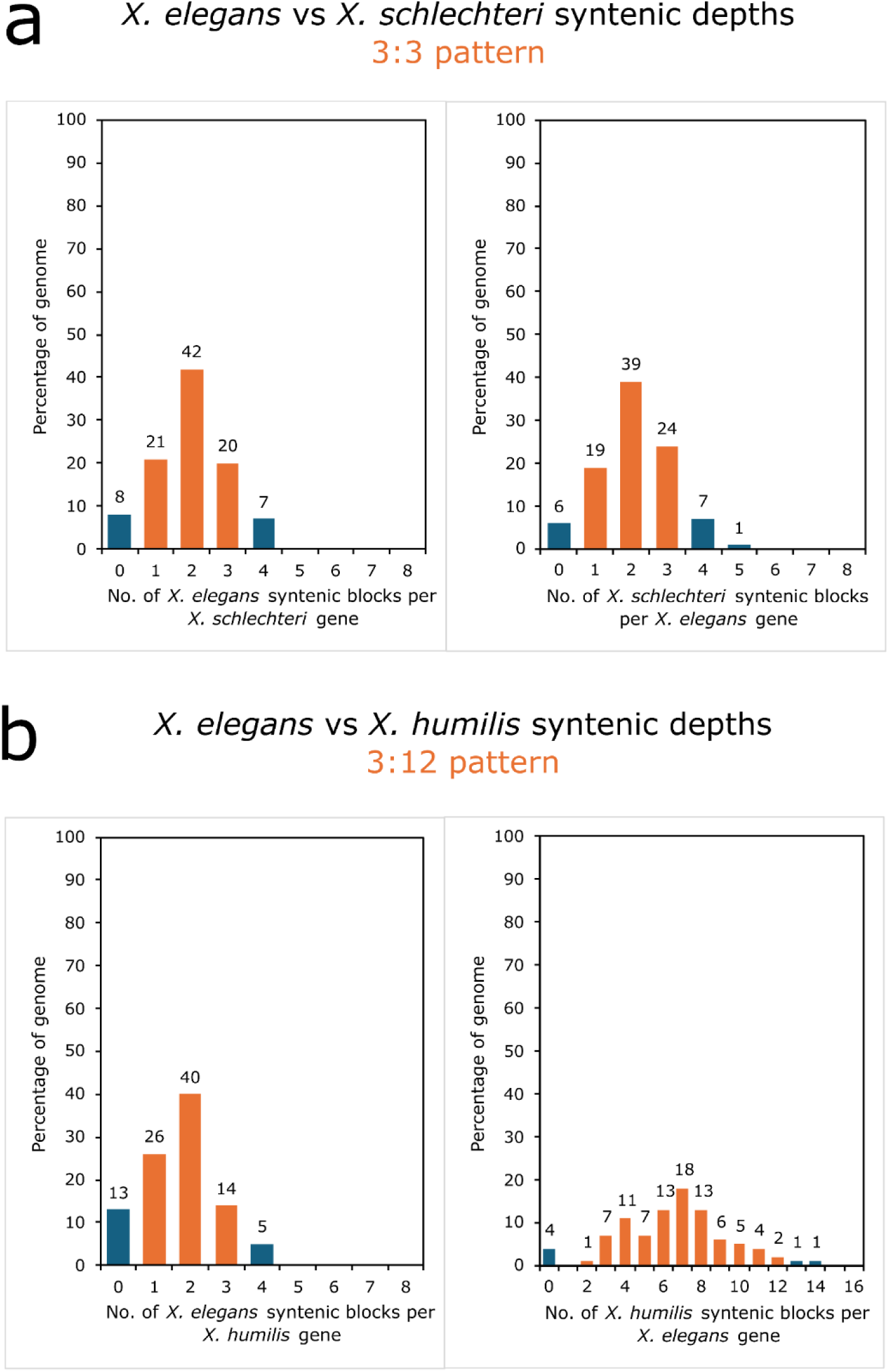
Syntenic depth in *Xerophyta* species. Analysis of the number of times a particular genomic region is found in a syntenic relationship with a genomic region of another species. The first comparison shows a 3:3 pattern for the haploid assemblies of *X. elegans* (*Xele*) vs *X. schlechteri* (*Xsch*), while the second shows a 3:12 pattern for *X. elegans* compared to the tetraploid assembly for *X. humilis* (*Xhum*).

**Extended Figure 3.**
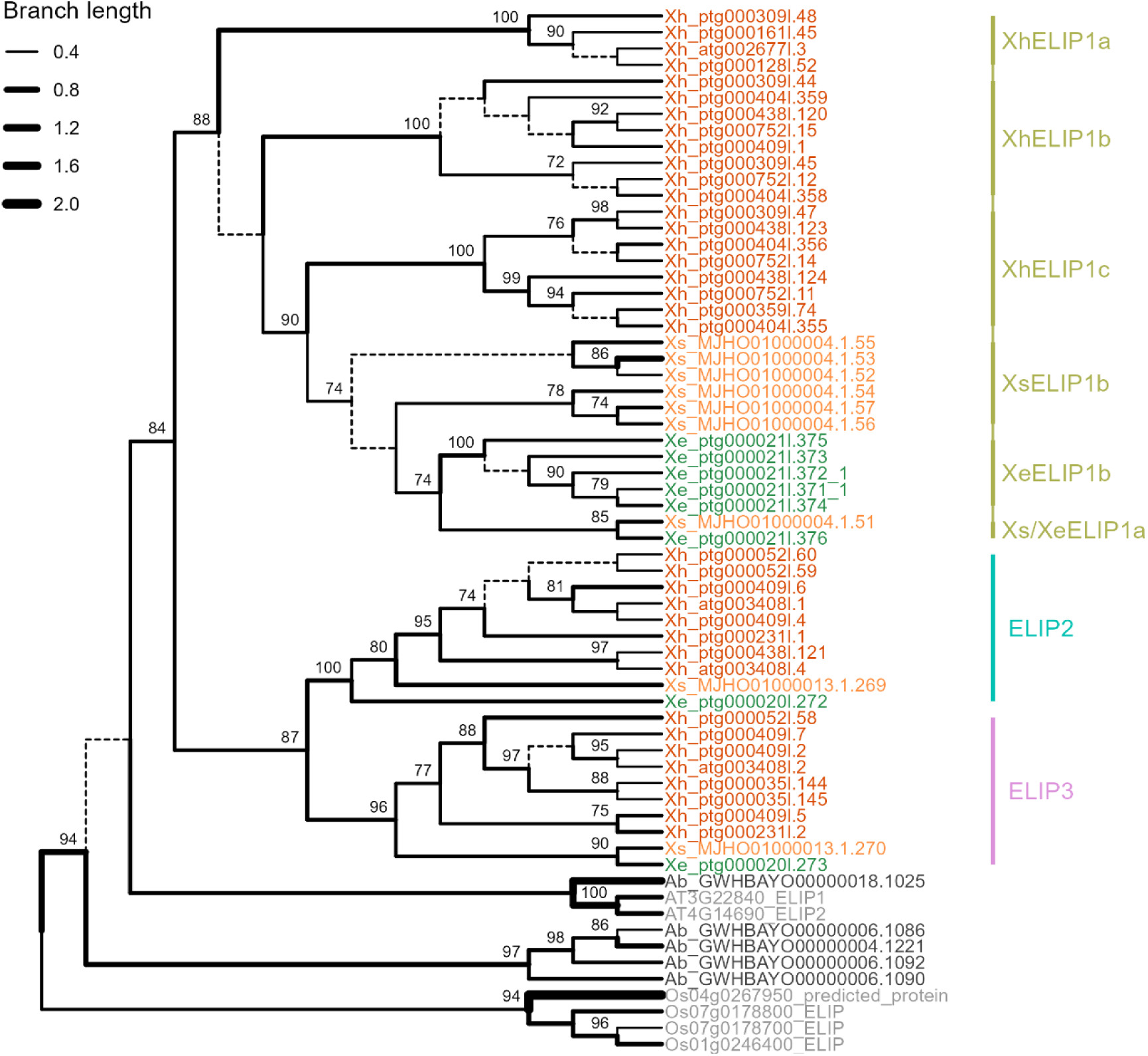
ELIP phylogeny. PhyML phylogenetic tree showing relationships between homologues in the expanded ELIP family in the haploid genome assemblies for *Xerophyta elegans (Xe)*, *Xerophyta schlechteri* (*Xs*) and *Acanthochlamys bracteata (Ab)*, in the tetraploid genome assembly for *Xerophyta humilis* (*Xh*), relative to *Arabidopsis thaliana* and *Oryza sativa*. Branch support >50% is given. *Xerophyta* clade IDs are indicated in colour.

**Extended Figure 4:**
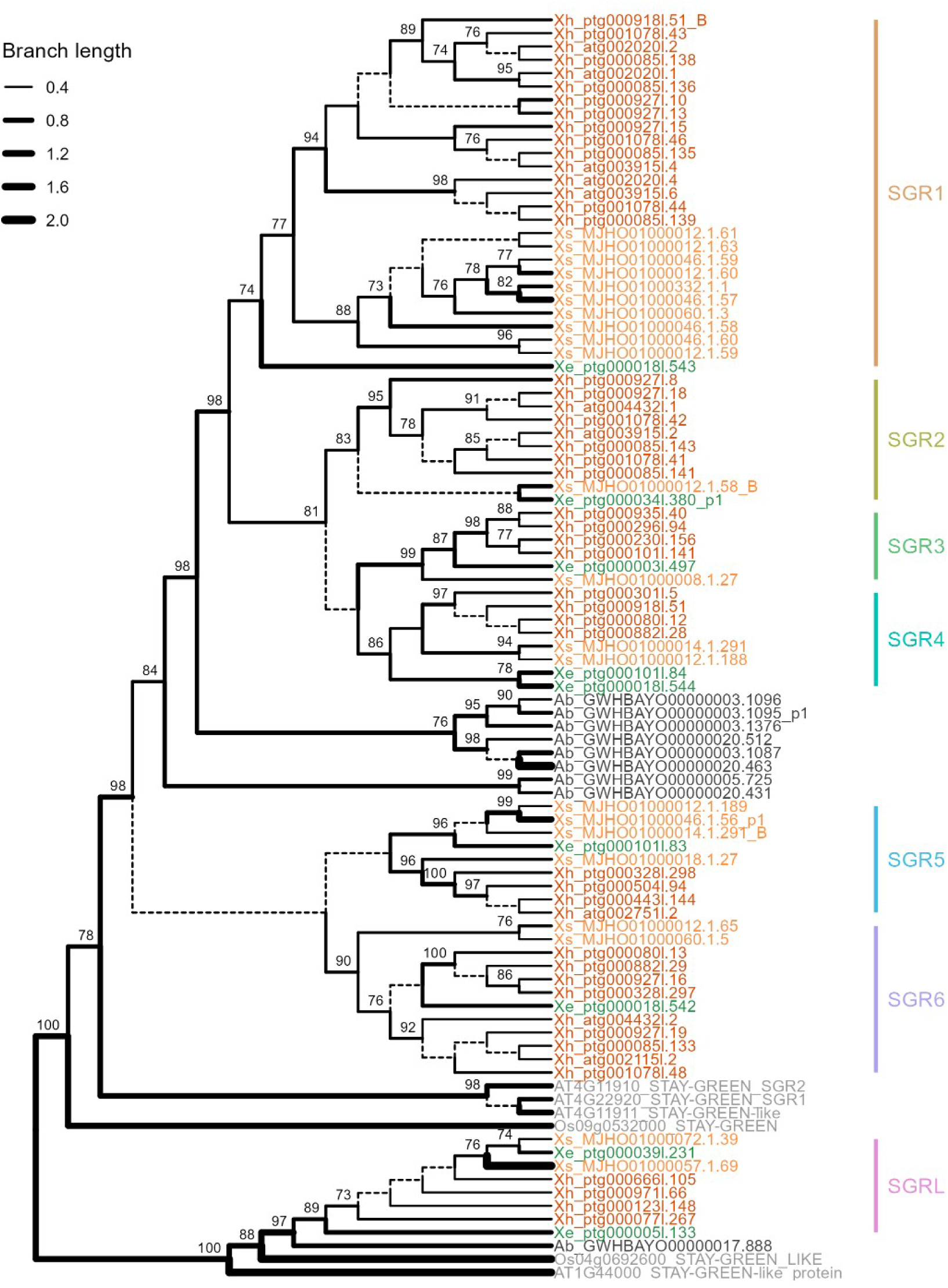
SGR phylogeny. PhyML phylogenetic tree showing relationships between homologues in the expanded *STAYGREEN* (SGR)and *STAYGREEN LIKE* (*SGRL*) family in the haploid genome assemblies for *Xerophyta elegans (Xe)*, *Xerophyta schlechteri* (*Xs*) and *Acanthochlamys bracteata (Ab)*, in the tetraploid genome assembly for *Xerophyta humilis* (*Xh*), relative to *Arabidopsis thaliana* and *Oryza sativa*. Branch support >50% is given. *Xerophyta* clade IDs are indicated in colour.

**Extended Figure 5:**
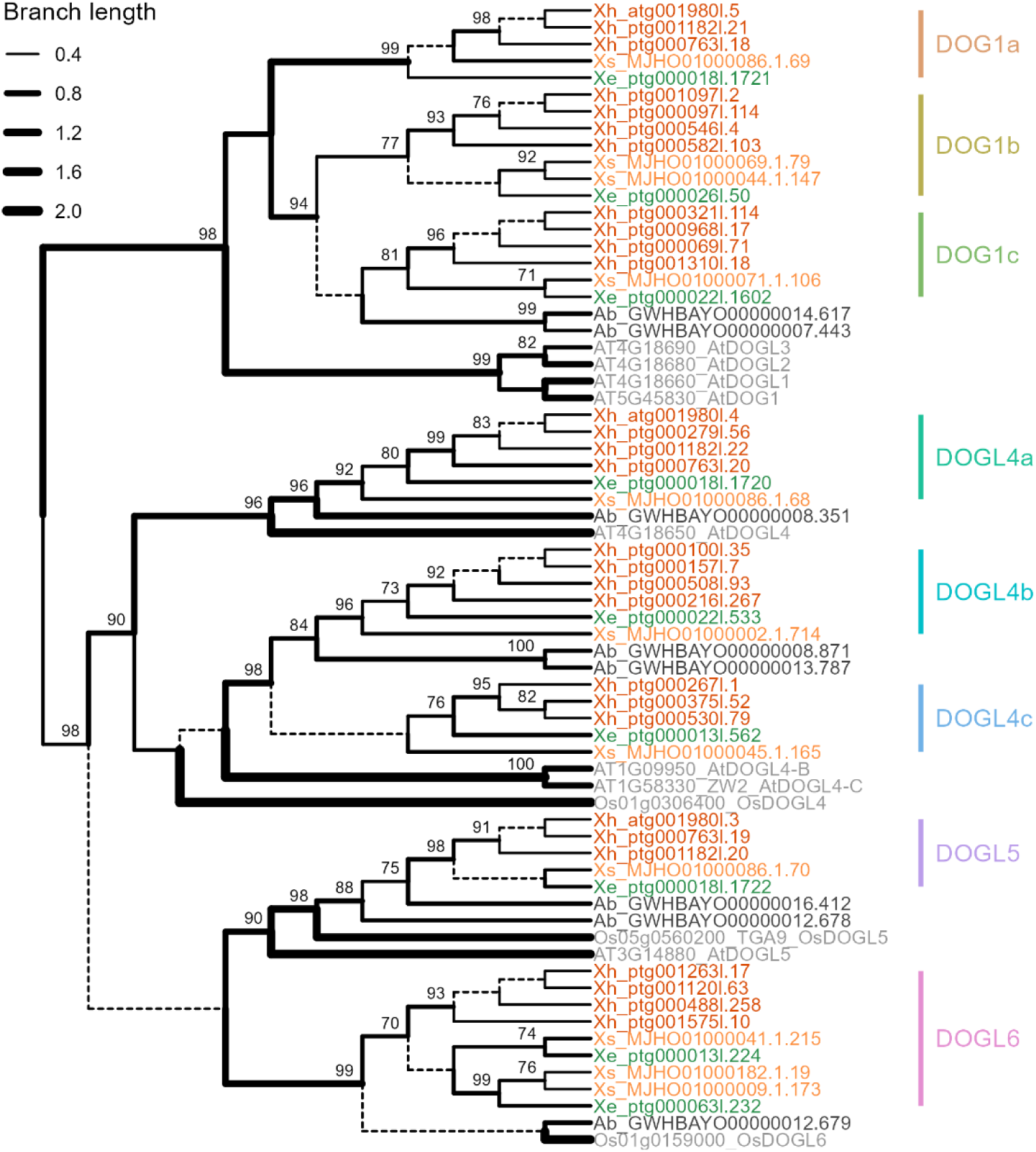
Delay of Germination (DOG) phylogeny. PhyML phylogenetic tree showing relationships between homologues in the expanded *DELAY OF GERMINATION* 1 (*DOG*1) and *DOG1-LIKE (DOGL)* gene family in the haploid genome assemblies for *Xerophyta elegans (Xe)*, *Xerophyta schlechteri* (*Xs*) and *Acanthochlamys bracteata (Ab)*, in the tetraploid genome assembly for *Xerophyta humilis* (*Xh*), relative to *Arabidopsis thaliana* and *Oryza sativa*. Branch support >50% is given. *Xerophyta* clade IDs are indicated in colour.

**Extended Figure 6:**
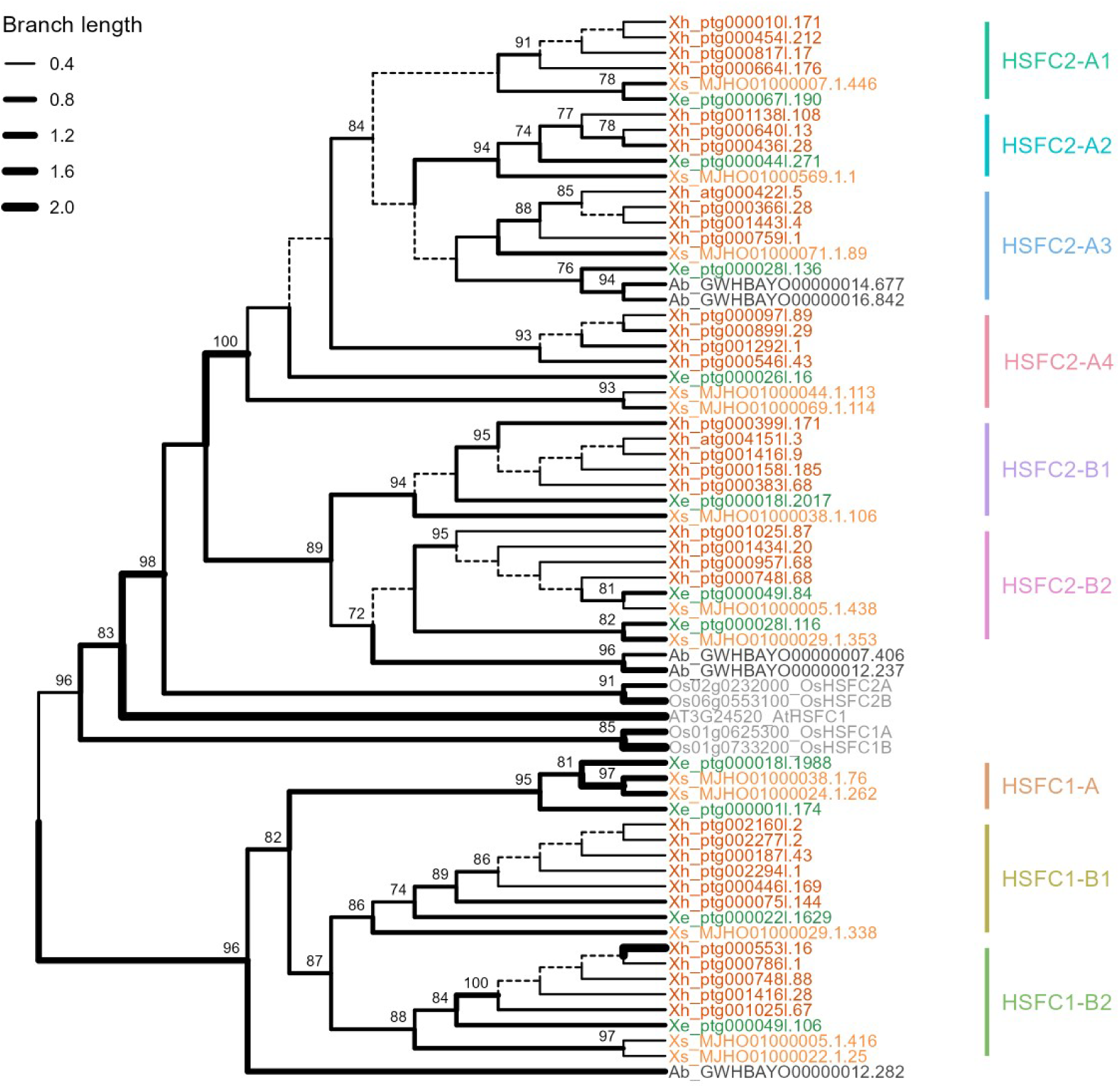
Heat Shock Factor C (HSFC) phylogeny. PhyML phylogenetic tree showing relationships between homologues in the expanded *HSFC* gene family in the haploid genome assemblies for *Xerophyta elegans (Xe)*, *Xerophyta schlechteri* (*Xs*) and *Acanthochlamys bracteata (Ab)*, and in the tetraploid genome assembly for *Xerophyta humilis* (*Xh*). There is one copy of HSFC in *Arabidopsis thaliana (AtHSFC1)* and four copies in *Oryza sativa (OsHSFC1A/B and OsHSFC2A/B)*. Branch support >50% is given. *Xerophyta* clade IDs are indicated in colour.

**Extended Figure 7.**
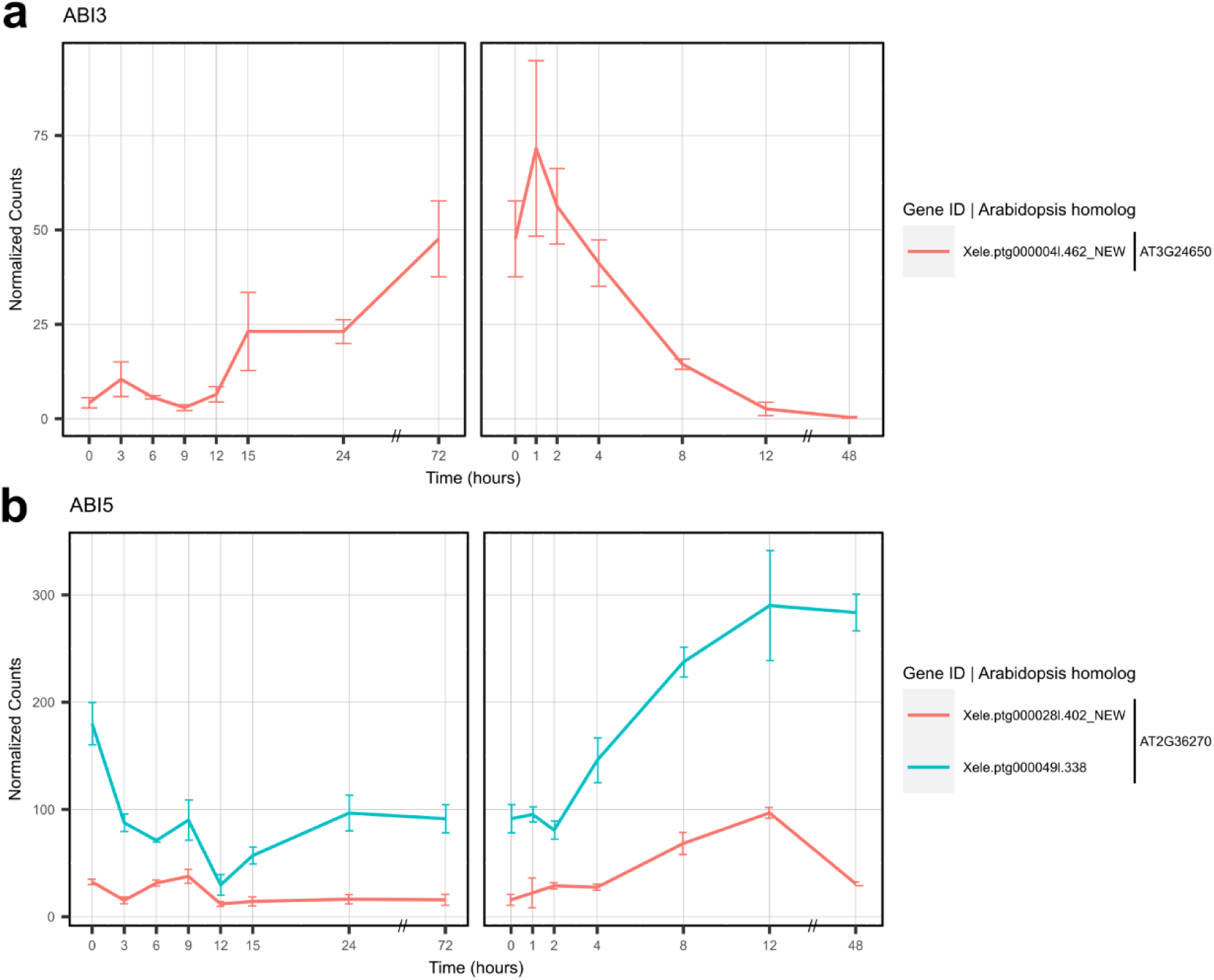
DESeq2 normalized expression values for *X. elegans ABI3* and *ABI5* homologues. The top blastp match to the Arabidopsis homologue is given for each *X. elegans* transcript. Time of sampling (hours) of *X. elegans* seedlings over a dehydration and rehydration course is given. The linear x-axis scale is discontinuous from 24 h to 72 h during dehydration and from 12 h to 48 h during rehydration. Error bars represent standard error of the mean (n = 3).

**Extended Figure 8.**
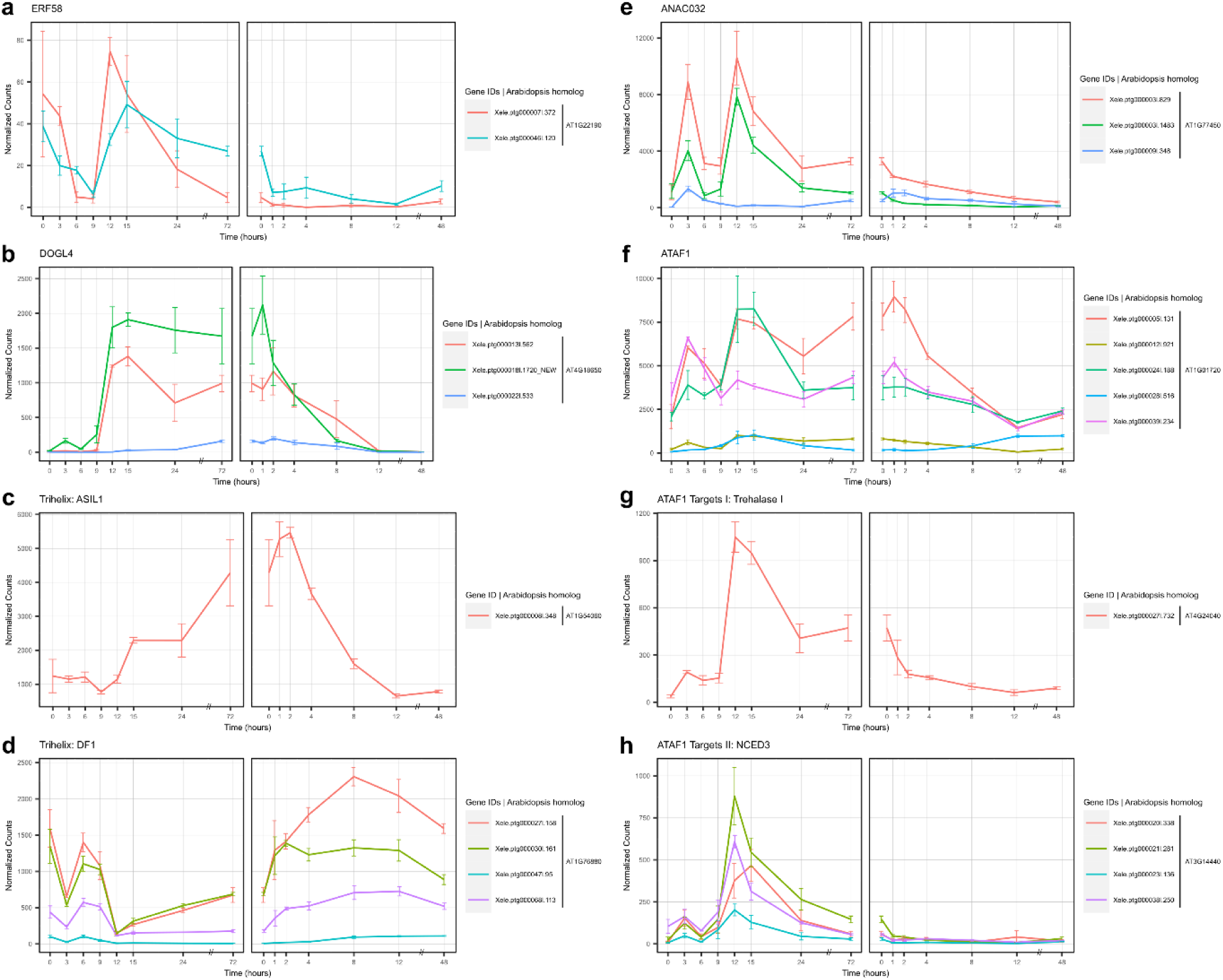
DESeq2 normalized expression values for *X. elegans* homologues of seed maturation TFs. **a)** *ERF58*, b) *DOGL4*, Trihelix c) *ASIL1* and d) DF1, and NAC TFs, e) *ANAC032* and f) *ATAF1*. Expression data for homologues of known Arabidopsis targets of ATAF1, g) Trehalase I and h) NCED3 are given. The top blastp match to the Arabidopsis homologue is given for each *X. elegans* transcript. Time of sampling (hours) of *X. elegans* seedlings over a dehydration and rehydration course is given. The linear x-axis scale is discontinuous from 24 h to 72 h during dehydration and from 12 h to 48 h during rehydration. Error bars represent standard error of the mean (n = 3).

**Extended Figure 9.**
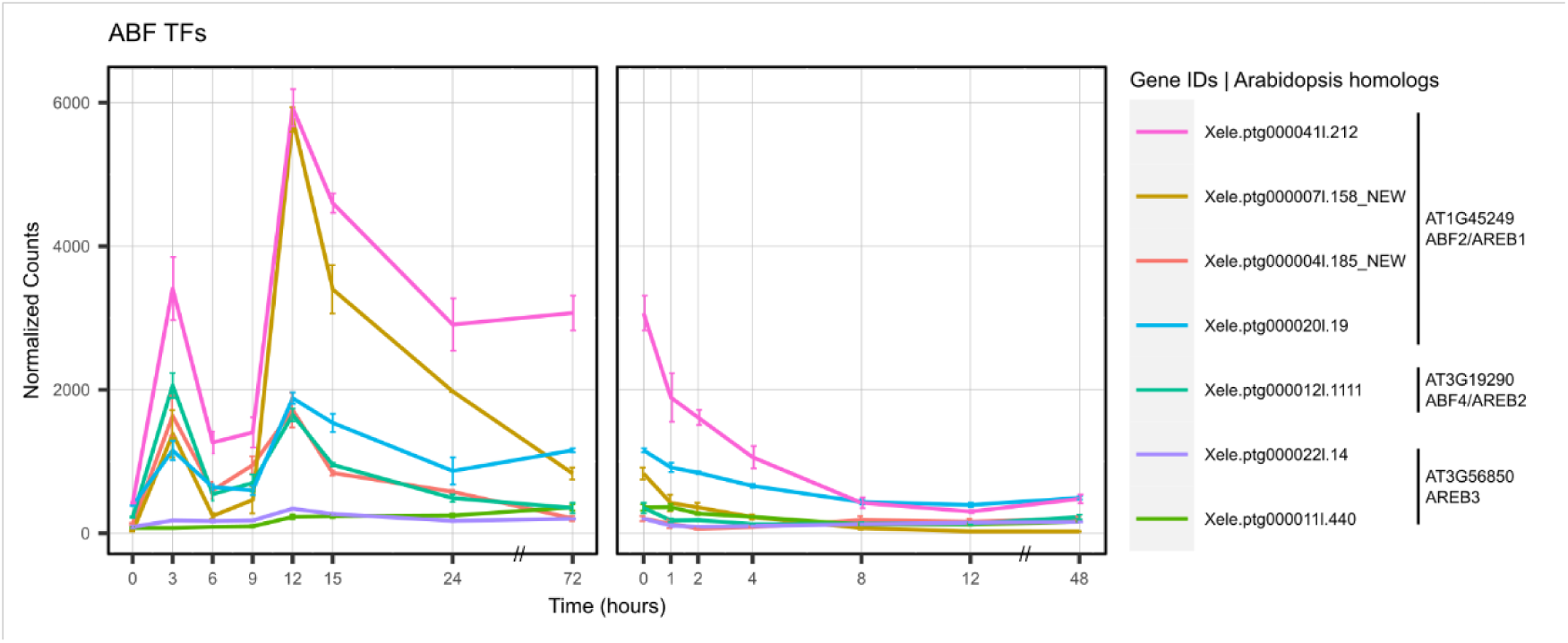
DESeq2 normalized expression values for X. elegans homologues of ABF TFs. The top blastp match to the Arabidopsis homologue is given for each *X. elegans* transcript. Time of sampling (hours) of *X. elegans* seedlings over a dehydration and rehydration course is given. The linear x-axis scale is discontinuous from 24 h to 72 h during dehydration and from 12 h to 48 h during rehydration. Error bars represent standard error of the mean (n = 3).

**Extended Figure 10:**
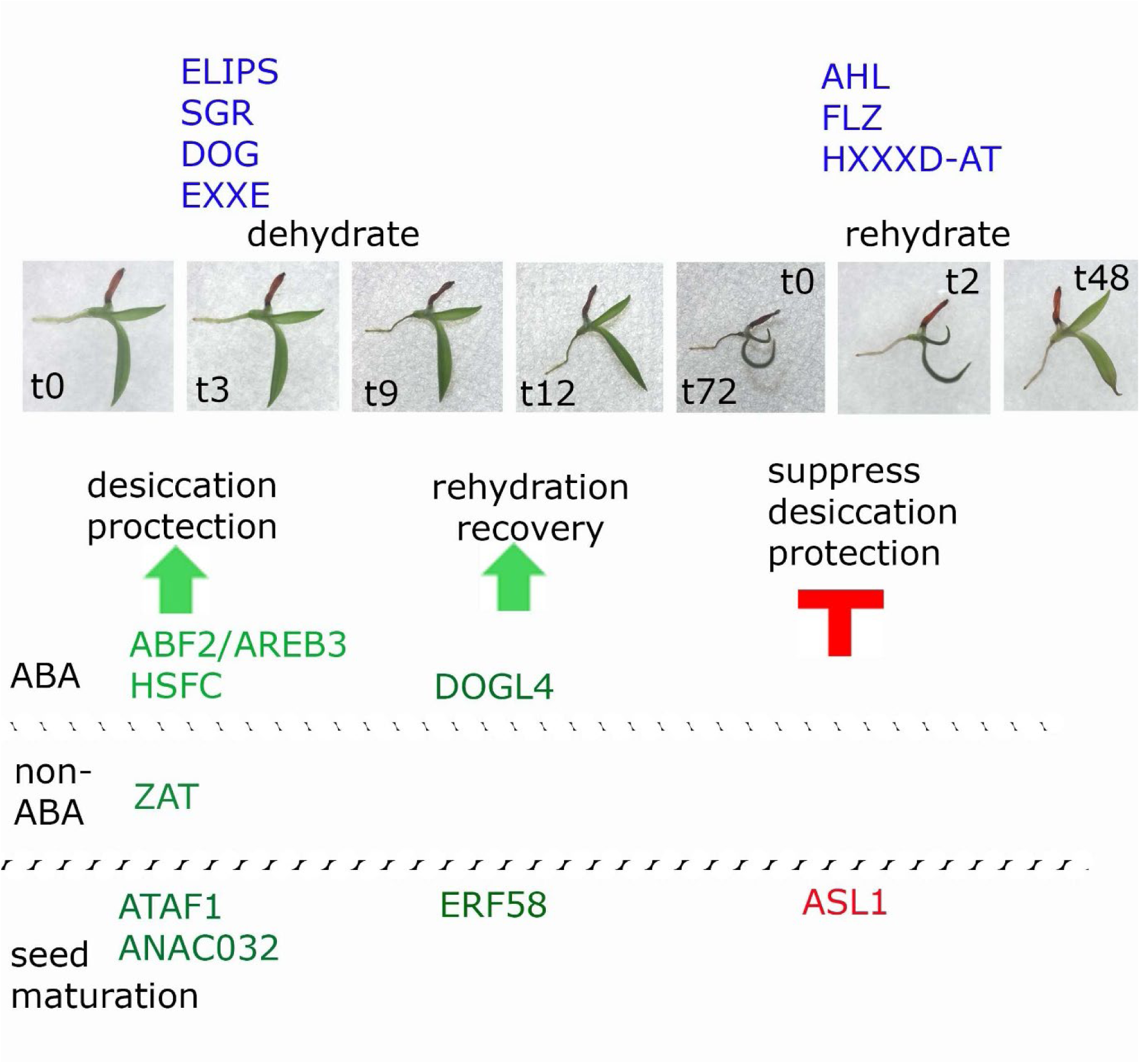
Integration of ABA signalling and seed maturation pathways regulates VDT in *Xerophyta elegans* seedlings. The bZIP *ABF2/AREB*1 and *AREB3* transcription factors (TFs) and *HSFCs* are activated in response to ABA signalling within 3 hours of dehydration. The *ZAT7* and *ZAT12* TFs are known to be induced by abiotic stresses in Arabidopsis. TFs which are linked to the regulation of seed maturation, including two NAC TFs (*ATAF1* and A*NACO32)*, *ERF58*, *DOGL4* and the Trihelix repressor *ASL1* are activated at different stages of dehydration and rehydration in *X. elegans* seedlings. The timing of activation of gene families (in blue) which are expanded in Xerophyta species is placed relative to the dehydration and rehydration of *X. elegans* seedlings. While the *ELIP*, *SGR*, *DOG* and *EXXE* gene families are important for protection from the damage incurred by desiccation, the *AHL* and *FLZ* gene families are likely to play an important role in co-ordinating the resumption of growth

