## Supplementary Note for "Transcriptional regulation of the response to water availability in the resurrection plant *Xerophyta elegans*"

### *Xerophyta* supplementary note: Additional analysis and notes on expanded gene families.

We used a combination of syntenic and phylogenetic analysis to predict the contributions of whole genome duplication events, segmental duplication and tandem gene duplication events to gene family expansion of OGs identified by CAFE analysis.

#### Materials and Methods

The synteny analysis was performed using the python version of the MCScan tool[1], which was implemented in the JCVI utilities (https://github.com/tanghaibao/jcvi/wiki/MCscan-(Python-version)). Syntenic blocks across the genomes of the various species were identified using LAST v1608[2], and blocks that contained the expanded gene families of interest were then selected for visualization. The genome assembly of *Dioscorea alata*[3] was used as the desiccation sensitive outgroup species for the synteny analyses as it was the closest related species with a high-quality chromosome-scale assembly. Analysis of syntenic depth was performed using the 'depth' function of the MCScan tool using the *X. elegans* genome assembly as the anchor for each comparison.

For the phylogenetic analysis, the amino acid sequences of rice and Arabidopsis homologues were retrieved from the OG analysis for each gene family. The online tool NGPhylogeny.fr [4] was used to align sequences by Muscle. BMGE was used to curate the sequence alignments before submitting them PhyML for phylogenetic analysis. Branch support was evaluated by approximate Likelihood Ratio Test (aLRT), and branches with less than 50% support were collapsed. Phylogenetic trees were imported and visualized in R using treeio and ggtree[5].

#### Results

*Genome duplication events*

Gene duplication is a major driving force in evolution, arising through whole-genome duplication (WGD), tandem duplication, segmental duplication or transposition[6]. Duplications can result in a gene dosage effect, subfunctionalization or neofunctionalization, all of which can facilitate adaptation to changing environments[7,8].

Several independent whole genome duplication (WGD) events have occurred in the Velloziaceae. Three ploidy levels are found across the South American and African genera, with a predicted base number of x=8. It is likely that a whole genome triplication (WGT) event occurred at the base of the *Xerophyta* lineage in Africa; *X. elegans, X. retinervis* and *X. dasylirioides* are ancient hexaploids with n=24 chromosomes[9]. It is estimated that this WGT event occurred 26 Mya in the *Xerophyta* stem lineage[10], prior to their diversification ~10 Mya [11]. It is thought that *A. bracteata* (n=20)[12] which is the earliest diverging lineage in the Velloziaceae (80-100 Mya) likely underwent an independent WGD event 24 Mya[10].

The WGT event in the *Xerophyta* lineage is supported by an analysis of syntenic depths between the assembled (1) *X. elegans* and *X. schlechteri* and (2) *X. elegans* and *X. humilis* genomes (Extended Fig. 2).

*Phylogenetic and Syntenic analysis*

We combined phylogenetic analyses with an analysis of synteny to determine how expansion of gene families associated with photoprotection and ABA signalling took place. The following assumptions were used to predict these events.

1. The expansion took place at base of Velloziaceae if family members from *A. bracteata* were represented within the clades of the *Xerophyta* expanded OGs.
2. The expansion was considered to be a consequence of WGT at the base of the *Xerophyta* genus if gene clades had representatives from all three *Xerophyta* species, i.e. were represented in multiples of three, with *A. bracteata* gene family members nested outside of these clades.
3. Segmental duplication of chromosomes was considered if genes were not directly adjacent to each other on the same chromosome.
4. Genes were considered to have been expanded by tandem duplication if they were situated adjacent to each other on the same chromosomes.

A phylogenetic analysis of several of the expanded gene families supports the ancient WGT in the *Xerophyta* genus and an independent WGD in *A. bracteata*.

The initial expansion of the *ZAT*, *PYL*, *HFS C-type (HSFC)*, *FLZ* gene families is ancient, occurring prior to the diversification of *Acanthochlamys* from the Velloziaceae lineage (Table SN1). For example, the *ZAT* family underwent an ancient expansion from one to three copies by tandem duplication. All *A. bracteata ZAT* genes cluster within the same clades as the *Xerophyta ZAT* genes confirming their ancient homology (Figure SN1). Whereas *Dioscorea alata*, a species in a sister order has single copies of *ZAT* on chromosomes 9 and 18, there are three adjacent *ZAT* copies present on several of the *A. bracteata* chromosomes and *Xerophyta* contigs (Fig. SN3-SN4).

A detailed analysis based on synteny and phylogeny for expanded gene families linked to photoprotection and ABA signalling pathways follows, together with a more detailed literature review.

| OG | PANTHER classification | Name | Expansion occurred | | | Most likely type of expansion | | | Gene clade expansion[*1] |
|---|---|---|---|---|---|---|---|---|---|
| | | | At base of Velloziaeceae | Prior to divergence of *Xerophyta* | Within *Xerophyta* species | *Xerophyta* WGT | Segmental duplication | Tandem duplication | |
| OG0000506 | PTHR26374 | *ZAT* | yes | yes | no | yes | no | yes (ancient) | three contigs, each with three *ZAT*s for Ab, Xe, Xs, Xh |
| OG0000272 | PTHR14154 | *ELIP* | no | yes | yes | yes | no | yes | *ELIP*1 (Xe, Xh, Xs)<br>*ELIP*2/3 (Xh) |
| OG0000891 | PTHR31750 | *STAYGREEN* | no | yes | yes | yes | no | yes (Xh,Xs) | SGR1 Xh (4), Xs (10)<br>SGR2 Xh (2)<br>SGR4 Xe (2), Xs (2)<br>SGR5 Xs (4)<br>SGR6 Xh (2), Xs(2)<br>SGRL Xe (2), Xs (2) |
| OG0000869 | PTHR31213 | *PYL* | yes | yes | no | yes | yes | yes (ancient) | Seven copies in all three *Xerophyta* species, adjacent copies in *A. bracteata*. |
| OG0001464 | PTHR10015 | *HSFC* | yes | yes | yes | yes | Yes (ancient) | no | Duplication of *HSFC1*, with further duplication of *HSFC1b* in Xe and Xs. Six fold expansion of *HSFC2* for all three *Xerophyta* species |
| OG0000526 | PTHR46354 | *DOG* | no | yes | yes | yes | no | yes | Specific expansion of *DOG1* and *DOGL4* families, additional |

| | | | | | | | | | |
|---|---|---|---|---|---|---|---|---|---|
| | | | | | | | | | duplication of *DOGL6* for Xe/Xs |
| OG0000179 | PTHR46057 (FLZ1,2,3) PTHR33059 (FLZ15) PTHR47847 (FLZ17,18) | *FLZ (DU581)* | yes | Yes | Yes | Yes | Yes | yes | Specific expansion of *FLZi* and *FLZv* across three *Xerophyta* species, duplication in *X.humilis* for *FLZiii* |
| OG0001453 | PTHR33782 | *EXXE* | no | Yes | no | yes | no | yes | |

Table SN1: Summary of phylogenetic and synteny analysis for the expanded gene families linked to either photoprotection or the ABA signalling pathway. Ancient refers to whether an expansion occurred prior to the diversification of the Velloziaceae.

[*1] *X. humilis* is a tetraploid, and the assembly comprises all four haplotypes. The inclusion of all four haplotypes in this analysis reinforces the conclusions drawn. However, the total number of genes in each analysis has been divided by four for comparison with *X. elegans* and *X. schlechteri*.

#### I. Photoprotection*: ZAT*s, *ELIPS* and *STAYGREEN*

Three of the expanded gene families (*ZAT*s, *ELIP*s and *STAYGREEN*) are linked to chloroplasts, having roles in photoprotection, interactions with chlorophyll and pigment binding proteins.

*ZAT*

This expanded orthogroup is the C1 subfamily of plant Zn fingers which are characterized by two Zn fingers (C-X2-C-X3-F-X5-L-X2-H-X3-H) each containing the QALGGH motif[13]. They may function as transcriptional repressors as their C-terminal domains all contain the EAR domain (ethylene-responsive element binding factor-associated amphiphilic repression domain[14]. This is the smallest known repressive domain which has been shown to bind to the ethylene-responsive GCC box.

Whereas Arabidopsis has two *ZAT* clades (C1-2iA and C1-2iB)[13], nine clades of *ZAT* can be identified in the Velloziaceae (Fig, SN1). The best characterized Arabidopsis member of this orthogroup is RHL41/*AtZAT12*. RHL41 was first identified as a cDNA that was upregulated in Arabidopsis leaves during acclimatization to high irradiation[15]. *AtZAT12* expression is also elevated in response to $H_2O_2$ accumulation[16]. Overexpression of *AtZAT12* in Arabidopsis leads to the activation of genes involved in the light and oxidative stress response, including several chloroplast genes. These plants are more tolerant to high light, osmotic and oxidative stresses[14,17].

The expansion of the C1-2iA/B family is very ancient, preceding the diversification of the Velloziaceae. Several contigs containing three tandemly repeated *ZAT* genes can be identified in *A. bracteata* and in all of the *Xerophyta* genome assemblies (Figure SN2, Figure SN3).

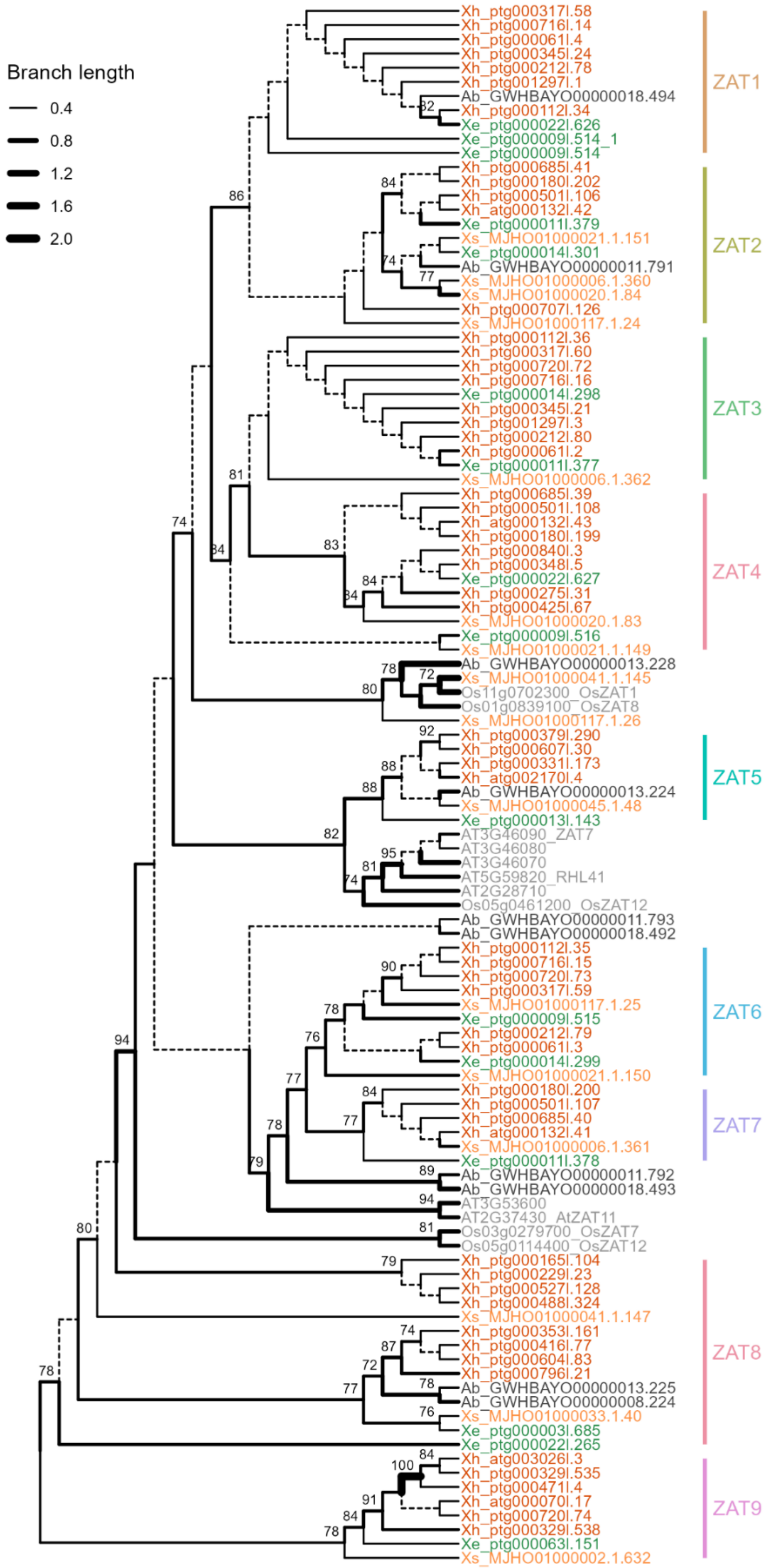


**Figure SN1**. **ZAT phylogeny**. PhyML phylogenetic tree showing relationships between homologues in the expanded ZAT family in the haploid genome assemblies for *Xerophyta elegans (Xe)*, *Xerophyta schlechteri* (*Xs*) and *Acanthochlamys bracteata (Ab)*, in the tetraploid genome assembly for *Xerophyta humilis* (*Xh*), relative to *Arabidopsis thaliana* and *Oryza sativa*.. Branch support (aLRT) >50% is given. Clades for the Arabidopsis members of this gene family are taken from Engelbrecht et al. 2024.

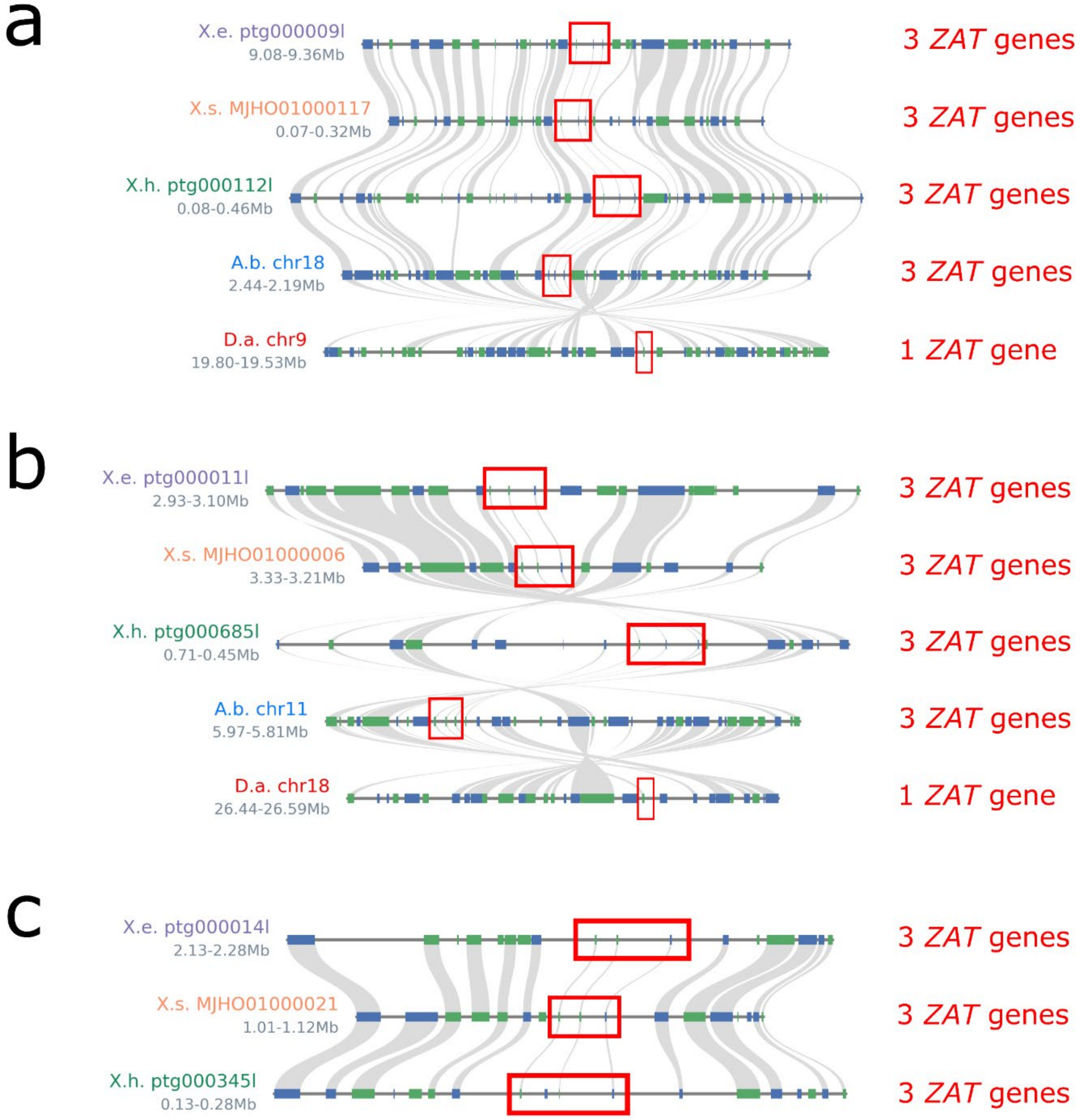


**Figure SN2. ZAT Macrosynteny plot**. Three clusters (a), (b) and (c) of ZAT homologues (indicated by red borders) are illustrated for *Xerophyta elegans* (X.e.), *Xerophyta humilis* (X.h.) and *Xerophyta schlechteri* (X.s.) relative to *Acanthochlamys bracteata* (A.b.) and *Dioscorea alata* (D. a.) . Blue blocks represent genes on the forward strand. Green blocks represent genes on the reverse strand. Text on the left represent contig names. Values on the left represent genomic locations on the contig or chromosome.

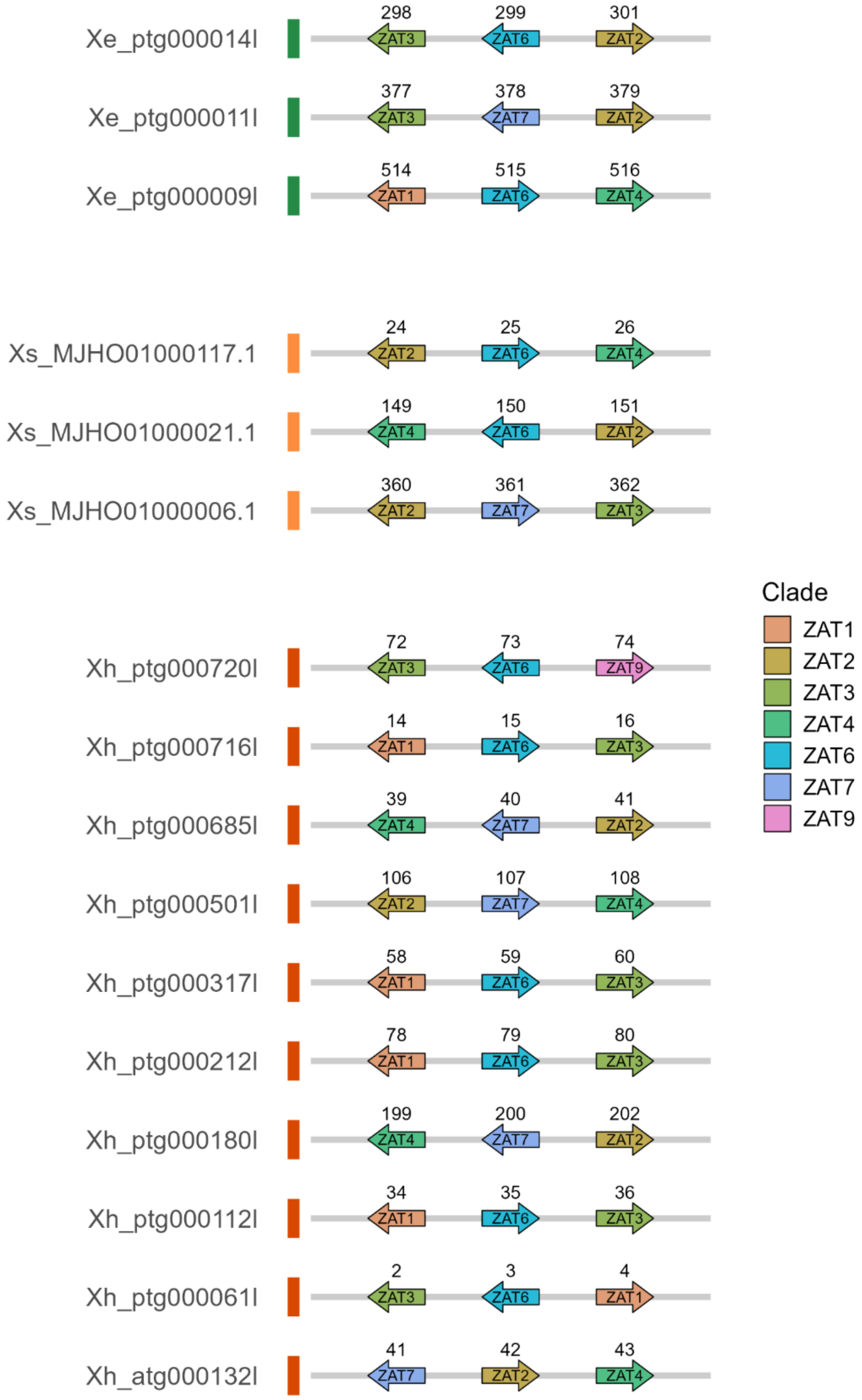


**Figure SN3. *ZAT* microsynteny plot**. Illustration of expansion of *ZAT* homologues, as defined by phylogenetic analysis, in *Xerophyta elegans* (*Xe*), *Xerophyta humilis* (*Xh*) and *Xerophyta schlechteri* (*Xs*). Colouring corresponds to clades in PhyML tree for *ZATs*. Arrows pointing to the right represent genes on the forward strand. Arrows pointing to the left represent genes on the reverse strand.

*ELIP*s

Early light-induced proteins (*ELIP*)s are important for photoprotection in plants in response to high light stress. They can function either by transiently binding to chlorophyll released during the turnover of pigment-binding proteins or by stabilization of the association of pigment-binding proteins with chlorophyll[18]. This gene family has expanded in all desiccation tolerant plants sequenced to date and these genes are expressed at high levels during desiccation and early stages of rehydration[19], likely to protect chloroplasts during desiccation from excess light energy as photosynthesis shuts down. Poikilochlorophyllous desiccation tolerant plants (*X. humilis, X. schlectheri*), *X. viscosa* dismantle the photosynthetic apparatus and thylakoid membranes completely and degrade chlorophyll. Homoiochlorophyllous desiccation tolerant plants (*X. elegans*) retain their chlorophyll, partially dismantle their photosynthetic apparatus, and retain their thylakoid membranes after dismantling grana. It has been argued that the *ELIP*s have specifically expanded in homoiochlorophyllous DT plants to protect and stabilize PSII complexes and bind free chlorophyll[19,20]. The authors noted that *X. viscosa* had the lowest number of *ELIP*s, and that this might reflect their alternative strategy of reducing photooxidative damage during desiccation.

Three *ELIP* clades are present in *Xerophyta*, with a species-specific expansion of *ELIP*1 (Extended Fig. 3). All three clades have the conserved C-terminal transmembrane domain characteristic of chlorophyll a-b binding (CAB) proteins. We do not find a correlation between *ELIP* copy number and homoiochlorophylly versus poikilochlorophylly in the same genus. Homoiochlorophyllous *X. elegans* had a similar expansion of *ELIP*1 by tandem duplication (sixfold) compared to poikilochlorophyllous *X. schlechteri* (7X expansion) (Figure SN4,5). *ELIP*s2/3 are adjacent to each other on *X. schlelchteri* and *X. elegans* contigs but have not undergone further expansion. Independent expansion of *ELIP*s has occurred in *X.humilis*, with expansion of all three classes (Figure SN7). The poikilochlorophyllous *X. humilis* and *X. schlechteri* have nine *ELIP*s per haploid genome, whereas *X. elegans* has eight. *ELIP*s must thus be playing a photoprotective role during the shutdown of photosynthesis, independent of the dismantling of the photosynthetic apparatus, breakdown of chlorophyll and turnover of thylakoid membranes.

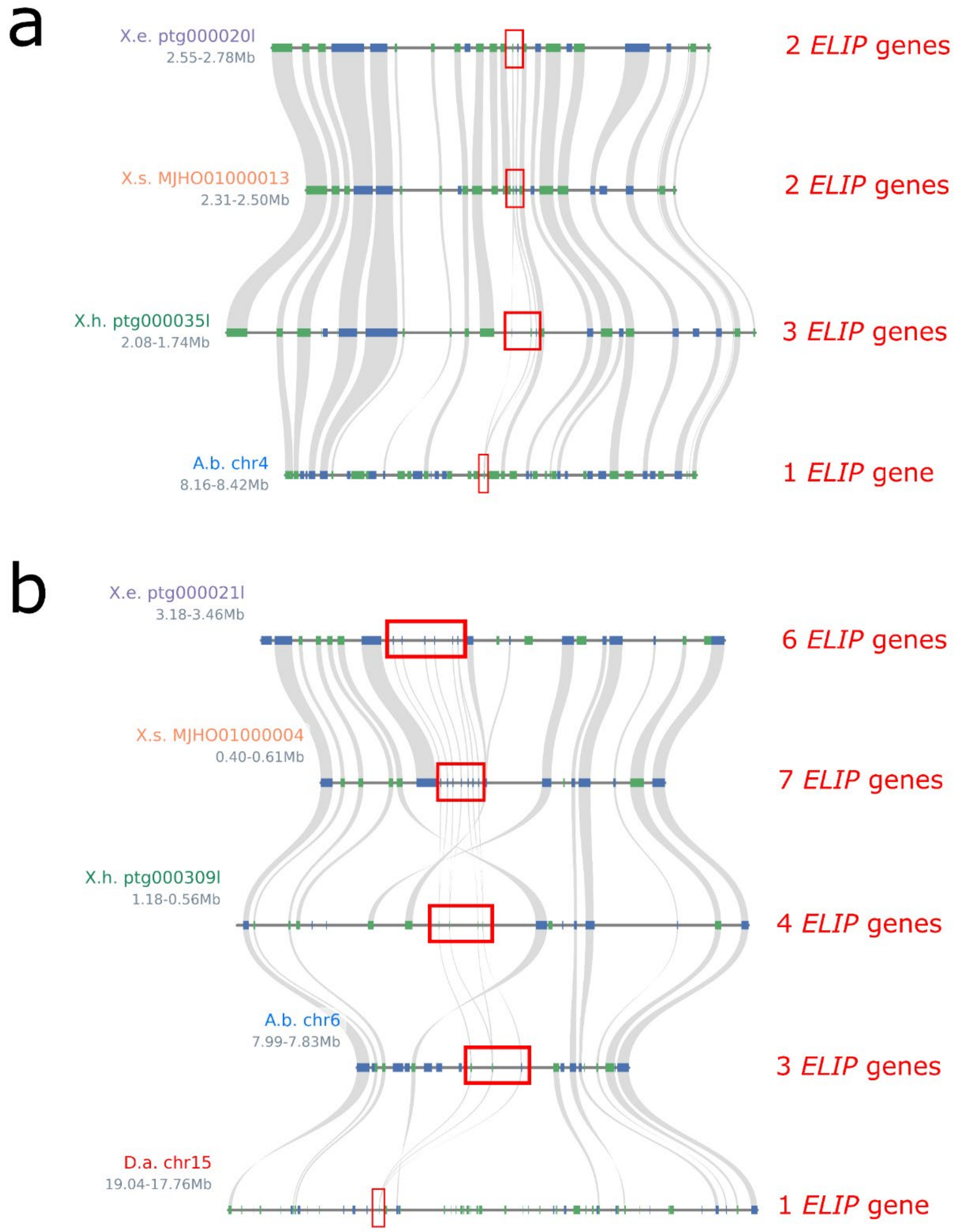


**Figure SN4. ELIP Macrosynteny plot**. Two clusters (a) and (b) of ELIP homologues (indicated by red borders) are illustrated for *Xerophyta elegans* (X.e.), *Xerophyta humilis* (X.h.) and *Xerophyta schlechteri* (X.s.) relative to *Acanthochlamys bracteata* (A.b.) and *Dioscorea alata* (D. a.) . Blue blocks represent genes on the forward strand. Green blocks represent genes on the reverse strand. Text on the left represent contig names. Values on the left represent genomic locations on the contig or chromosome.

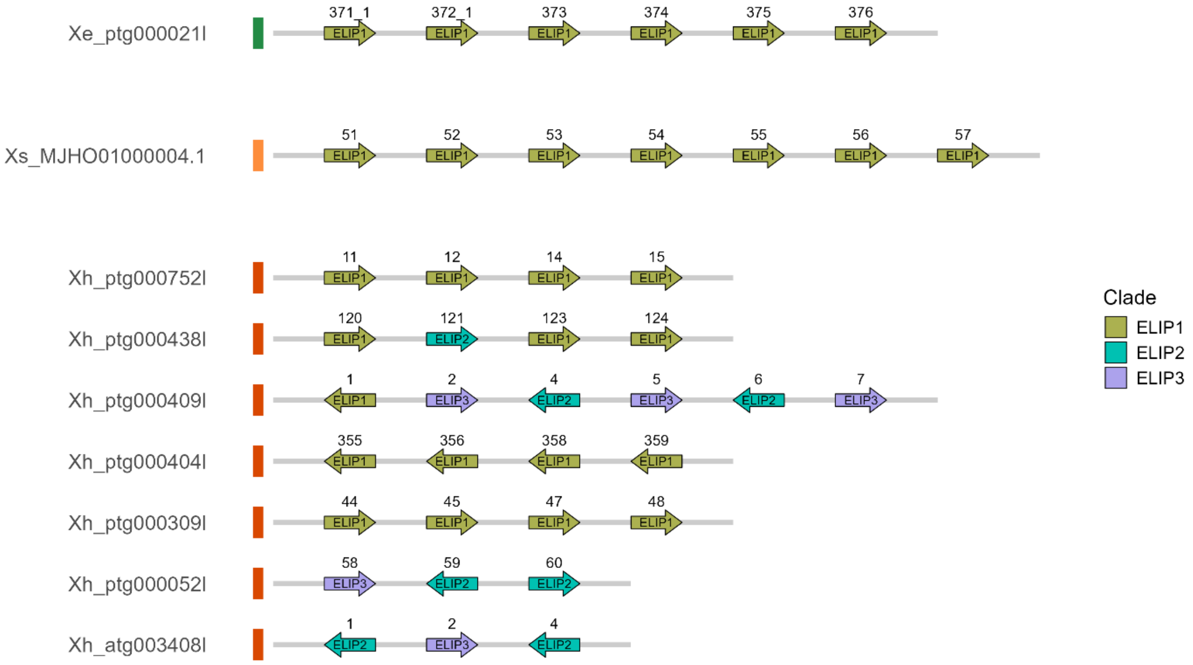


**Figure SN5. *ELIP* microsynteny plot**. Illustration of expansion of *E*LIP homologues, as defined by phylogenetic analysis, in *Xerophyta elegans* (*Xe*), *Xerophyta humilis* (*Xh*) and *Xerophyta schlechteri* (*Xs*). Colouring corresponds to clades in PhyML tree for *ELIPs*. Arrows pointing to the right represent genes on the forward strand. Arrows pointing to the left represent genes on the reverse strand.

*STAYGREEN*

The stay-green phenotype of the *sgr* mutation was originally described by Mendel[21] . Two sub-families have been defined for this gene family. The Type I SGR family includes the Arabidopsis SGR1 and SGR2 proteins, and contain both the conserved core STAY-GREEN domain and a conserved Type I SGR C-terminal domain (P-X2-C-X3C-X-C2-F-P-X5-P) that is important for the Mg-dechelatase activity. The Type II SGR family (SGR-like) lack the C-terminal domain and are referred to as STAY-GREEN LIKE (SGRL) in Arabidopsis. The expansion identified here is in the Type I SGR family and we refer to the *Xerophyta* genes in this family as SGR1-6 (Extended Fig. 4).

SGR catalyzes the first step in chlorophyll degradation by removal of the magnesium ion from the tetrapyrrole ring of chlorophyll a (Chla). This degradation of chlorophyll triggers the subsequent disassembly of the photosynthetic complexes, for example by the destabiliion of Lhcb2 in LHCII[22]. Overexpression of SGR in non-illuminated *Nicotiana benthamiana* leaves induces changes in chloroplast structure, including the accumulation of plastoglobuli[22].

The *SGR* family has undergone multiple rounds of expansion. WGD/T events lead to a two-fold expansion in *Acanthochlamys* and a three-fold expansion in *Xerophyta*. There has been a further species-specific expansion by tandem duplication of the *SGR1* genes in poikilochlorophyllous, *X. humilis* (4x) and *X. schlechteri* (10x) compared to a single *SGR1* gene in homoiochlorophyllous *X. elegans*. Other gene expansion events (Figure SN6, 7) led to a final tally of nine *SGR* family genes in *X. elegans*, compared to 22 in *X. schlectheri*, in contrast to two *SGR* genes in rice. WGCNA analysis grouped all members of the SGR family members into clusters 2 and 4, while the SGRL genes were grouped in clusters 3 and 5 (Fig. 4).

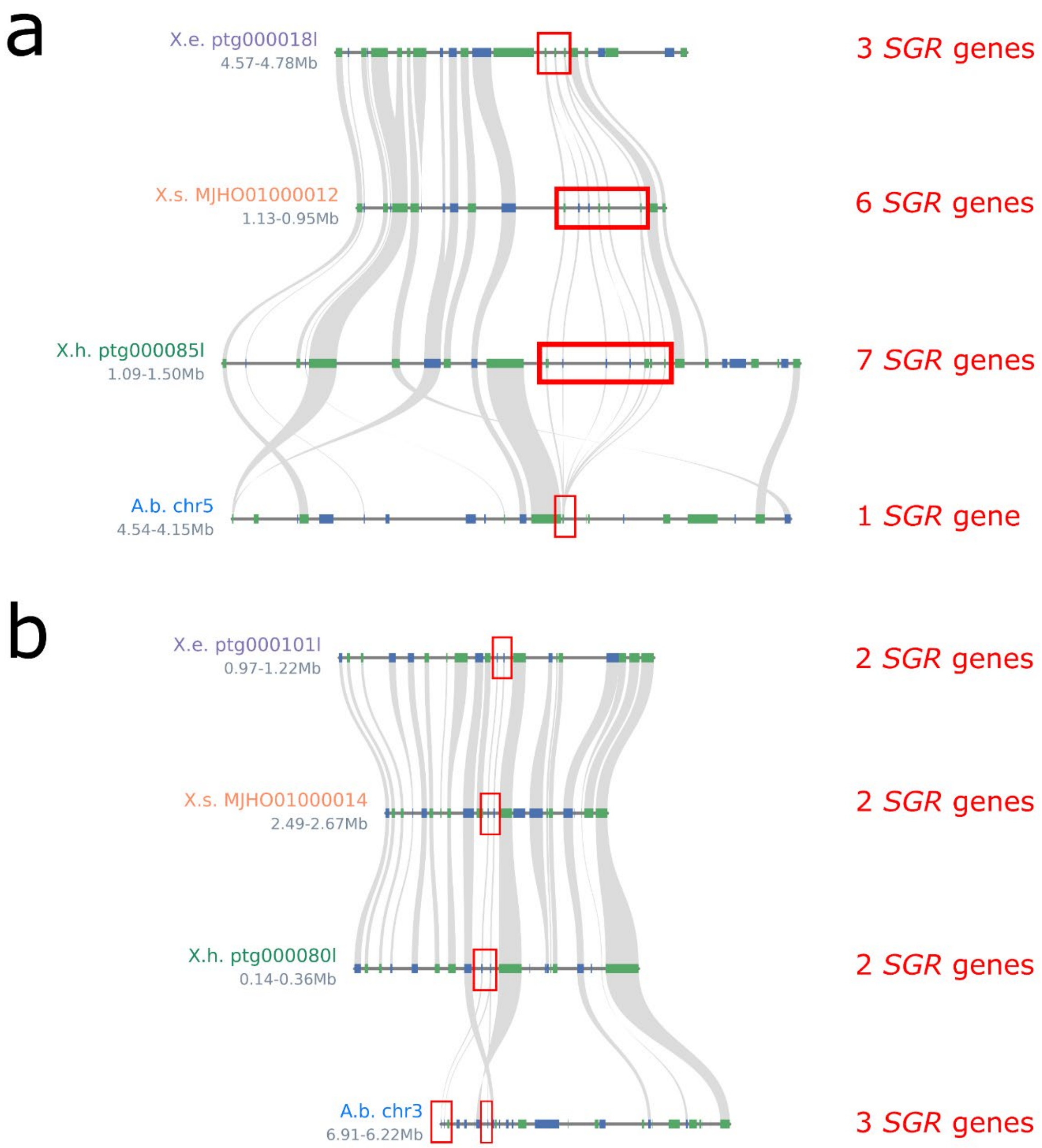


**Figure SN6. SGR Macrosynteny plot**. Two clusters (a) and (b) of SGR homologues (indicated by red borders) are illustrated for *Xerophyta elegans* (X.e.), *Xerophyta humilis* (X.h.) and *Xerophyta schlechteri* (X.s.) relative to *Acanthochlamys bracteata* (A.b.). Blue blocks represent genes on the forward strand. Green blocks represent genes on the reverse strand. Text on the left represent contig names. Values on the left represent genomic locations on the contig or chromosome.

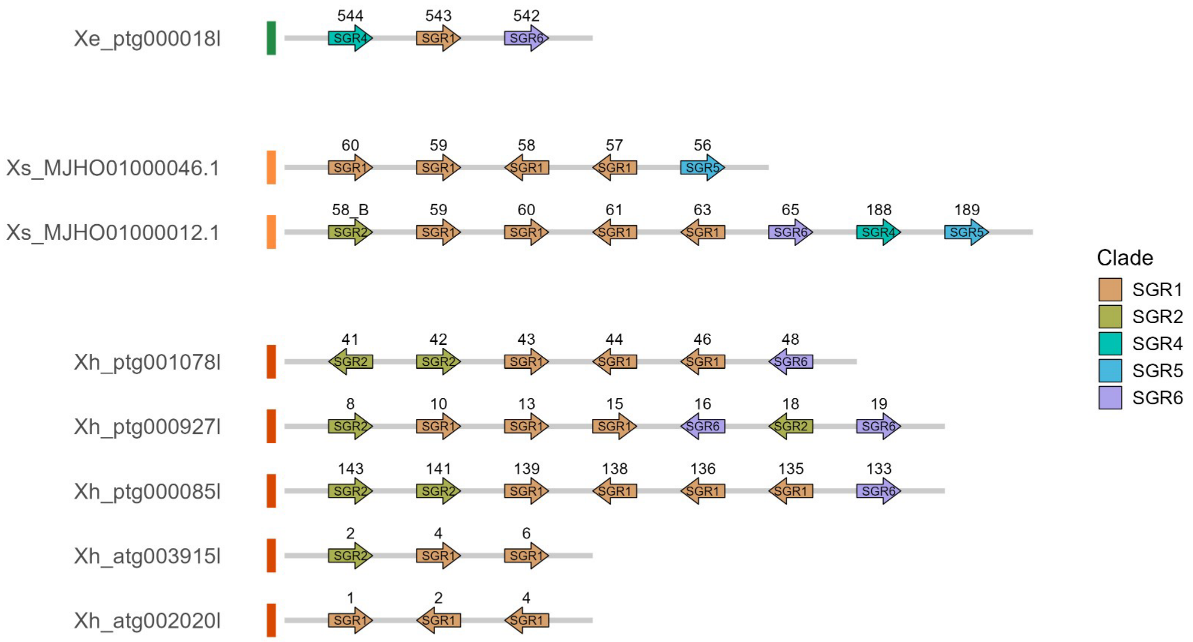


**Figure SN7. *SGR* Microsynteny plot**. Illustration of expansion of *SGR* homologues, as defined by phylogenetic analysis, in *Xerophyta elegans* (*Xe*), *Xerophyta humilis* (*Xh*) and *Xerophyta schlechteri* (*Xs*). Colouring corresponds to clades in PhyML tree for *SGRs*. Arrows pointing to the right represent genes on the forward strand. Arrows pointing to the left represent genes on the reverse strand.

#### II. ABA signalling in response to water loss:

##### *DOGs*, *PYLII*, *HSFC*

Three of the expanded gene families, *DOG*s, *PYLII*, and *HSFC* have links to the activation of the ABA signalling pathway in response to water loss. The plant Group A protein phosphatases type 2C (PP2Cs) play a central role in mediating the effects of ABA signalling during seed maturation and in response to stress. It has been argued that ancestral plants were constitutively desiccation tolerant, at the cost of growth[23]. This constitutive tolerance was suppressed by the expansion of the *PP2C* family in land plants which suppress SnRK1 and SnRK2 in hydrated conditions (Fig. 1b). Support this hypothesis comes from the observation that targeted deletion of the two Group A PP2C genes in the moss *Physcomitrella patens*, leads to a dwarf phenotype that is constitutively desiccation tolerant in the absence of ABA treatment[23]. Thus, expansion of gene families that suppress PP2C function would be predicted to enhance desiccation tolerance of vegetative tissue.

*DOG*

*DOG1* was first identified as a QTL linked to an increase in seed dormancy[24]. Homologues of four *DOG1* genes, *DOG1*, *DOGL4*, *DOGL5* and *DOGL6* are conserved across angiosperms, co-localize on the same chromosomes, and are thus likely to result from an ancient tandem duplication of the ancestral *DOG1* gene[25]. Consistent with an expansion due to the WGT event preceding the radiation of the *Xerophyta*, three copies of *DOG1* and *DOGL4* can be identified in all three *Xerophyta* species (Extended Fig. 5). The synteny of *DOG1a*, *DOGL4a* and *DOGL5* supports the ancient history of *DOG1* and *DOGL4* in angiosperms (Figure SN8-SN9). *DOGL5* is present as a single copy in the three *Xerophyta* species, as seen in rice, but there has been an expansion of *DOGL6* in *X. elegans* and *X. schlechteri* (Figure SN11). Expression of *DOG* family genes is restricted to seed maturation and important for the induction of dormancy in orthodox seeds, as well as reserve accumulation and desiccation tolerance[25].

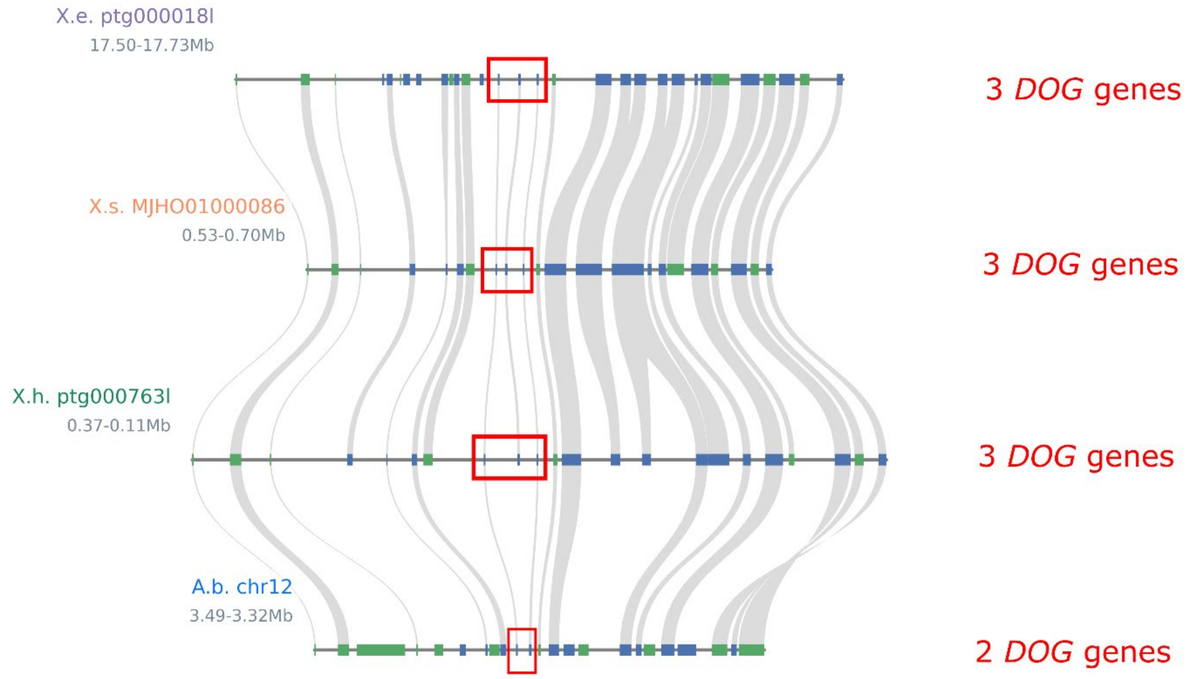


**Figure SN8. DOG Macrosynteny plot**. Relationship of homologues (indicated by red borders) are illustrated for *Xerophyta elegans* (X.e.), *Xerophyta humilis* (X.h.) and *Xerophyta schlechteri* (X.s.) relative to *Acanthochlamys bracteata* (A.b.). Blue blocks represent genes on the forward strand. Green blocks represent genes on the reverse strand. Text on the left represent contig names. Values on the left represent genomic locations on the contig or chromosome.

DOG1 regulates dormancy by interacting with Group A PP2Cs, both independently of ABA, via hypersensitive germination1 (AHG1), and via the ABA signalling pathway through AHG3. AHG1 and AHG3 are seed-specific group A PP2Cs which keep SnRK2 inactive by dephosphorylation. DOG1 binds directly to AGH1 and AGH3, suppressing their activity, independently of ABA (Figure 1b). SnRK2 activity in maturing seed is therefore released either by the presence of high levels of ABA, which bind to the PYL receptors allowing them to sequester the inhibitory AGH3, and/or by high levels of DOG1, which can sequester both AGH1 and AGH3. In addition to inactivating SnRK2, AHG1 desphosphorylates the ABI5-binding proteins (AFPs) which leads to their degradation in the proteosome[26]. Although AFPs are in fact initially activated by ABA-signalling, they are thought to be important for attenuating ABA signaling by binding to *ABI5* and the *ABF/AREBs* and suppressing their expression, together with the general co-repressor TOPLESS[26–28]. Thus, AFPs are thought to allow the escape from ABA-mediated stress responses, for the resumption of growth- for example during rehydration. AFP2 in turn, can directly suppresses *DOG1* expression, together with the co-repressor TOPLESS RELATED PROTEIN2 (TRP2) and WRKY36 transcription factor[29].

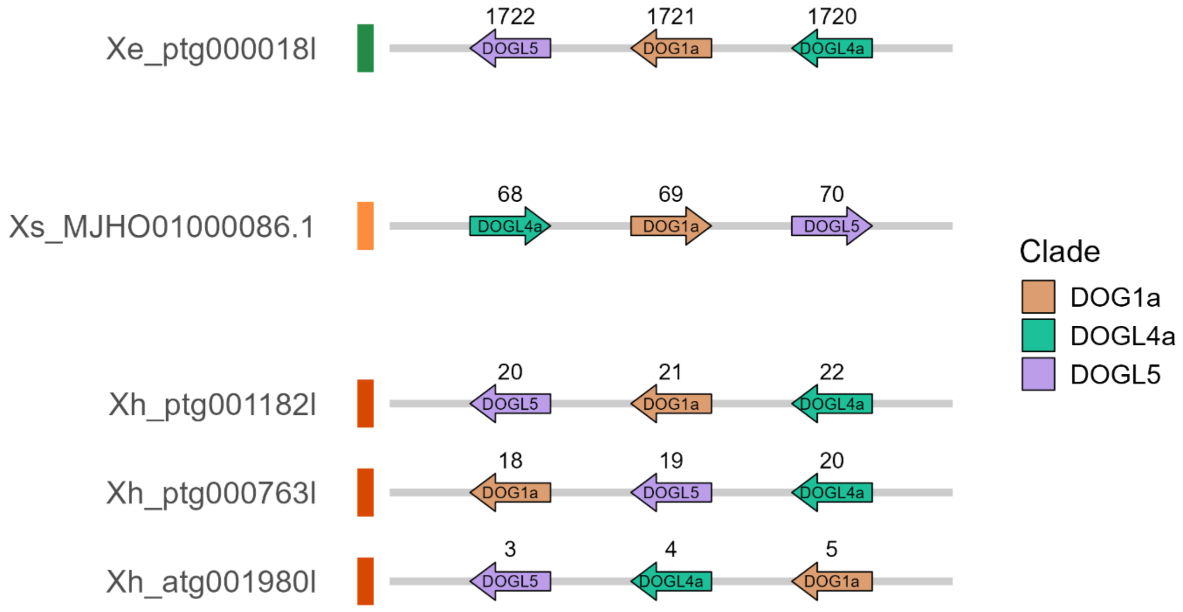


**Figure SN9. *DOG* Microsyteny plot**. Illustration of expansion of *DOG* homologues, as defined by phylogenetic analysis, in *Xerophyta elegans* (*Xe*), *Xerophyta humilis* (*Xh*) and *Xerophyta schlechteri* (*Xs*). Colouring corresponds to clades in PhyML tree for *DOGs*. Arrows pointing to the right represent genes on the forward strand. Arrows pointing to the left represent genes on the reverse strand.

*PYLII*

The *PYR* (*pyrabactin resistance*)/*PYL*(*PYR1-like*)/*RCAR* (regulatory components of ABA receptor) family proteins, hereafter referred to as *PYL*, has 14 members in Arabidopsis, all of which bind to ABA. These can be subdivided into three classes. The expanded *Xerophyta PYL* orthogroup are members of the Class II *PYR* genes (Fig. SN10) which act as monomers, and which inhibit PP2Cs in the presence of ABA[30]. Whereas rice has three copies of the Class II *PYR* genes located on different chromosomes, pairs of *Acanthochlamys* and *Xerophyta PYLII* genes cluster together, pointing to an ancient tandem duplication event, prior to the WGT in the *Xerophyta* lineage (Figs. SN11, SN12). *PYL* receptors bound to ABA, inhibit Group A PP2Cs, leading to the autophosphorylation of SnRK2, which in turn leads to the phosphorylation and activation of the ABI5 and ABF/AREB transcription factors (Fig. 1b).

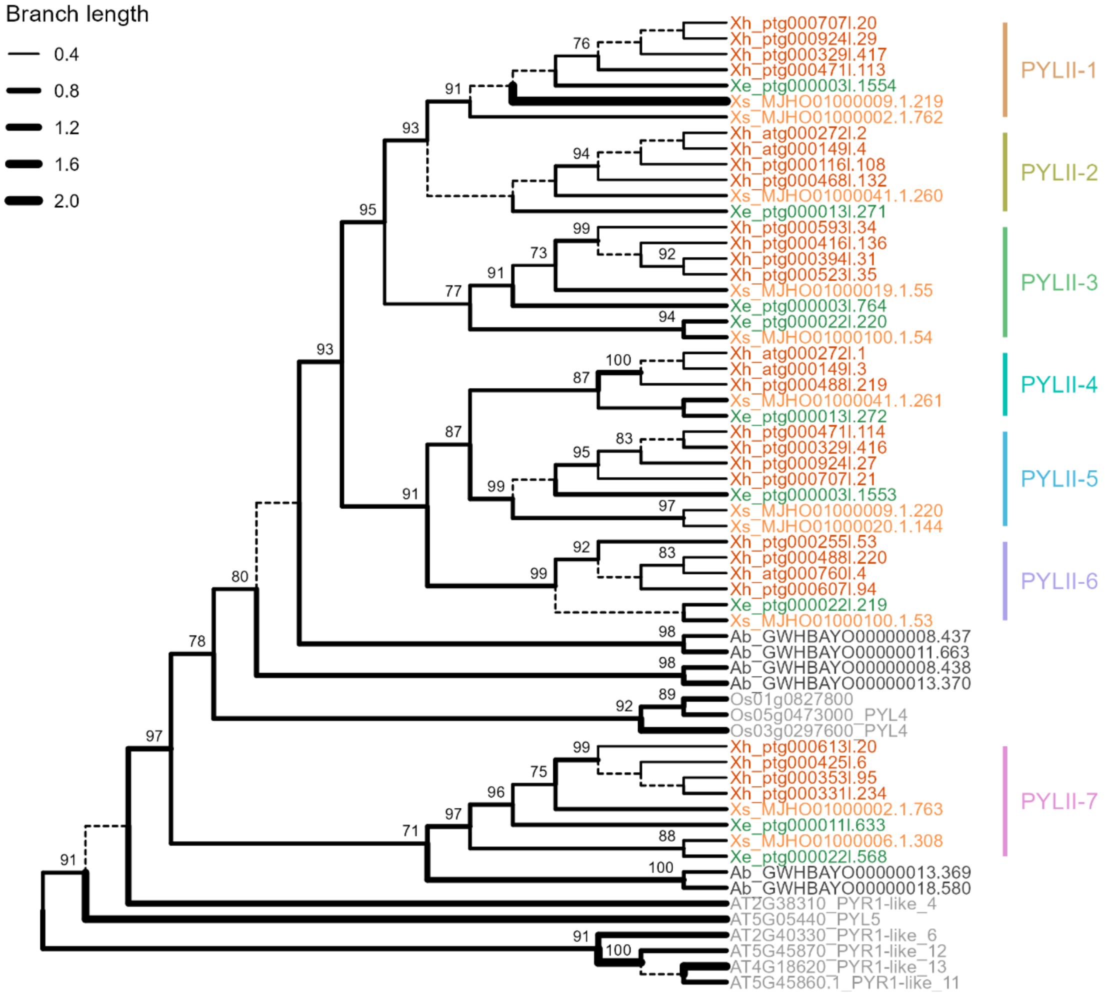


**Figure SN10: *PYLII* phylogeny**. PhyML phylogenetic tree showing relationships between homologues in the expanded *PYLII* gene family in the haploid genome assemblies for *Xerophyta elegans (Xe)*, *Xerophyta schlechteri* (*Xs*) and *Acanthochlamys bracteata (Ab)*, in the tetraploid genome assembly for *Xerophyta humilis* (*Xh*), relative to *Arabidopsis thaliana* and *Oryza sativa*. Branch support >50% is given. *Xerophyta* clade IDs are indicated in colour.

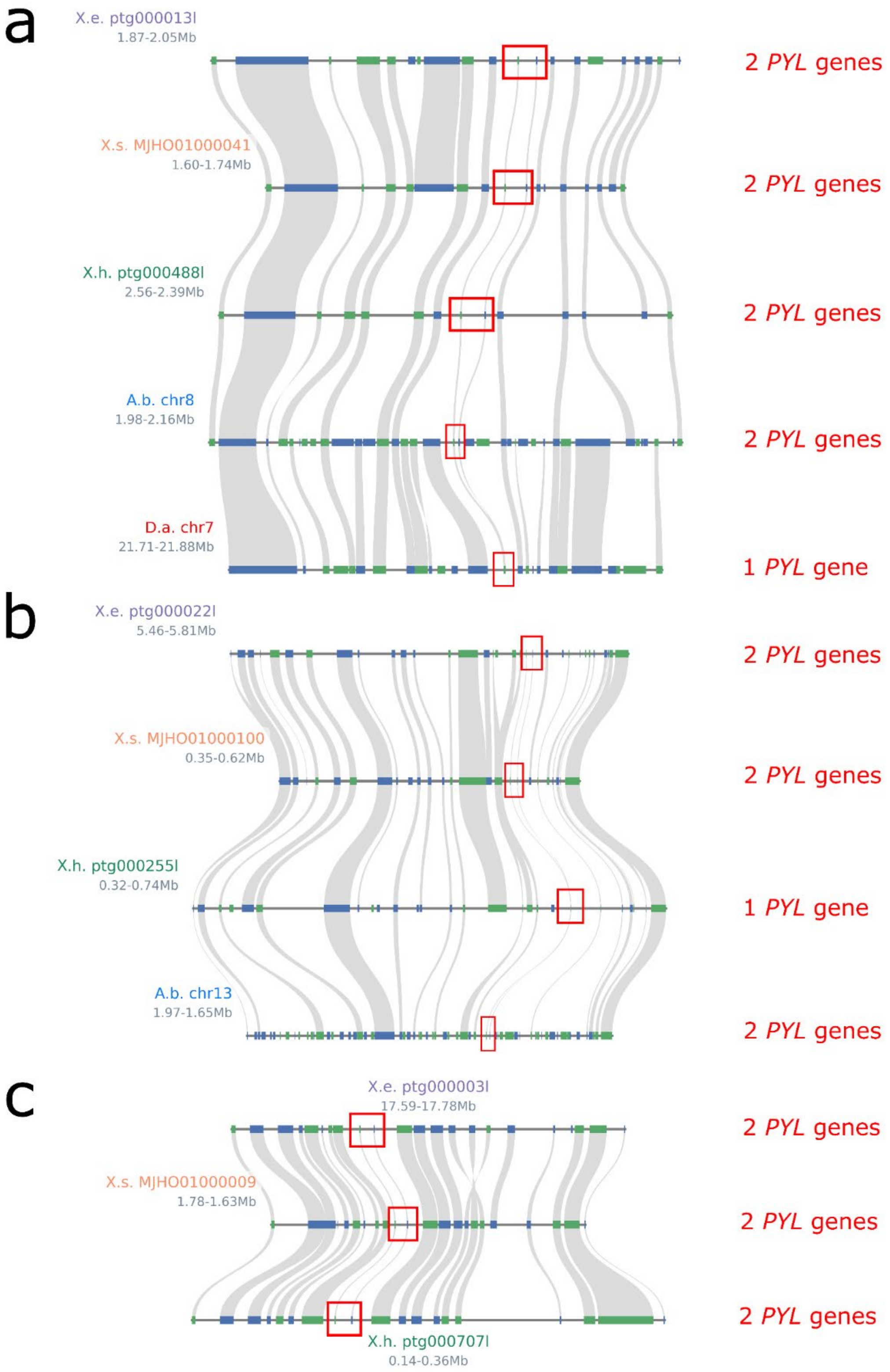


**Figure SN11. PYLII Macrosynteny plot**. Three clusters (a), (b) and (c) of homologues (indicated by red borders) are illustrated for *Xerophyta elegans* (X.e.), *Xerophyta humilis* (X.h.) and *Xerophyta schlechteri* (X.s.) relative to *Acanthochlamys bracteata* (A.b.) and *Dioscorea alata* (D. a.). Blue blocks represent genes on the forward strand. Green blocks represent genes on the reverse strand. Text on the left represent contig names. Values on the left represent genomic locations on the contig or chromosome.

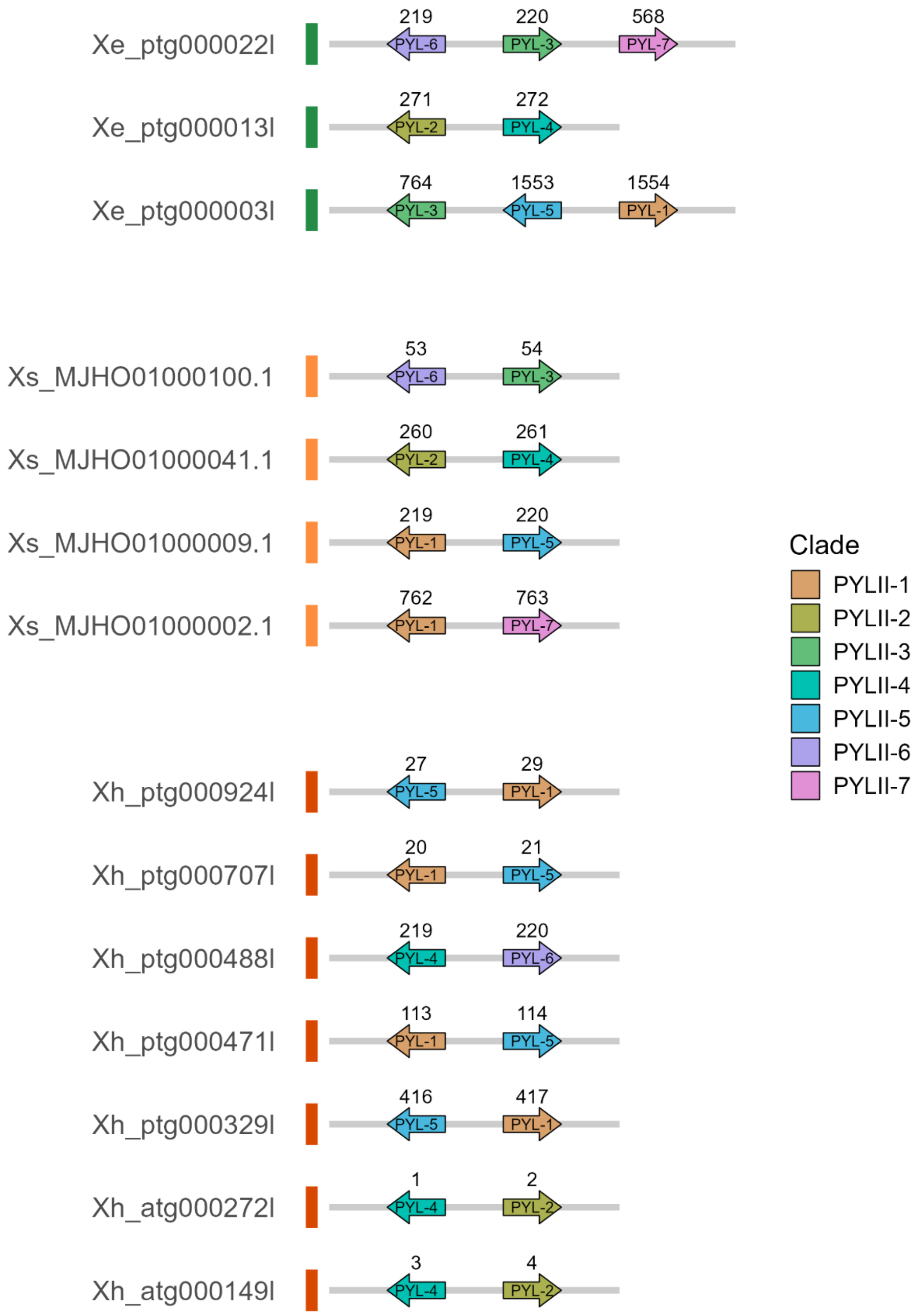


**Figure SN12 *PYL II* Microsynteny plot**. Illustration of expansion of *PYL* homologues, as defined by phylogenetic analysis, in *Xerophyta elegans* (*Xe*), *Xerophyta humilis* (*Xh*) and *Xerophyta schlechteri* (*Xs*). Colouring corresponds to clades in PhyML tree for *PYLs*. Arrows pointing to the right represent genes on the forward strand. Arrows pointing to the left represent genes on the reverse strand.

*HSFC*

The *HSFC* transcription factors are a known downstream target of ABF/AREB TFs. An analysis of genome wide microarray datasets covering eight abiotic and two biotic stress treatments ranked *HSFC1* as the most stress responsive of all the Arabidopsis HS transcription factors[31].

Monocots have two copies of *HSFC*, namely *HSFC1* and *HSFC2*[32], and this is also the case in the Velloziaceae (Extended Fig. 6). *A. bracteata* has a copy of the *HSFC1* and *HSFC2* gene at either ends of chromosome 12, in addition to other chromosomes with single copies of *HSFC2* genes. This pattern of two, non-adjacent *HFSC* genes on a single contig, is conserved for three contigs in *X. elegans* and *X. schlectheri*, corresponding to the WGT of this ancient segmental duplication. This WGT also accounts for the further expansion of the 'single *HSFC2*' genes in *X. elegans* and *X. schlechteri* (Fig. SN13).

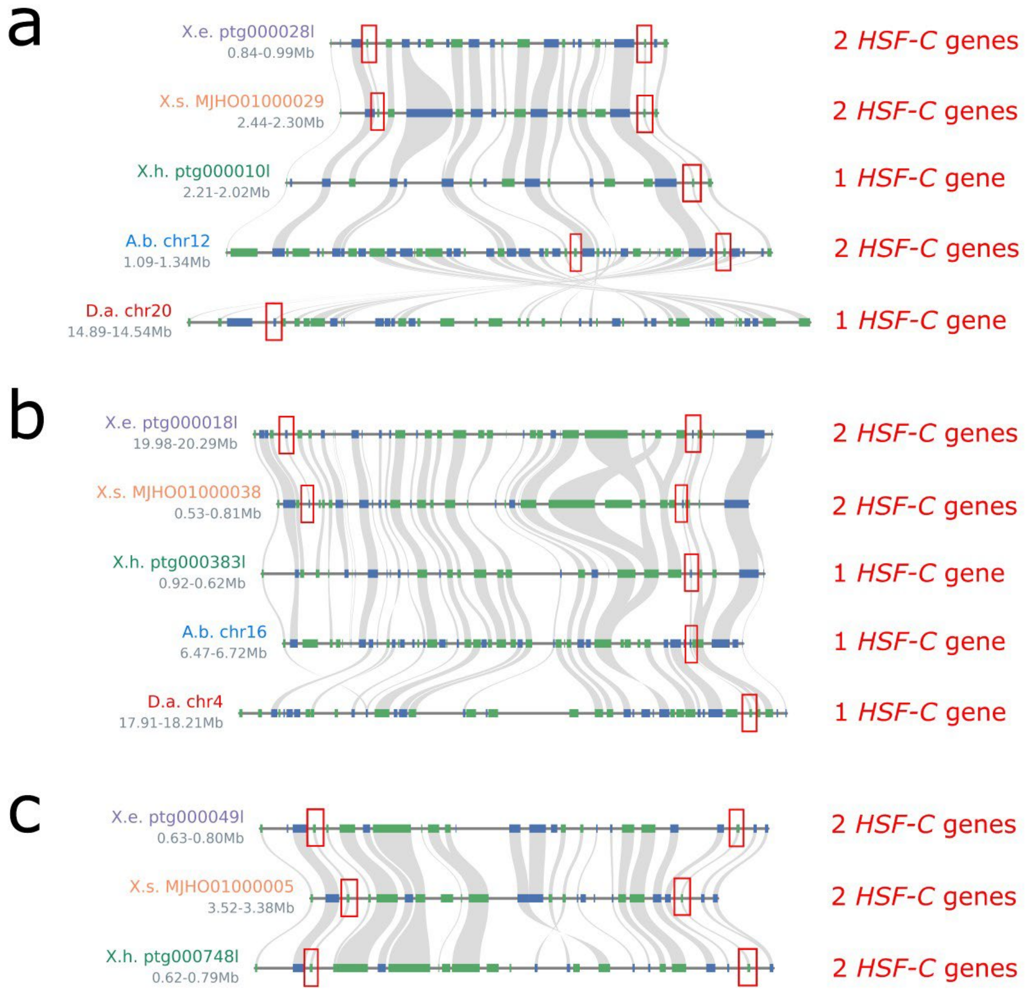


**Figure SN13. *HSFC* Macrosynteny plot**. Three clusters (a), (b) and (c) of *HSFC* homologues (indicated by red borders) are illustrated for *Xerophyta elegans* (X.e.), *Xerophyta humilis* (X.h.) and *Xerophyta schlechteri* (X.s.) relative to *Acanthochlamys bracteata* (A.b.) and *Dioscorea alata* (D. a.). Blue blocks represent genes on the forward strand. Green blocks represent genes on the reverse strand. Text on the left represent contig names. Values on the left represent genomic locations on the contig or chromosome.

**3. Rehydration and Resumption of Growth: FLZ and Unknown (EXXE)**

The *FLZ* and *EXXE* family genes may play a role in restricting SnRK1 activity ensuring plant growth in favourable conditions (Fig. 1b).

*FLZ*

The *FCS-like Zn finger* (*FLZ*) family has 18 members in Arabidopsis, which can be divided into five subfamilies[33]. This expanded orthogroup comprises three of the subfamilies, i, iii and v, with the expansion occurring primarily in the i subgroup. The *FLZ* family is characterized by the DUF581 domain, which has been shown to interact with SnRK1[34].

Whereas SnRK2 mediates the response of the plant to ABA, SnRK1 is important for energy sensing and suppressing growth under stress. There is substantial overlap in transcriptional targets of these pathways[35]. The Arabidopsis *FLZi* protein (At2g44670/FLZ3) DUF581-9, has been shown to directly interact with the catalytic subunit of SnRK1a1 to repress its activity under energy-sufficient conditions[36]. A second member of this subfamily, DUF581-12/FLZ2 (At4g17670) was identified as one of the core SnRK1a1 interactors in a Y2H screen[35].

The expansion of the *FLZi* subfamily genes by tandem duplication is ancient, preceding the diversification of *Acanthochlamys* from the Velloziaceae (Fig. SN20-SN21). The ancestral cassette is predicted to have contained three subfamilies of the *FLZi* gene. Coupled with WGT events, there has been a nine-fold expansion of the *FLZi* genes in *Xerophyta*, and a two-fold expansion of *FLZv* across all three *Xerophyta* species.

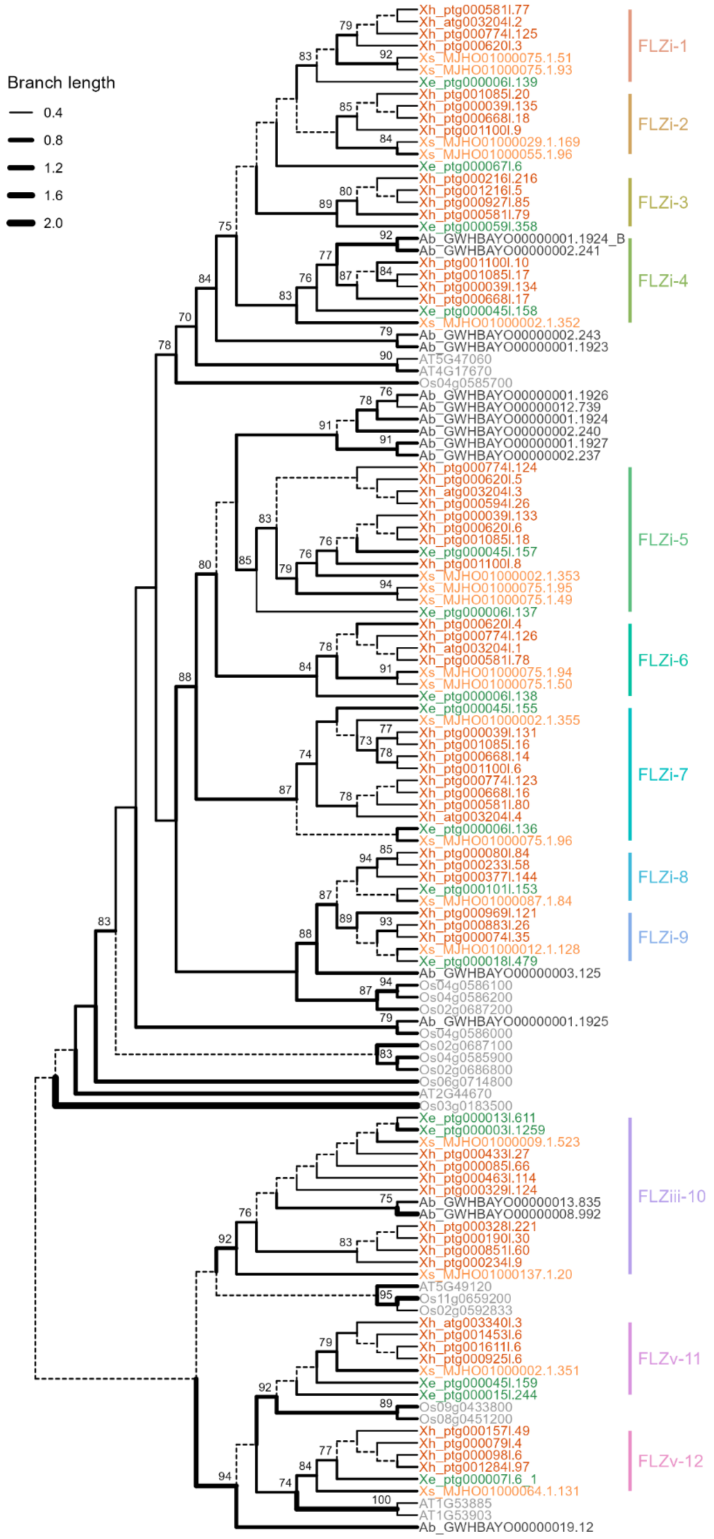


**Figure SN14: *FLZ* phylogeny**. PhyML phylogenetic tree showing relationships between homologues in the expanded *FLZ* gene family in the haploid genome assemblies for *Xerophyta elegans (Xe)*, *Xerophyta schlechteri* (*Xs*) and *Acanthochlamys bracteata (Ab)*, in the tetraploid genome assembly for *Xerophyta humilis* (*Xh*), relative to *Arabidopsis thaliana* and *Oryza sativa*. Branch support >50% is given. *Xerophyta* clade IDs are indicated in colour.

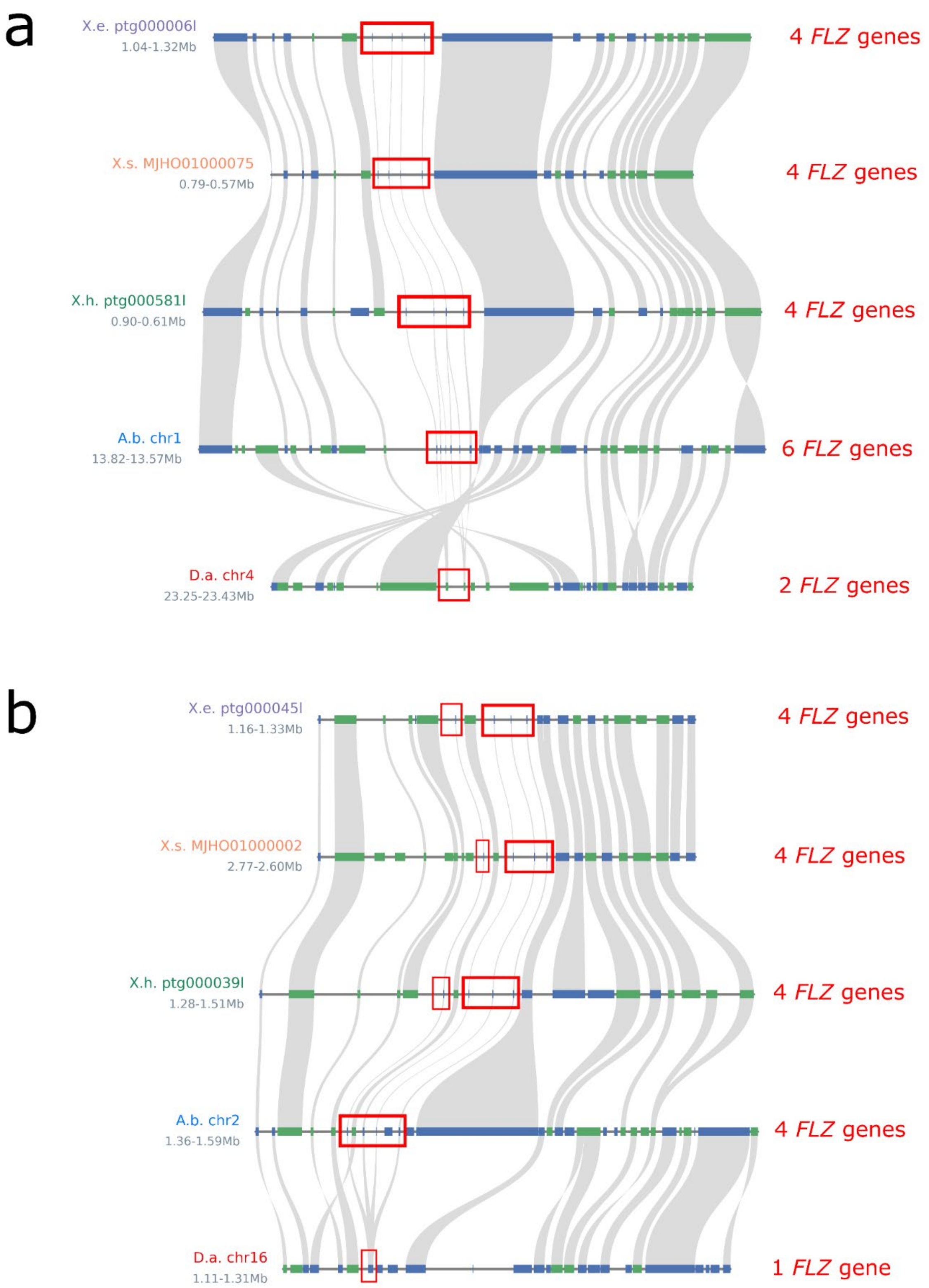


**Figure SN15. *FLZ* Macrosynteny plot**. Two clusters (a), (b) of *FLZ* homologues (indicated by red borders) for *Xerophyta elegans* (X.e.), *Xerophyta humilis* (X.h.) and *Xerophyta schlechteri* (X.s.) relative to A*canthochlamys bracteata* (A.b.) and *Dioscorea alata* (D. a.). Blue blocks represent genes on the forward strand. Green blocks represent genes on the reverse strand. Text on the left represent contig names. Values on the left represent genomic locations on the contig or chromosome.

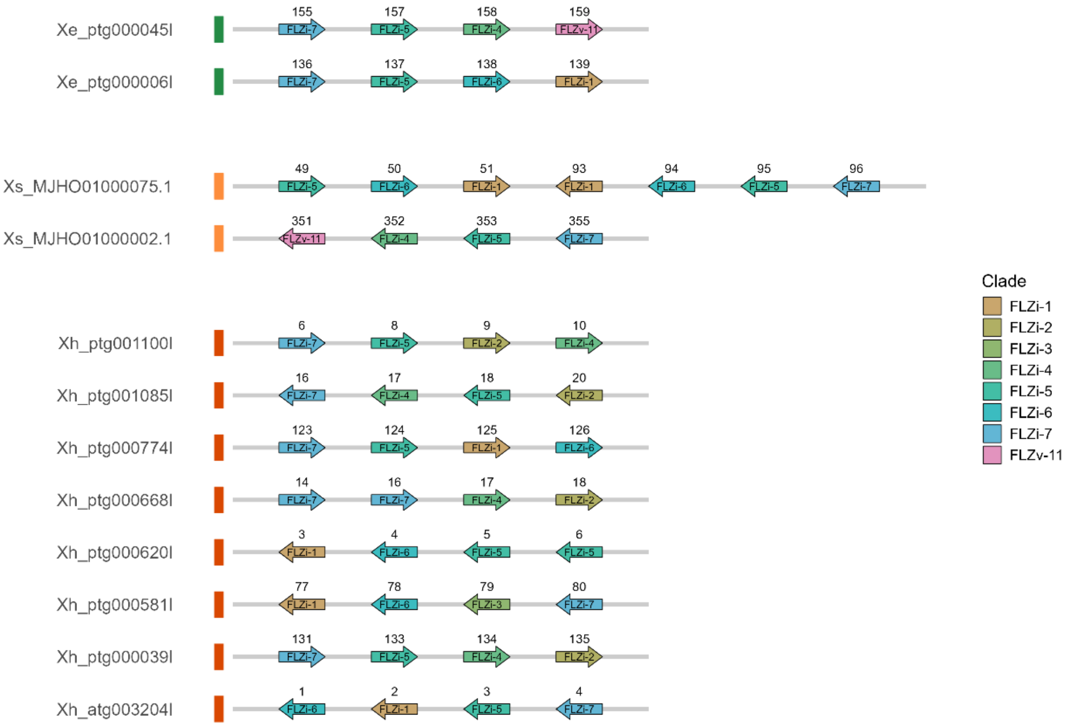


*Figure*

**Figure SN16 *FLZ* Microsynteny Plot**. Illustration of expansion of *FLZ* homologues, as defined by phylogenetic analysis, in *Xerophyta elegans* (*Xe*), *Xerophyta humilis* (*Xh*) and *Xerophyta schlechteri* (*Xs*). Colouring corresponds to clades in PhyML tree for *FLZs*. Arrows pointing to the right represent genes on the forward strand. Arrows pointing to the left represent genes on the reverse strand.

*EXXE*

Two paralogues of this expanded gene family are found in Arabidopsis, Arabidopsis *At1g15010* and *At2g01300*, which are annotated as “Mediator of RNA polymerase II subunit”. However, this is an automatic annotation, and no references are given on how these genes are related to Mediator of RNA polymerase II subunit, nor do they share any sequence similarity. The family is characterised by the presence of the PTHR33782 domain, but no function is known for this domain. We refer to this expanded family as *Expanded in Xerophyta* (*EXXE*), rather than *UNKNOWN*, to distinguish from the many protein families of unknown function in the database. At2g01300 has also been identified as a candidate for interactions with regulatory SnRK1β2 subunit[35]. The SnRK1b2 subunit is important for substrate specificity of the SnRK1 complex.

The *EXXE* family expansion arose independently in *A. bracteata* and *Xerophyta*. Whereas Arabidopsis and rice have two and three *EXXE* genes, respectively, these are the results of recent expansions in these lineages. The expansion of the *EXXE* family genes in the *Xerophyta* lineage is likely to have been the result of the WGT event that preceded their diversification (Fig. SN17). Although several of the *EXXE* genes are located adjacent to each other, this pattern is also seen in the desiccation-sensitive, sister genus *Dioscorea alata* (Fig. SN18-19).

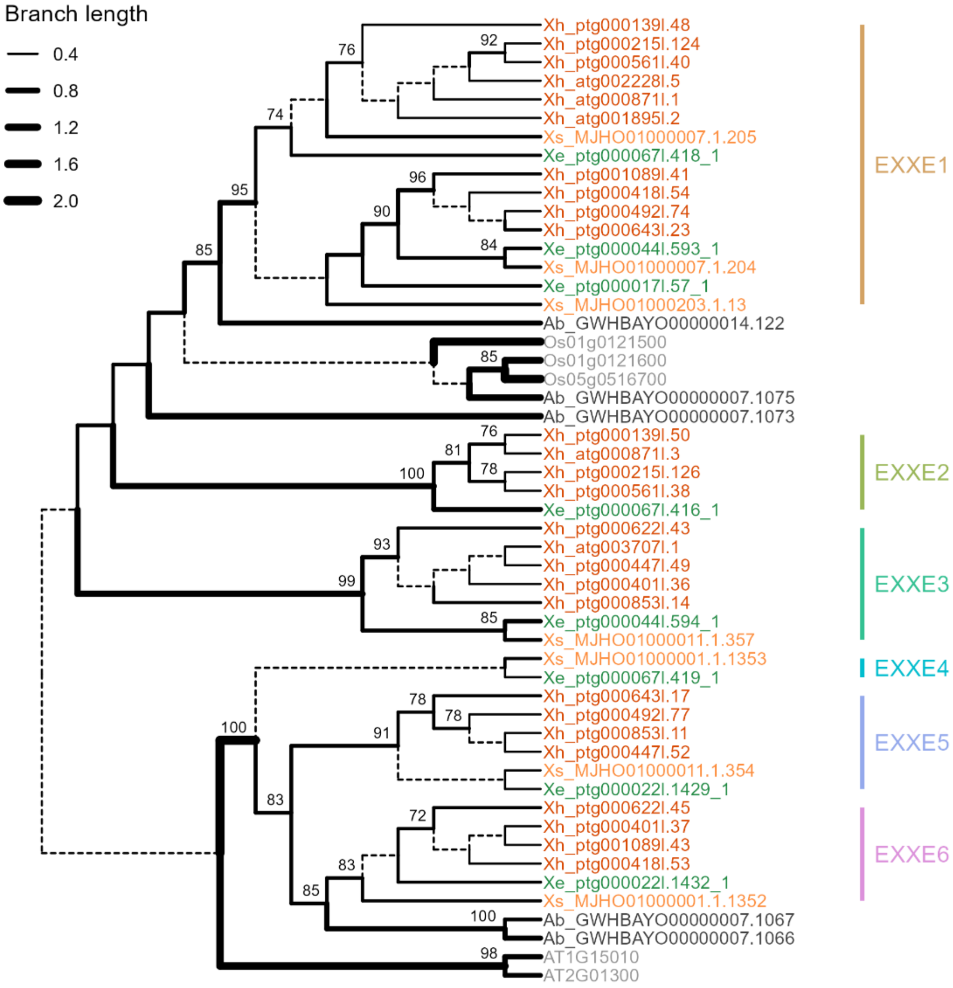


**Figure SN17: *EXXE* phylogeny**. PhyML phylogenetic tree showing relationships between homologues in the expanded *EXXE* gene family in the haploid genome assemblies for *Xerophyta elegans (Xe)*, *Xerophyta schlechteri* (*Xs*) and *Acanthochlamys bracteata (Ab)*, in the tetraploid genome assembly for *Xerophyta humilis* (*Xh*), relative to *Arabidopsis thaliana* and *Oryza sativa*. Branch support >50% is given. *Xerophyta* clade IDs are indicated in colour.

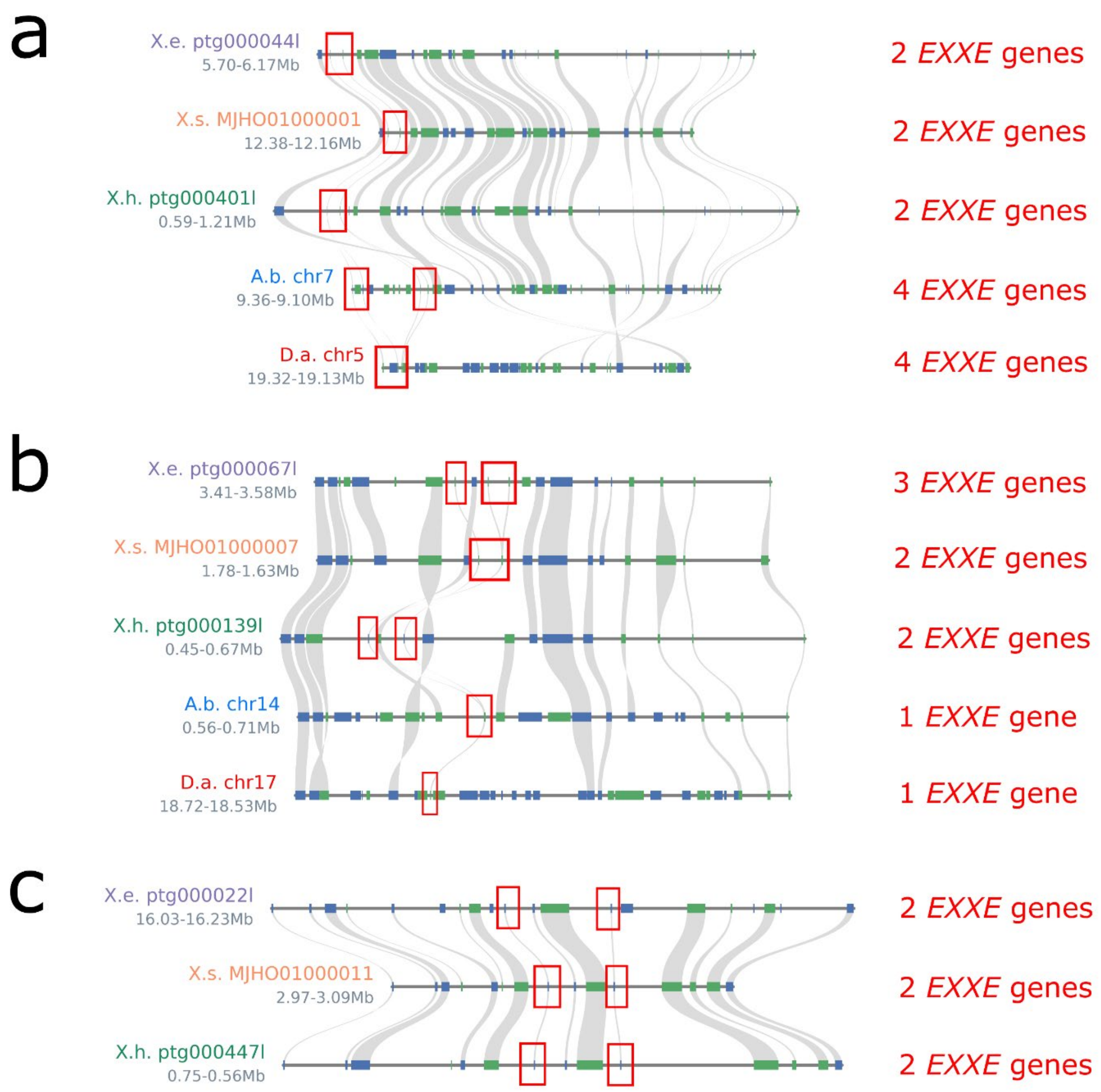


**Figure SN18. EXXE Macrosynteny plot**. Three clusters (a), (b) and (c)of *EXXE* homologues (indicated by red borders) are illustrated for *Xerophyta elegans* (X.e.), *Xerophyta humilis* (X.h.) and *Xerophyta schlechteri* (X.s.) relative to A*canthochlamys bracteata* (A.b.) and *Dioscorea alata* (D. a.). Blue blocks represent genes on the forward strand. Green blocks represent genes on the reverse strand. Text on the left represent contig names. Values on the left represent genomic locations on the contig or chromosome.

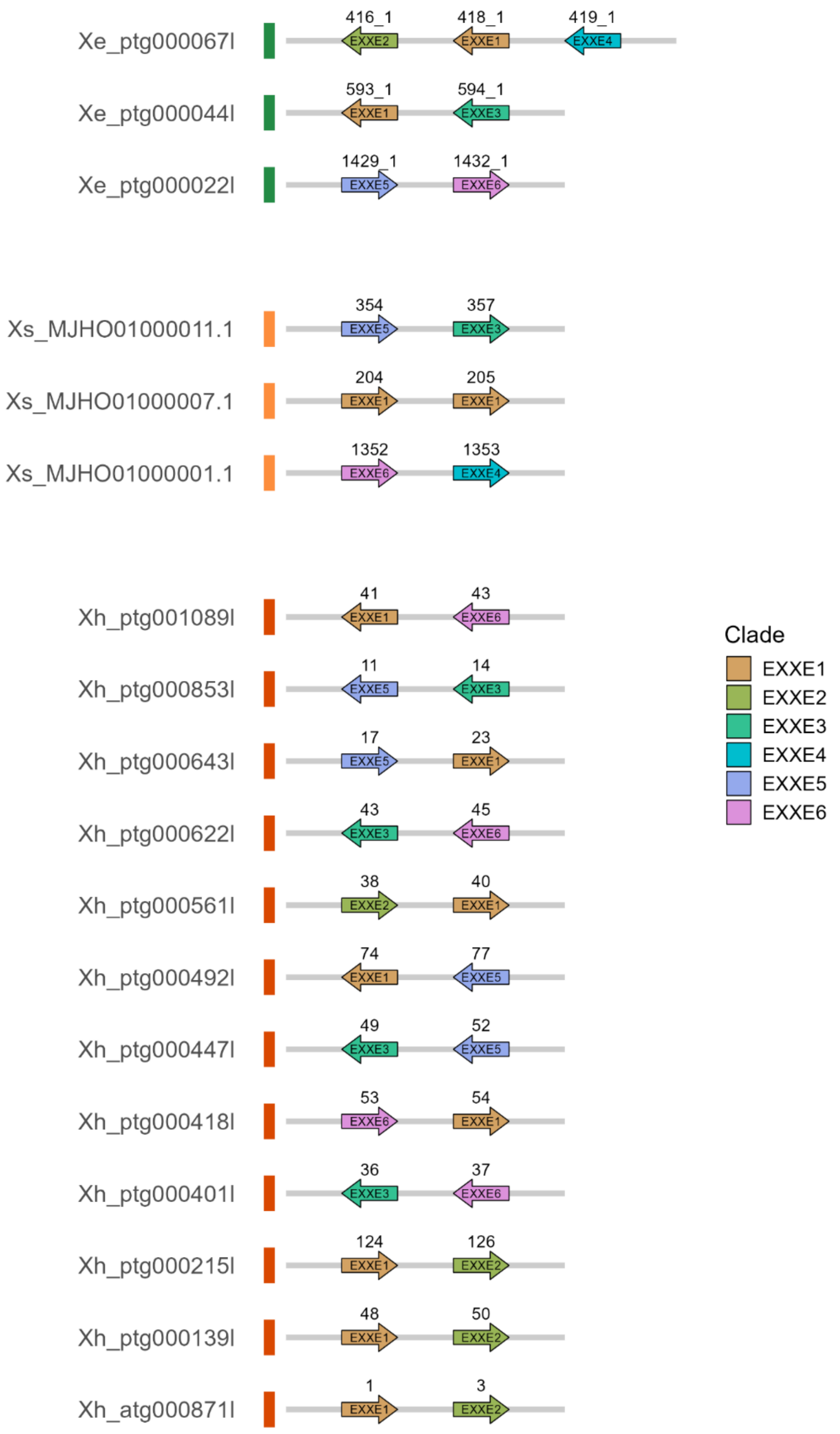


**Figure SN19 *EXXE* Microsynteny plot.** *Illu*stration of expansion of *EXXE* homologues, as defined by phylogenetic analysis, in *Xerophyta elegans* (*Xe*), *Xerophyta humilis* (*Xh*) and *Xerophyta schlechteri* (*Xs*). Colouring corresponds to clades in PhyML tree for *EXXEs*. Arrows pointing to the right represent genes on the forward strand. Arrows pointing to the left represent genes on the reverse strand.

#### 4. ***AT-hook (AHL)*** **and** ***HXXXF-AT*** **expanded gene families**

Members of both the *AT-Hook Motif Nuclear Localized (AHL)* and *HXXXD acyl transferase (HXXXF-AT)* expanded families were down-regulated during dehydration, with the majority of the members of the *AHL* OG upregulated within 2 hours of rehydration.

*AHL*

The AT-Hook is a small DNA binding domain that was first identified in one of the subgroups of the high mobility group (HMG), non -histone chromosomal proteins, namely HMG-1(Y). This group contain a 9 amino acid peptide (called a AT-hook) that binds to the minor groove of DNA, in AT-rich regions. AT-hook containing proteins have been shown to affect chromatin architecture, as well as acting as accessory factors which affect the association of transcription factors with chromatin.

In plants, the AT-hook domain is combined with a conserved Plants and Prokaryotes Conserved (PPC) domain, localized at the carboxy end of the protein. Although this PPC domain is annotated as domain of unknown function (DUF296), it has been shown to be important for nuclear localization of AHL proteins, as well as for protein-protein interactions and the recruitment of transcription factors[37].

Plant AHL proteins can be grouped into three types, Type-I, Type-II and Type-III, based on the combination of AT-hook and PPC domain motifs[37]. Type I AHLs form one clade (Clade A), while the Type-II and Type-III group together in Clade B. The type-I AHLs can further be divided into several sub-families. The genes in the expanded OG0000058 are all Type-I AHL protein. The Type-1 AHL family includes the Arabidopsis Family AHL15-29 genes, which lack introns, and are characterized by one copy of Type I AT hook motif and share the same type of Plant and Prokaryote Conserved (PPC) domain (IPR014476) domain.

It has been proposed that AHL proteins interact with each other, facilitated by the PPC domain, and other nuclear proteins to form macromolecular complexes that modulate plant growth and development[38]. AHL proteins have been shown to interact with NAC transcription factors, and MADS-box genes. In Arabidopsis the clade of AHL15/19/20 proteins are important for promoting the vegetative phase of growth, independently of miR156[39].

*HXXXD acyl-transferase*

This family falls into the BAHD superfamily of acyltransferase enzymes, named after the first four characterized members (B EAT or benzylalcohol O-acetyltransferase; A HCTs or anthocyanin O-hydroxycinnamoyltransferases; H CBT or anthranilate N-hydroxycinnamoyl/benzoyltransferase; D AT or deacetylvindoline 4-O-acetyltransferase)[40]. The BAHD superfamily share two short consensus motifs, HXXXD and DFGWG, but little else. The HXXXD forms the active site for the acyl transfer reaction. Arabidopsis has 55 HXXXD enzymes. These enzymes are involved in the synthesis and elaboration of a wide variety of secondary metabolites.

Phylogenetic analysis of these proteins divided the enzyme family into five distinct clades (I–V), which encode enzymes involved in malonylation of phenolic glucosides (clade I), very long-chain fatty acid (VLCFA) elongation (clade II), volatile ester synthesis (clades III, V), acetylation of amino groups to form amides, and transfer of hydroxycinnamoyl- or benzoyl-CoAs (clade V) [40]. The Arabidopsis members of the expanded orthogroup fall into Clade IIIb [41].

#### Supplementary Note References

11. Soto Gomez, M. *et al.* A bi-organellar phylogenomic study of Pandanales: inference of higher-order relationships and unusual rate-variation patterns. *Cladistics* **36**, 481–504 (2020).
12. Gao, Z.-Y., Li, Z.-H., Lin, D.-L. & Jin, X.-H. Chromosome-Scale Genome Assembly of the Resurrection Plant Acanthochlamys bracteata (Velloziaceae). *Genome Biol Evol* **13**, evab147 (2021).
13. Englbrecht, C. C., Schoof, H. & Böhm, S. Conservation, diversification and expansion of C2H2 zinc finger proteins in the Arabidopsis thaliana genome. *BMC Genomics* **5**, 39 (2004).
14. Ciftci-Yilmaz, S. & Mittler, R. The zinc finger network of plants. *Cell. Mol. Life Sci.* **65**, 1150–1160 (2008).
15. Iida, A., Kazuoka, T., Torikai, S., Kikuchi, H. & Oeda, K. A zinc finger protein RHL41 mediates the light acclimatization response in Arabidopsis. *The Plant Journal* **24**, 191–203 (2000).
16. Rizhsky, L., Davletova, S., Liang, H. & Mittler, R. The Zinc Finger Protein Zat12 Is Required for Cytosolic Ascorbate Peroxidase 1 Expression during Oxidative Stress in Arabidopsis* [boxs]. *Journal of Biological Chemistry* **279**, 11736–11743 (2004).
17. Davletova, S., Schlauch, K., Coutu, J. & Mittler, R. The Zinc-Finger Protein Zat12 Plays a Central Role in Reactive Oxygen and Abiotic Stress Signaling in Arabidopsis. *Plant Physiol* **139**, 847–856 (2005).
18. Hutin, C. *et al.* Early light-induced proteins protect Arabidopsis from photooxidative stress. *Proceedings of the National Academy of Sciences* **100**, 4921–4926 (2003).
19. VanBuren, R., Pardo, J., Man Wai, C., Evans, S. & Bartels, D. Massive Tandem Proliferation of ELIPs Supports Convergent Evolution of Desiccation Tolerance across Land Plants. *Plant Physiol.* **179**, 1040–1049 (2019).
20. Challabathula, D., Zhang, Q. & Bartels, D. Protection of photosynthesis in desiccation-tolerant resurrection plants. *Journal of Plant Physiology* **227**, 84–92 (2018).
21. Mendel, G. *Versuche Über Pflanzen-Hybriden*. (Naturforschender Verein, Brünn, 1866).
