## Supplementary Figures 1-5 for "Transcriptional regulation of the response to water availability in the resurrection plant *Xerophyta elegans*"

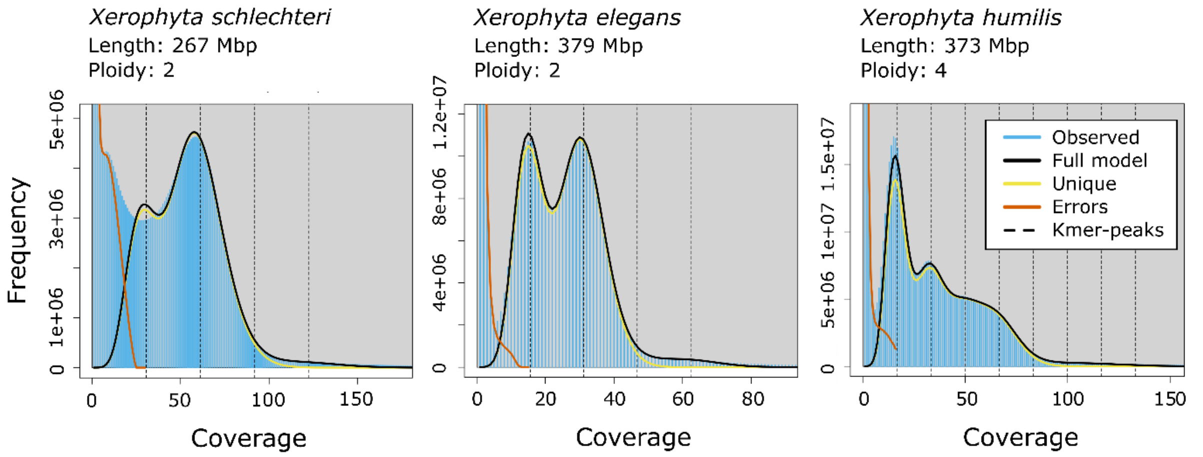


**Supplementary Figure 1**. **K-mer frequency plots of PacBio reads.** The panels display the K-mer frequency plots of the PacBio reads for three species **a.** *Xerophyta schlechteri*, **b**. *Xerophyta elegans*, **c**. *Xerophyta humilis*. The plots were generated using GenomeScope 2.0. The null model was generated based on the assumption of a diploid genome for *X. schlechteri* and *X. elegans*, and tetraploid genome for *X. humilis*.

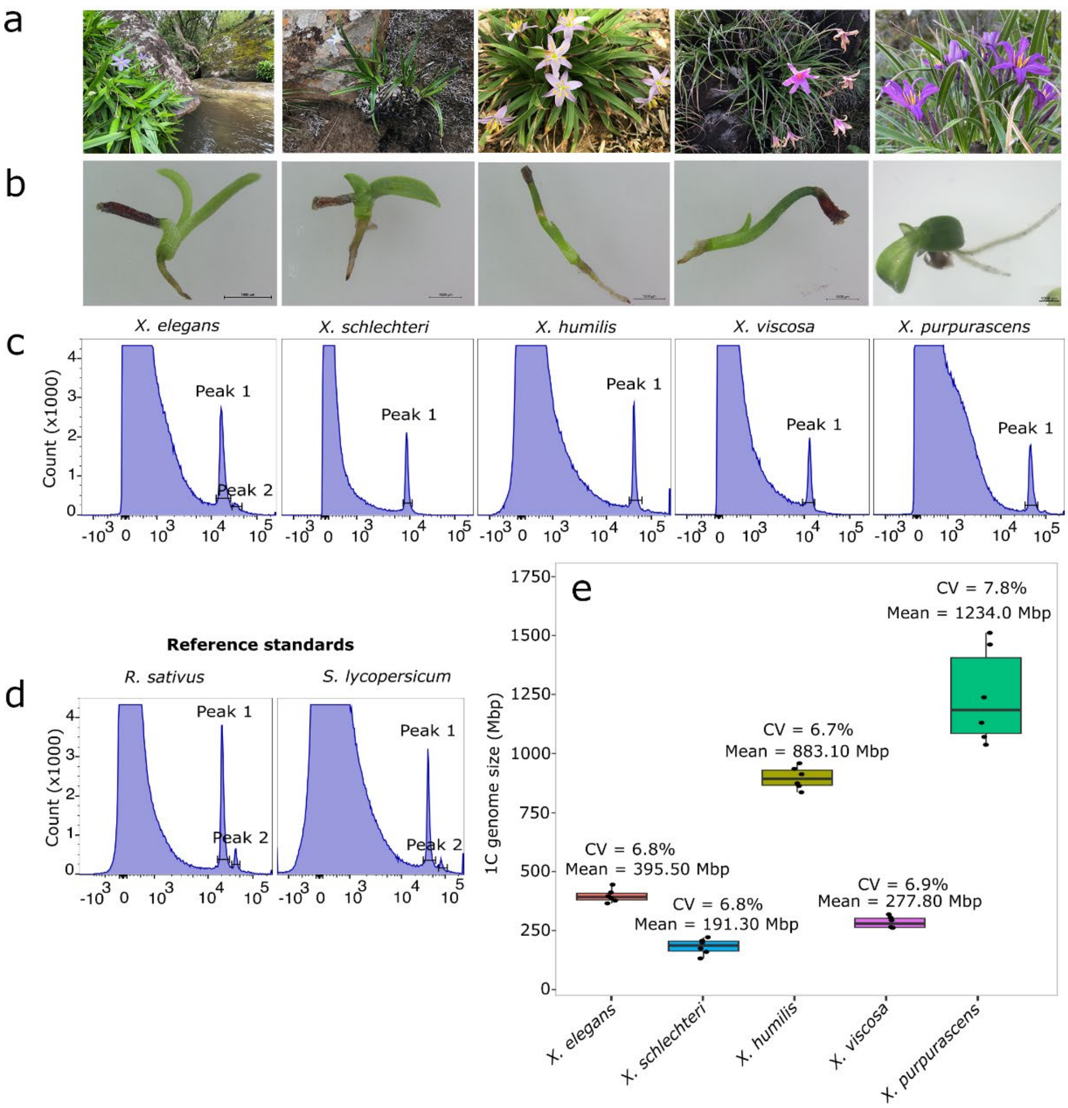


**Supplementary Figure 2. FACS analysis estimates of genome size for five Xerophyta species.** **a.** Photographs of adult *Xerophyta elegans*, *X. schlechteri*, *X. humilis*, *X. viscosa*, and *X. purpurascens* plants. **b.** *X. elegans*, *X. schlectheri* and *X. purpurescens* seedlings have a similar pattern of early seedling development, while *X. humilis* and *X. viscosa* are different. **c.** Histograms of propidium iodide stained nuclei purified from fresh leaf tissue from the five Xerophyta species. **d.** *Raphanus sativus* and *Solanum lycopersicum* were used as standards to estimate the genome sizes of the five *Xerophyta* species. Peaks 1 and 2 in the plots represent G1 and G2 phases of the cell cycle, respectively. **e**. A boxplot summary of means and coefficient of variations (CV) for estimates of genome size (1C) for the five Xerophyta species from six independent biological repeats.

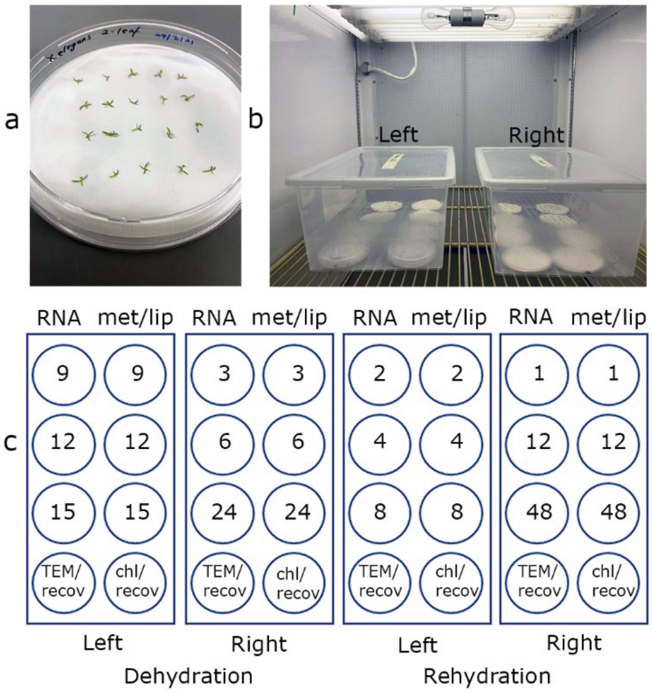


**Supplementary Figure 3. Experimental set-up**. **a**. Twenty-two-leaf-stage *Xerophyta elegans* seedlings were plated out on Whatmann paper wet with 1mL water. **b**. Eight Petri dishes, containing 20 seedlings each, were placed in clear plastic boxes, and dehydrated by the removal of the Petri dish lids. **c.** Layout in the left and right box for matched RNAseq (RNA) and metabolomics/lipid (met/lip) samples. Numbers indicate the hour at which seedlings were sampled in the dehydration and rehydration experiments.

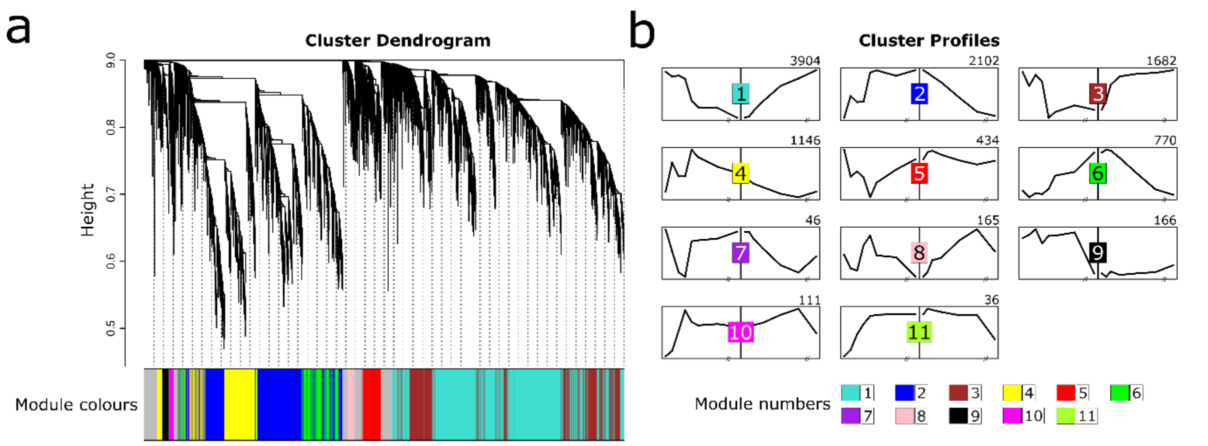


**Supplementary Figure 4. WGCNA clustering of TG DEGs. a**. Dendrogram for the TG DEGs. The resulting 11 clusters are denoted by a representative colour on the bar below the dendrogram. The cluster number which corresponds to these colours is given below. **b**. TG Cluster profiles across the dehydration and rehydration series are given for each cluster. TG Cluster size is indicated to the top right of each profile.

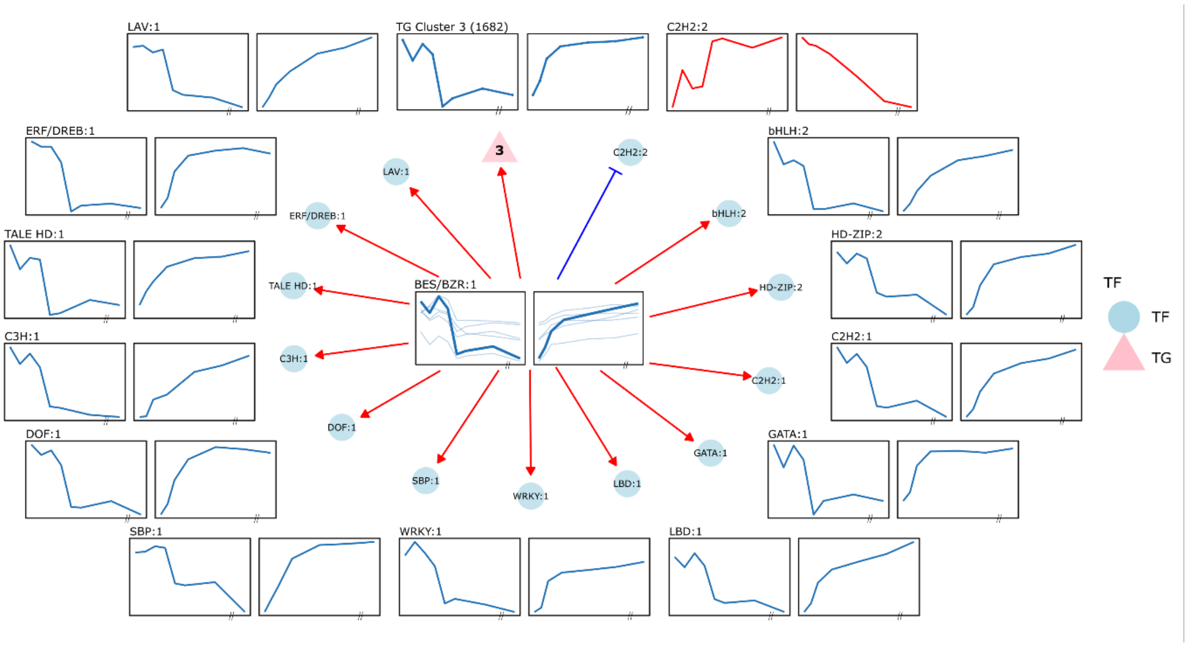


**Supplementary Figure 5: All predicted interactions for the BES/BZR:1 TF cluster over the top 10% of the GRN.** TF clusters are denoted with blue circles with the BES/BZR:1 and GATA:2 TF clusters labelled in bold and TG clusters are denoted with red triangles. Red arrows denote activation and blue flat-head arrows denote repression.
